# Admixture in a butterfly species complex creates a genomic mosaic of ancestry with distinct histories for different chromosomes

**DOI:** 10.1101/2025.11.28.691233

**Authors:** Zachariah Gompert, Lauren K. Lucas, Alia Donley, Matt L. Forister, James A. Fordyce, Chris C. Nice

## Abstract

**Impact Summary:** How well do branching evolutionary trees, in which species form distinct lineages, capture the histories of groups of organisms? Answering this question is central to understanding species, speciation, and the organization of biological diversity. We show that, at least for *Lycaeides* butterflies, the answer differs across the genome. Their genomes contain regions and chromosomes with distinct evolutionary histories, and these differences cannot be explained by the random sorting of ancestral genetic variation alone. Instead, we found substantial evidence of historical hybridization and mixed ancestry across the autosomes, but a distinct and more tree-like history for the Z sex chromosome. Differences among chromosomes were also related to genomic features such as recombination rate. Genetic mapping further suggests that these contrasting evolutionary histories are relevant to wing pattern traits, which influence mate choice in these butterflies. Together, our results reveal that species can contain a mosaic of evolutionary histories and help identify the processes that fuel evolution.

Hybridization and admixture give rise to populations with mosaic genomes composed of ancestry segments with distinct origins and histories. Despite a growing number of documented cases of admixture in nature, relatively little is known about how mosaic patterns of ancestry vary simultaneously across the genome and among multiple populations or lineages of hybridizing species. Here, using pooled whole-genome sequence data from more than 20 populations representing three nominal species and multiple admixed or taxonomically ambiguous populations, we show that widespread admixture in *Lycaeides* butterflies has resulted in distinct evolutionary histories within and among chromosomes, with the most notable differences between the autosomes and the Z sex chromosome. Many populations exhibit substantial evidence of mixed ancestry on the autosomes, whereas the Z chromosome shows a more tree-like evolutionary history. In some cases, the predominant ancestry of the autosomes and the Z chromosome differ. We also find evidence of variation in ancestry within chromosomes, though this appears more idiosyncratic. This variability in evolutionary histories cannot be explained by incomplete lineage sorting alone. Instead, this variability reflects different patterns and outcomes of admixture and introgression across the genome, with these differences explained, in part, by genomic features such as recombination rate and gene density. Finally, using genome-wide association mapping, we demonstrate that wing-pattern traits that are known to influence mate preference are affected by genetic variants on both the autosomes and the Z chromosome, with an elevated influence of the Z. This suggests that combinations of autosomal and Z-chromosome ancestry have the potential to drive wing pattern evolution. In sum, our results show that different chromosomes can exhibit distinct evolutionary histories, with the Z chromosome suggesting more discrete evolutionary lineages or species than the autosomes. These findings have implications both for our basic understanding of species and species boundaries and for how we view the sources of genetic and phenotypic variation that fuel ongoing evolutionary change.

## Introduction

The role of hybridization in adaptation and speciation has been the subject of long-standing debate, but growing evidence supports its common occurrence and (sometimes) creative role in evolution (Mallet, 2005; Taylor & Larson, 2019; Edelman & Mallet, 2021; Peñalba *et al*., 2024). Recent studies have shown that the genomes of many organisms contain DNA segments derived from other species through admixture and introgression, with examples documented in fish (Meier *et al*., 2023; Langdon *et al*., 2024), butterflies (Rosser *et al*., 2024; Boman *et al*., 2025), flies (Suvorov *et al*., 2022), snakes (Nikolakis *et al*., 2022), sunflowers (Hübner *et al*., 2019), conifers (Bolte *et al*., 2024), and humans (Iasi *et al*., 2024; Li *et al*., 2024). We also now know that introgression varies in extent across the genome. For example, introgression is sometimes limited to a few regions that drive adaptation (e.g., Oziolor *et al*., 2019; Rossi *et al*., 2024). In other cases introgression is genomically widespread, excluding only a few regions of the genome that negatively affect hybrid fitness (e.g., Poelstra *et al*., 2014; Schield *et al*., 2024).

The genomic extent of mixed ancestry and introgression is relevant for our understanding of reproductive isolation and the nature of species. Whereas foundational studies of speciation emphasized mechanisms of reproductive isolation and considered it a property of whole organisms or species (e.g., Dobzhansky, 1937; Mayr, 1963; Coyne & Orr, 1989), genomic studies typically measure reproductive isolation as a reduction in effective gene flow and thus regard it as something that varies across the genome (Barton & Bengtsson, 1986; Wu, 2001; Westram *et al*., 2022). Under this genetic framework, genomic regions with limited introgression can function as good species, that is, distinct genetic clusters or independently evolving lineages, even if admixture and introgression are widespread across much of the genome. Consequently, understanding how patterns of introgression and admixture vary across the genome is important for understanding the nature of species and the structure of biological diversity.

Along these lines, patterns of ancestry and introgression often differ between autosomes and sex chromosomes. For example, numerous hybrid zone studies have shown reduced introgression at Z or X loci relative to autosomal loci (Dod *et al*., 1993; Carling & Brumfield, 2008; Storchová *et al*., 2010; Maroja *et al*., 2015; Gompert *et al*., 2017; Hooper *et al*., 2019; Chaturvedi *et al*., 2020; Banker *et al*., 2022; Swaegers *et al*., 2022; Zhang *et al*., 2023). Such patterns, together with findings from interspecific crosses, trait mapping, transcriptomics, population genomics, and molecular evolution, suggest that sex chromosomes play a disproportionate role in speciation and the maintenance of species boundaries (Sperling, 1994; Masly & Presgraves, 2007; Kitano *et al*., 2009; Irwin, 2018; Payseur *et al*., 2018; Sánchez-Ramírez *et al*., 2021). This is consistent with the “large X-effect” and Haldane’s rule, and may arise because of faster sex chromosome evolution or because sex-linked incompatibilities are expressed in the heterogametic sex (Orr, 1997; Carling & Brumfield, 2008; Presgraves, 2008; Lenormand & Roze, 2025). Regardless of the mechanism, and particularly from the perspective of a genetic conception of reproductive isolation, the nature of species and the organization of biodiversity may thus generally differ between sex chromosomes and autosomes, with sex chromosomes behaving more like well-defined species with a bifurcating, tree-like history. The same could be true for other non-recombining regions of the genome, such as some chromosomal rearrangements.

Many studies have examined patterns of admixture and introgression in nature, but much of this work has focused either on introgression between individual species pairs or has not explicitly analyzed how introgression varies among and within chromosomes (but see, e.g., Thawornwattana *et al*., 2023; Langdon *et al*., 2024; van der Heijden *et al*., 2025). Thus, additional studies that quantify patterns of mixed ancestry and introgression across multiple populations and species, and that assess how and why such patterns differ across the genome, are needed to better understand genome-wide variation in evolutionary histories and entities. Moreover, combining such work with trait genetics is important for assessing the contribution of hybridization to fueling phenotypic adaptation (Marques *et al*., 2019).

Here, we use North American butterflies in the genus *Lycaeides* to address this gap (*Lycaeides* has been reclassified as being within the genus *Plebejus* but we retain the older name for consistency with past work and because it more precisely defines our focal lineages; Talavera *et al*., 2013). In North America, *Lycaeides* comprise a complex of several nominal species (i.e., *L. anna*, *L. idas*, *L. melissa*, and *L. samuelis*) as well as admixed or taxonomically ambiguous lineages (e.g., the Sierra Nevada, Warner Mountain, and Rocky Mountain populations described below) (Nabokov, 1949; Gompert *et al*., 2006, 2010; Forister *et al*., 2011; Gompert *et al*., 2012; Nice *et al*., 2013; Gompert *et al*., 2014). At least one contemporary hybrid zone also exists (Chaturvedi *et al*., 2020; Zhang *et al*., 2023). Despite past and ongoing hybridization, *Lycaeides* species differ in several traits, including male genitalic morphology and wing patterns, which are primarily used to delineate species, as well as host plant use and preference, voltinism and phenology, and mate preference (Nice & Shapiro, 1999; Fordyce *et al*., 2002; Nice *et al*., 2002; Forister *et al*., 2006; Lucas *et al*., 2008; Forister *et al*., 2009; Gompert *et al*., 2013b,a; Lucas *et al*., 2018). Putative admixed populations display unique combinations of parental, intermediate and transgressive traits and ecologies. For example, putative admixed lineages in the Sierra Nevada and White Mountains (hereafter collectively referred to as Sierra Nevada populations) occupy alpine habitats, feed on an alpine endemic host plant, *Astragalus whitneyi*, and have a unique, locally adaptive lack of egg adhesion (Gompert *et al*., 2006; Nice *et al*., 2013). Suspected admixed populations in the Warner Mountains also use an *Astragalus* host and exhibit a mosaic of *L. anna*, *L. melissa*, and intermediate morphological traits (Nice *et al*., 2013). Admixed populations are also known from the Rocky Mountains (specifically in and around Jackson Hole, hereafter referred to as Rocky Mountain populations); these mostly use the same host and occupy similar habitats (montane forest meadows and sagebrush-dominated buttes) as nearby *L. idas*, but have *L. melissa*-like genitalic morphology and show evidence of mixed *L. melissa* and *L. idas* ancestry (Gompert *et al*., 2010, 2012, 2013b). Relevant to our genomic analyses, *Lycaeides*, like Lepidoptera generally, have female heterogamety (females are ZW and males are ZZ) and achiasmatic female meiosis (i.e., crossing over does not occur in females) (Rasmussen, 1977; López & Mercier, 2024). Our current *L. melissa* genome assembly comprises 22 autosomes and the Z sex chromosome, ranging from 10.3 to 33.3 megabase pairs (Mb) in length; the Z chromosome is 27.4 Mb (Zhang *et al*., 2023).

Past population genomic studies of *Lycaeides* suggest that genetic differentiation and introgression vary across the genome, but these inferences have been based either on limited genomic sampling (e.g., genotyping-by-sequencing [GBS] data), limited spatial sampling (e.g., a single hybrid zone), or both (Gompert *et al*., 2012; Chaturvedi *et al*., 2020; Zhang *et al*., 2023). Moreover, previous work has largely lacked *Lycaeides* samples from Alaska (where colonization of North America occurred) and outgroup taxa from Eurasia (where the closest extant relatives of the North American taxa occur) (Vila *et al*., 2011; Toro-Delgado *et al*., 2025), which together have limited our ability to infer the direction of evolutionary change. In the current study, we overcome these limitations by using pooled whole-genome sequencing of 23 *Lycaeides* populations–comprising the nominal species *L. anna*, *L. idas* (including Alaskan individuals), and *L. melissa*, as well as several named and unnamed putative hybrid taxa–combined with population genomic and phylogenetic methods to quantify patterns of ancestry and admixture throughout this genus and within and among chromosomes.

We address three main questions: (i) how much do patterns of ancestry vary across the genome, (ii) does the Z chromosome exhibit a more tree-like evolutionary history than the autosomes, and (iii) to what extent can variation in ancestry within and among chromosomes be explained by chromosomal features, such as recombination rate, or the relative contributions of incomplete lineage sorting (ILS) and introgression? Our central hypothesis is that the Z chromosome disproportionately influences ecologically important traits and hybrid fitness and therefore has retained a more tree-like evolutionary history despite widespread hybridization and introgression across much of the genome. Alternatively, admixture may be either limited throughout the genome or sufficiently pervasive that the Z chromosome does not differ appreciably from the autosomes. To further evaluate the biological significance of these predictions, we integrate genome-wide association mapping of previously published wing pattern data with our population genomic analyses. This allows us to test two related questions: whether the inferred patterns of admixture are likely to have phenotypic consequences and whether the Z chromosome contributes disproportionately to variation in wing pattern, a trait that moderately influences mate choice in *Lycaeides* butterflies (Fordyce *et al*., 2002; Gompert *et al*., 2013b).

## Methods

### Genetic data

We obtained DNA from 996 butterflies representing 23 populations (678 newly extracted and 318 from previous studies, Gompert *et al*., 2014, 2021; Table 1 and Figure 1A). These included 142 *L. anna* from three populations, 148 *L. idas* from four populations, 339 *L. melissa* from seven populations, 357 unnamed, admixed, or taxonomically ambiguous *Lycaeides* from eight populations, and 10 *L. argyrognomon* from a single population (Menee, France, population code MEN) as an outgroup. We extracted the 678 new DNA samples using Qiagen’s DNeasy 96 Blood and Tissue Kit. We measured DNA concentrations with a fluorescence microplate reader (Turner Biosystems Modulus Microplate) and combined equal total amounts of DNA (by mass) from butterflies for each population. These population pools of DNA were used for pooled whole-genome sequencing. Sequencing libraries were constructed and sequenced on the DNBseq platform (paired-end 100 bp reads) by BGI. We repeated the pooling, library preparation and sequencing twice for four populations for quality control. Raw sequences were trimmed and filtered with SOAPnuke (Chen *et al*., 2018). See “Library preparation, sequencing and read filtering” in the Online Supplemental Material (OSM) for details.

**Figure 1:**
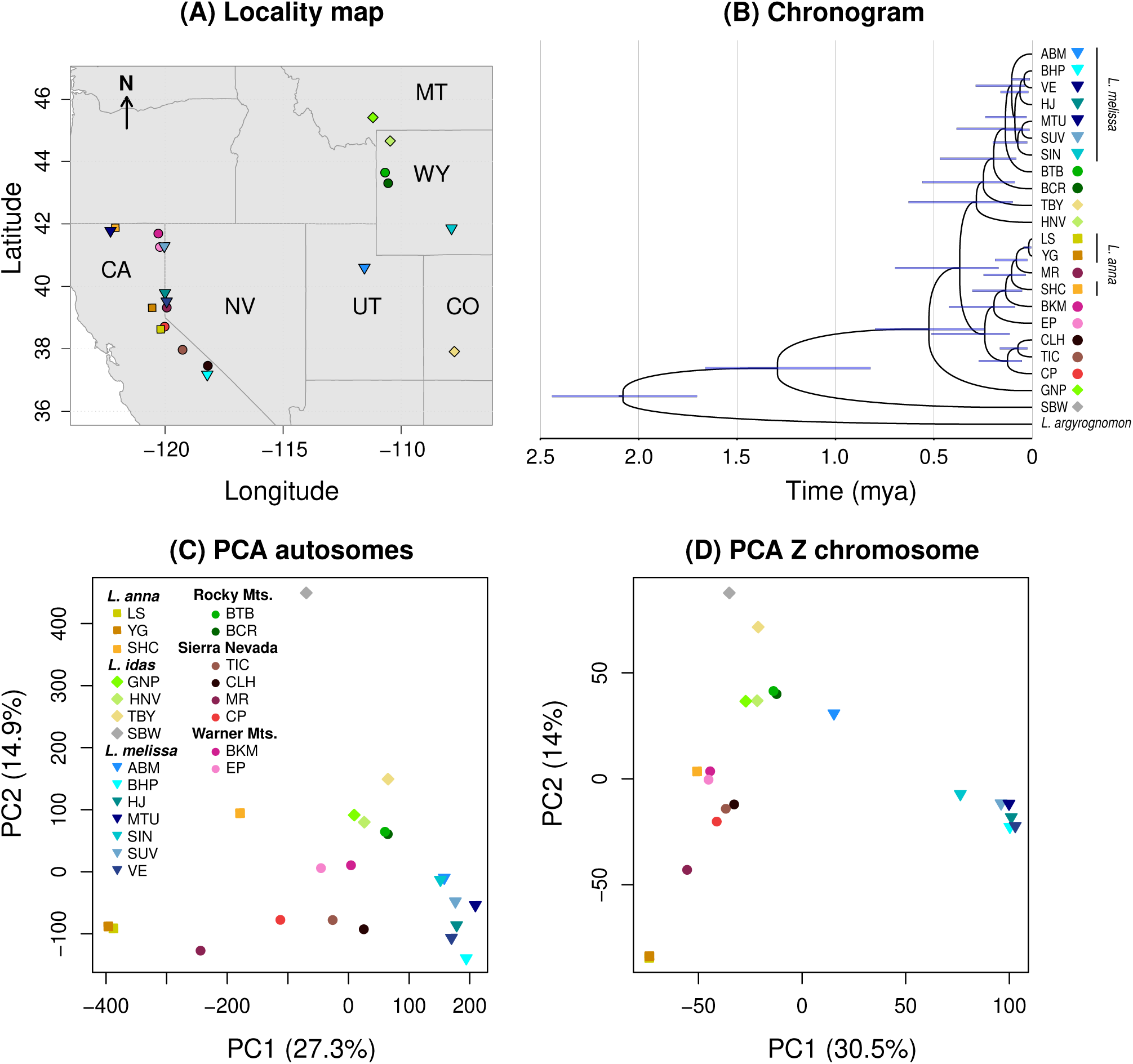
Sample localities and summary of phylogenetic relationships and genetic variation. (A) The map shows the locations of the butterfly populations in the contiguous, western USA (the *L. idas* population from Alaska, SBW, and the outgroup, *L. argyrognomon*, are not shown). State abbreviations are given. (B) A chronogram depicts our Bayesian reconstruction of the (consensus) best-tree approximation to the evolutionary history of *Lycaeides*. Time estimates are in millions of years ago (mya) and shaded bars denote 95% highest-posterior densities for node ages. Statistical summaries of patterns of population genetic structure for the sampled *Lycaeides* based on principal component analyses of allele frequencies for the autosomes (C) or Z chromosome (D). Colored symbols denote populations, which are organized in the inset legend by species or geographic region for taxonomically ambiguous populations.

**Table 1:**
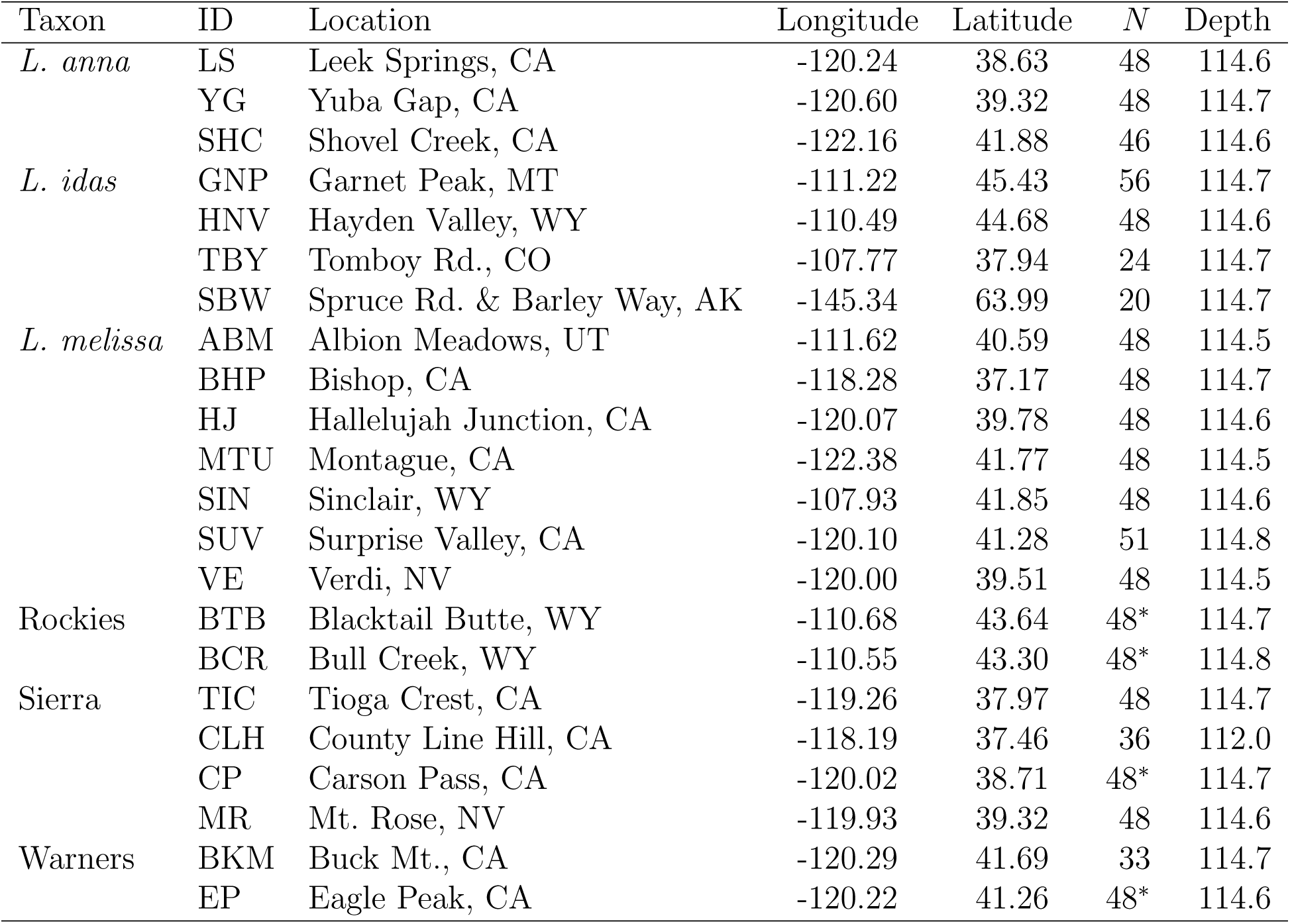
Population IDs, locations, geographic coordinates (in degrees) and sample sizes (*N*, in terms of number of diploid individuals), and read depth (i.e., depth or coverage) for the 22 *Lycaeides* populations (i.e., excluding *L. argyrognomon*, N = 10, depth = 114.7). Asterisks denote samples that were pooled and sequenced twice. The “Rockies” designation refers to Jackson Hole *Lycaeides* from past work (Gompert *et al*., 2012); “Sierra” includes the Sierra Nevada and CLH, which is from the nearby White Mountains.

| Taxon | ID | Location | Longitude | Latitude | $N$ | Depth |
| --- | --- | --- | --- | --- | --- | --- |
| *L. anna* | LS | Leek Springs, CA | -120.24 | 38.63 | 48 | 114.6 |
|  | YG | Yuba Gap, CA | -120.60 | 39.32 | 48 | 114.7 |
|  | SHC | Shovel Creek, CA | -122.16 | 41.88 | 46 | 114.6 |
| *L. idas* | GNP | Garnet Peak, MT | -111.22 | 45.43 | 56 | 114.7 |
|  | HNV | Hayden Valley, WY | -110.49 | 44.68 | 48 | 114.6 |
|  | TBY | Tomboy Rd., CO | -107.77 | 37.94 | 24 | 114.7 |
|  | SBW | Spruce Rd. & Barley Way, AK | -145.34 | 63.99 | 20 | 114.7 |
| *L. melissa* | ABM | Albion Meadows, UT | -111.62 | 40.59 | 48 | 114.5 |
|  | BHP | Bishop, CA | -118.28 | 37.17 | 48 | 114.7 |
|  | HJ | Hallelujah Junction, CA | -120.07 | 39.78 | 48 | 114.6 |
|  | MTU | Montague, CA | -122.38 | 41.77 | 48 | 114.5 |
|  | SIN | Sinclair, WY | -107.93 | 41.85 | 48 | 114.6 |
|  | SUV | Surprise Valley, CA | -120.10 | 41.28 | 51 | 114.8 |
|  | VE | Verdi, NV | -120.00 | 39.51 | 48 | 114.5 |
| Rockies | BTB | Blacktail Butte, WY | -110.68 | 43.64 | 48* | 114.7 |
|  | BCR | Bull Creek, WY | -110.55 | 43.30 | 48* | 114.8 |
| Sierra | TIC | Tioga Crest, CA | -119.26 | 37.97 | 48 | 114.7 |
|  | CLH | County Line Hill, CA | -118.19 | 37.46 | 36 | 112.0 |
|  | CP | Carson Pass, CA | -120.02 | 38.71 | 48* | 114.7 |
|  | MR | Mt. Rose, NV | -119.93 | 39.32 | 48 | 114.6 |
| Warners | BKM | Buck Mt., CA | -120.29 | 41.69 | 33 | 114.7 |
|  | EP | Eagle Peak, CA | -120.22 | 41.26 | 48* | 114.6 |

We aligned the sequence reads to the *L. melissa* genome (Zhang *et al*., 2023) using bwa-mem2 (version 2.0pre2) (Li & Durbin, 2009; Vasimuddin *et al*., 2019). PCR duplicates were then marked and removed with samtools (version 1.16) (Li *et al*., 2009; Ebbert *et al*., 2016). We called genetic variants using bcftools (version 1.16) (Li, 2011) and filtered the resulting initial SNP set with GATK (version 4.1.4.1) (McKenna *et al*., 2010), resulting in 15,914,826 SNPs. We extracted the counts of high-quality reads supporting the reference and alternate allele for each SNP and population; we then used these counts to estimate allele frequencies and conduct downstream population genomic and phylogenetic analyses. See “Alignment, variant calling, filtering and allele frequency estimation” in the OSM for additional details.

### Phylogeny and population structure

We began by characterizing the overall relationships among the *Lycaeides* populations. We took two complementary approaches. First, we used BEAST (version 2.7.7) to estimate a time-calibrated phylogeny (a chronogram) from a subset of genome-wide SNPs (Drummond & Rambaut, 2007; Bouckaert *et al*., 2019). Specifically, we used a concatenated set of 5408 SNPs interspersed across the genome (as required for computational feasibility); variable sites within populations were coded as the more common allele when its frequency was ≥ 0.95 and as ambiguous (N) when this was not true. Invariant sites were also accounted for in the model. We used existing estimates for the divergence of *L. argyrognomon* and North American *Lycaeides* and for the colonization of North America by *Lycaeides* to construct informative priors for the molecular clock in our analysis (Vila *et al*., 2011; Kawahara *et al*., 2023). See “Data processing, models and priors for BEAST” in the OSM for additional details. Second, we summarized patterns of genetic variation and genetic similarity using a principal components analysis (PCA) of genome-wide allele frequencies. We did this in R. We separately analyzed the 14,907,399 autosomal SNPs and the 1,007,427 Z chromosome SNPs. In both cases, we included only the *Lycaeides* populations (i.e., we excluded the outgroup, *L. argyrognomon*) and analyzed the centered, but not standardized, allele frequency matrix.

### Tests for tree-like histories

We next asked how well the evolutionary relationships among the *Lycaeides* populations could be explained by a bifurcating population tree for each of the 22 autosomes and the Z chromosome. To this end, we used TreeMix (version 1.13) to estimate a composite maximum-likelihood population tree for each chromosome from the allele-frequency covariance matrix (residuals from the tree model are shown in Figure S1). We rooted each tree using *L. argyrognomon* and used a block-resampling approach with blocks of 100 SNPs to estimate the unobserved population allele-frequency covariance matrix from the observed sample covariance matrix. We next sequentially added migration (admixture) edges to each maximum-likelihood tree to generate admixture graphs (note that admixture graphs efficiently summarize population genetic variation but, nonetheless, suffer from limited identifiability and should not be interpreted as unique reconstructions of demographic history; Lipson, 2020). Because covariance explained necessarily increases as migration edges are added, we use this trajectory descriptively rather than as a formal model-selection criterion. This procedure was repeated 20 times for each chromosome and each number of migration edges to increase the likelihood of identifying the optimal solution. We then calculated the proportion of allele-frequency covariance explained by each graph, from zero (a bifurcating tree) to eight migration edges. A higher proportion of covariance explained with fewer migration edges indicates a more tree-like evolutionary history. We examined the nature and weights (proxies for admixture proportions) of the migration edges across chromosomes and graphs to identify the most noteworthy instances of putative admixture.

We also analyzed the residual covariance matrices from the population trees (i.e., graphs without migration edges) to better characterize departures from a strictly bifurcating history. Specifically, we asked whether residual covariance was concentrated among a small number of population pairs or distributed more broadly across the tree, and whether these patterns were consistent across chromosomes (see “Residual structure for TreeMix” in the OSM).

We then tested whether chromosome-level features predicted the proportion of allele-frequency covariance explained by population graphs with different numbers of migration edges. We focused on the 22 autosomes and on features associated with recombination (chromosome size and direct estimates of population recombination rate) and selection (gene density). This analysis was motivated by studies in other systems reporting a negative relationship between per-base-pair recombination rate (often inferred from chromosome size) and introgression of minor-parent ancestry (e.g., Brandvain *et al*., 2014; Schumer *et al*., 2018). Unlike those studies, however, we do not distinguish major and minor parental lineages and instead ask whether chromosome-level features predict the fit of population trees and admixture graphs, rather than introgression itself. Consequently, expectations may differ for bifurcating trees, which exclude introgression, and admixture graphs, which explicitly model migration. For example, if selection acts against foreign ancestry, longer chromosomes with lower recombination rates (per base-pair) and higher gene densities might retain less introgressed ancestry and therefore be better explained by bifurcating trees (akin to, Brandvain *et al*., 2014; Schumer *et al*., 2018), whereas this relationship might weaken or disappear for admixture graphs that explicitly model migration. Gene density and chromosome size were calculated from the *L. melissa* genome assembly and annotations (Zhang *et al*., 2023; Goodwin *et al*., 2026), whereas population recombination rates (*ρ* = 4*N_e_r*) were estimated for five lineages from moderate-coverage whole-genome sequence data using iSMC (Barroso *et al*., 2019) (see “Estimating recombination-rate landscapes”). We then fit Bayesian regression models to test for associations between these chromosome-level features and the proportion of allele-frequency covariance explained by population trees and admixture graphs with different numbers of migration edges (see “Bayesian regression methods” in the OSM for details).

### Genomic variation in evolutionary relationships

We used the program CASTER (version 1.20.2.5) to quantify variation in tree topologies within and among chromosomes and thereby further assess genome-wide variability in evolutionary histories (Zhang *et al*., 2025). CASTER implements a site-based, coalescent-aware species-tree estimation method for rapid, genome-scale phylogenetic analyses. Importantly for our purposes, the approach produces a score for each SNP that can be used to detect changes in evolutionary histories across the genome.

We first used CASTER to estimate trees for each chromosome. We used the CASTER-site model for this, set *L. argyrognomon* as the outgroup, and coded SNPs within populations with the common allele when its frequency exceeded 0.95 and otherwise as ambiguous (N). We used the generalized Robinson-Foulds distance to quantify differences in tree topologies among chromosomes; this metric captures differences in mutual clustering information (in bits) among tree topologies (Smith, 2020a). We did this using the TreeDist R package (version 2.11.0) (Smith, 2020b). We then asked whether the tree distances among chromosomes were greater than expected under a null model where all chromosomes shared the same species tree (i.e., under a null model where SNPs were permuted among chromosomes). SNPs were repeatedly randomized among chromosomes (*n* = 50 permutations) to generate null expectations.

We next used CASTER to quantify variability in evolutionary histories both within and among chromosomes. To do this, we computed scores for alternative four-taxon trees in sliding windows along chromosomes (similar to earlier methods for topology weighting, e.g., Martin & Van Belleghem, 2017). We focused on four-taxon sets representing putative cases of admixture suggested by our TreeMix results, differences in tree topologies among chromosomes from CASTER, and results from previous population genomic studies of *Lycaeides* (Gompert *et al*., 2006, 2010; Nice *et al*., 2013; Gompert *et al*., 2014), as well as control four-taxon sets where admixture was not expected. In all cases, the Alaskan *L. idas* population (SBW) was included as an outgroup for the other three populations analyzed. We computed scores in 10-kilobase windows using CASTER-slidingwindow (https://github.com/chaoszhang/ASTER) and then averaged and normalized these scores across 50 kilobase sliding windows (with 10-kilobase steps) in R. The normalized scores sum to one over the possible topologies. Substantial variation in scores within and among chromosomes, especially over large genomic regions, is strongly suggestive of admixture, with the scores representing the relative support for, or contribution of, different histories (Zhang *et al*., 2025).

We summarized and analyzed the window-based topology scores to test whether variation in local genealogies could be explained by incomplete lineage sorting (ILS) alone. Our goal was not to infer ILS for individual windows, but rather to determine whether ILS could account for differences in ancestry patterns between the autosomes and the Z chromosome or whether these differences instead required additional processes, primarily gene flow or introgression. For these analyses, we used the scores described above but without normalization. To this end, we focused on two summaries of the topology scores: (i) the summed unnormalized topology score (Σ), which quantifies phylogenetic signal (consistent support for a single topology) (following Zhang *et al*., 2025), and (ii) asymmetries in the scores for the alternative (non-species-tree) topologies, which are not expected under ILS (Degnan & Rosenberg, 2009; Zhang *et al*., 2025). We used permutation tests to assess whether phylogenetic signal was greater on the Z chromosome than on the autosomes (Σ*_Z_ >* Σ*_A_*) and whether asymmetry in the alternative topologies differed between them (Δ*_Z_* ≠ Δ*_A_*). We also computed bootstrap confidence intervals for the asymmetry statistic (Δ) for the Z chromosome, autosomes, and the genome as a whole. Under ILS, Δ ≈ 0; thus, confidence intervals overlapping zero are consistent with ILS (Degnan & Rosenberg, 2009; Zhang *et al*., 2025).

To evaluate the broader determinants of topology variation, we also fit regression models to test whether variation in Σ and Δ was associated with recombination-rate variation within and among chromosomes (see “Tests of incomplete lineage sorting and its alternatives” in the OSM).

### Genetic analyses of wing pattern

Finally, we asked whether the chromosome-specific evolutionary histories inferred above have phenotypic consequences. We used wing pattern as a test case and specifically asked whether (i) the Z chromosome harbors a greater density of wing pattern-associated SNPs than the autosomes and (ii) both the autosomes and the Z chromosome make substantial contributions to wing pattern variation. Support for the latter prediction would suggest that admixture among lineages can generate novel combinations of wing pattern alleles through the reassortment of autosomal and Z-linked variants. *Lycaeides* wing patterns comprise a series of black spots and aurorae (border ocelli containing black and orange pigments, together with structural coloration) (Nabokov, 1943; Fordyce *et al*., 2002; Lucas *et al*., 2018). We reanalyzed wing pattern measurements (the sizes of 17 wing pattern elements) and corresponding genetic data from three *Lycaeides* populations: YG (*L. anna*, *n* = 100), GNP (*L. idas*, *n* = 98), and SIN (*L. melissa*, *n* = 97) (Lucas *et al*., 2018) (Figure S2). We used Bayesian sparse linear mixed models implemented in gemma (version 0.95a) (Zhou & Stephens, 2012) to estimate, for each trait and population, the proportion of additive genetic variance explained by each chromosome using a polygenic genome-wide association framework (Zhou *et al*., 2013). For each trait and population, we ran 20 MCMC chains of 1,000,000 iterations with 500,000-iteration burn-ins and estimated additive genetic variances from the model-averaged SNP effect estimates and population allele frequencies. See “Wing pattern analyses” in the OSM for additional details.

## Results

### Phylogeny and population structure

Bayesian phylogenetic analysis suggests that the sampled North American *Lycaeides* diverged from a common ancestor approximately 1.3 million years ago (mya) (95% highest posterior interval [HPI] = 0.8-1.7), with the most basal split separating the Alaskan *L. idas* population (SBW) from the other populations, which occupy the contiguous United States and diverged from each other around 0.5 mya (95% HPI = 0.2-0.8) (Figure 1A,B). Our results indicate that *L. idas* is paraphyletic, even when considering only populations from the contiguous United States, whereas *L. melissa* and *L. anna* are monophyletic, at least if putative admixed populations (specifically MR) are excluded. The most recent common ancestors of *L. melissa* and *L. anna* were estimated to have existed 130 thousand (95% HPI = 29-238) and 120 thousand (95% HPI = 33-247) years ago, respectively. The taxonomically ambiguous and putative hybrid lineages from the Rocky Mountains, Sierra Nevada, and Warner Mountains generally formed paraphyletic clades nested within *L. idas*, but not within *L. melissa* or *L. anna* (with the exception of MR). Results from PCA were largely consistent with the phylogenetic tree but also highlighted differences in population genetic structure between the autosomes and the Z chromosome (Figure 1C,D). In both cases, PC1 separated *L. anna* and *L. melissa*, with *L. idas* and taxonomically ambiguous populations showing intermediate PC1 scores; however, these populations tended to be more similar to *L. anna* (i.e., lower PC1 scores) for the Z chromosome than for the autosomes. Interestingly, one *L. melissa* population (ABM) exhibited a more *L. anna*-like PC1 score for the Z chromosome. PC2 separated the Alaskan *L. idas* population (SBW) from the other *Lycaeides* populations, with a notable gap in PC space for the autosomes but not for the Z chromosome, which instead showed a more continuous distribution along PC2.

### Tests for tree-like histories

A strictly bifurcating tree (i.e., a population graph with 0 migration edges) explained allele-frequency covariance better for the Z chromosome (96.0%) than for the autosomes (mean = 87.1%, range = 84.2–89.6%) (Figure 2A). Adding migration edges increased the variance explained for all chromosomes, particularly the autosomes, such that graphs with eight migration edges explained 99.5% of the covariance for the Z chromosome and an average of 98.1% for the autosomes (range = 97.2–98.7%). Residuals from the bifurcating trees (*m* = 0) were generally similar across chromosomes, suggesting that the same population relationships deviated from the tree across much of the genome (Figure S1). However, this similarity in residual structure was weaker between the Z chromosome and the autosomes than among the autosomes themselves (Figure S3). Consistent with this distinction, although the Z chromosome had less residual variation overall, that variation was distributed across more effective dimensions than for the autosomes, where larger residuals were concentrated in fewer dominant patterns (Figure 2B).

**Figure 2:**
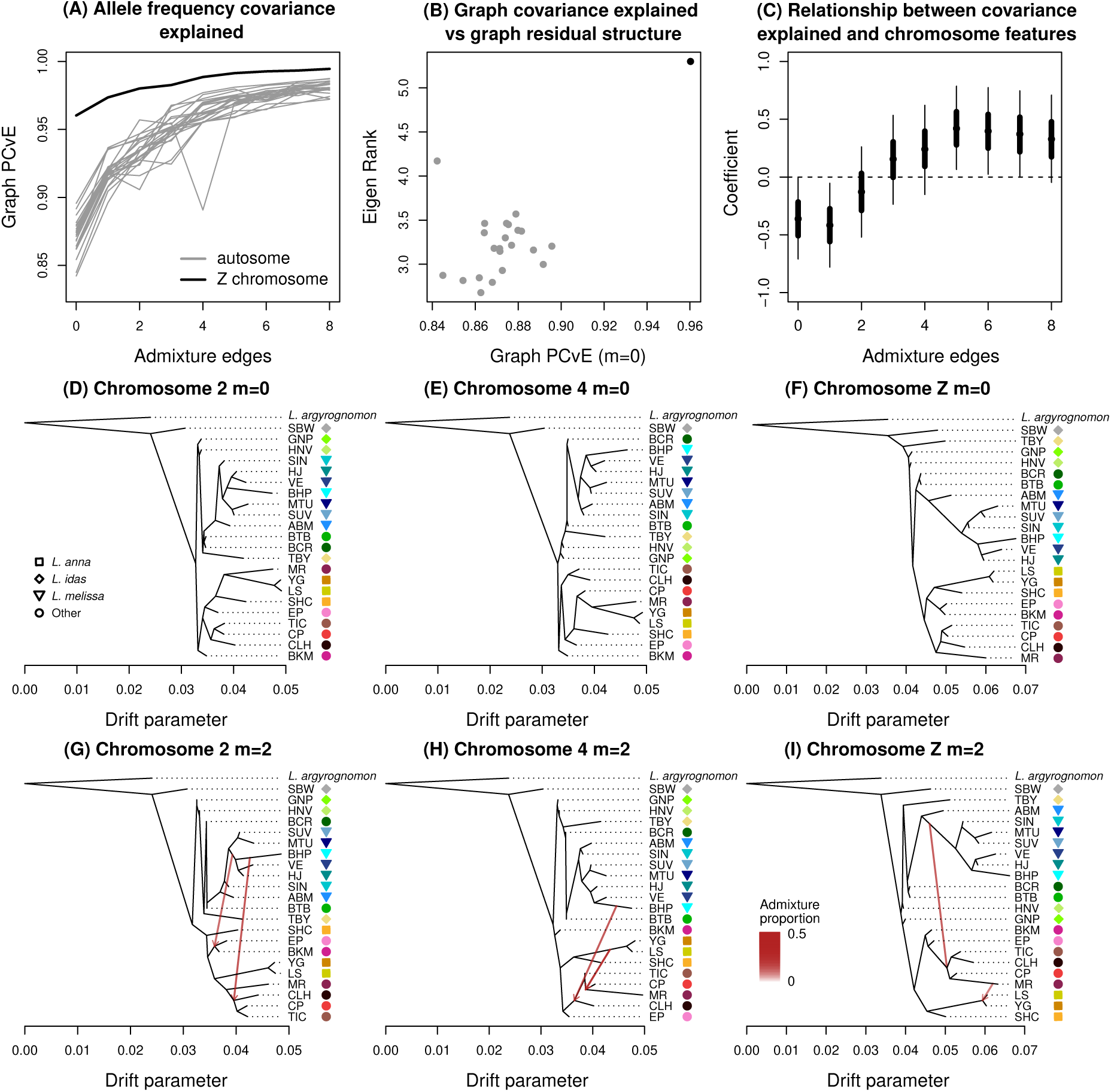
Patterns of genetic variation explained by population trees and admixture graphs from TreeMix. Panel (A) shows the proportion of allele frequency covariance explained (PCvE) by graphs with 0 to 8 admixture (i.e., migration) edges (a graph with 0 edges is a bifurcating tree). Gray lines correspond to the 22 autosomes; the black line denotes the Z chromosome. Panel (B) depicts the PCvE for a graph with no admixture edges (*m* = 0) versus the effective rank from a principal components analysis (PCA) of the residuals from the graph for each autosome (gray dots) and the Z chromosome (black dot); Pearson correlation = 0.65 (*P <* 0.001). The effective rank provides a measure of the intrinsic dimensionality of the residuals. Panel (C) provides Bayesian estimates of regression coefficients for the association between chromosomal features and the PCvE of graphs with different numbers of admixture edges (details of each regression analysis are shown in Figure S7). Only autosomes were included in this analysis. Three chromosome features–size, mean recombination rate, and gene density–were summarized using PCA for these regression analyses; larger positive values of PC1 (which captured 69% of the variation in these features) denote larger chromosomes with lower recombination rates per bp and higher gene density. Consequently, positive regression coefficients (y-axis) indicate higher PCvE for chromosomes that are larger, experience less recombination, and have higher gene density (and vice versa for negative regression coefficients). Posteriors from the regressions are summarized in terms of the median (points), 50% equal-tail probability intervals (ETPIs; thick lines), and 90% ETPIs (thin lines); the Pearson correlation between the number of admixture edges and the point estimates of the regression coefficients was 0.89 (*P* = 0.0013). Panels (D), (E), and (F) depict population graphs for chromosomes 2, 4, and Z with no migration events, with branch lengths expressed in units of a drift parameter proportional to evolutionary change. Colored symbols and population IDs correspond to Table 1 and Figure 1. The *L. argyrognomon* population is included as an outgroup. Panels (G), (H), and (I) show graphs for the same three chromosomes with *m* = 2 migration edges (red arrows). The admixture proportions associated with each migration edge are indicated by the intensity of the red arrows; however, all migration edges shown had relatively high weights (0.30–0.45). See Figures S8–S30 for the full set of graphs for all chromosomes and for models with 0 to 5 admixture edges.

Chromosome size, recombination rate, and gene density were correlated with one another and jointly explained variation among autosomes in the proportion of allele-frequency covariance captured by population graphs (Figures 2C, S4, S5 and S6). However, this relationship depended on the number of migration edges. PC1 of these chromosomal features, where higher scores denote larger chromosomes with lower per-base-pair recombination rates and higher gene density, was negatively associated with covariance explained by graphs with zero or one migration edges, showed little association for intermediate numbers of migration edges, and became positively associated for graphs with five or more migration edges (Figure S7). Thus, bifurcating trees and graphs with little admixture fit better for smaller chromosomes with higher recombination rates and lower gene density, whereas the opposite was true for graphs with more migration edges. Consequently, the regression coefficient for PC1 changed sign and increased in magnitude with the number of migration edges (Pearson correlation = 0.89, *P* = 0.0013; Figure 2C).

We detected several admixture events that were consistent or largely consistent across chromosomes (Figure 2D-I and S8-S30). Here, we focus on events from graphs with one or two migration edges, as adding the first few edges produced the largest improvements in the proportion of allele frequency covariance explained and yielded the most easily interpretable results (note that an absence of admixture in the history of a population in the *m* = 2 graph does not preclude some admixture in that population’s history; Lipson, 2020). For autosomes with one migration edge, we found consistent evidence of admixture from *L. melissa* in the history of the Sierra Nevada populations, especially TIC and CLH, and sometimes CP. Based on the tree topology, these populations were often otherwise most closely related to *L. anna* (for chromosomes 21 and 22, the majority ancestry was instead from *L. melissa*, with minority ancestry from the ancestor of MR, CP, and two *L. anna* populations). The inferred *L. melissa* source populations were either BHP or the ancestor of BHP and VE (both from the western USA near the Sierra Nevada). Admixture (migration) weights indicated substantial contributions from this event, with a mean admixture weight or proportion (across chromosomes) of 0.39 (range = 0.27–0.48).

For the case of two migration edges, most chromosomes suggested the same (or similar) admixture event described above, along with evidence of (i) admixture between *L. melissa* and the two Warner Mountains populations (EP and BKM), which were otherwise most closely related to *L. anna* and the Sierra Nevada populations; (ii) admixture in the history of the SHC *L. anna* population; or (iii) admixture within the Sierra Nevada populations (CP, MR, both CP and MR, or all Sierra Nevada populations). These putative admixture events involved introgression from *L. anna*, *L. melissa*, the Sierra Nevada populations, or an ancestor of some subset of these (Figure 2D-I and S8-S30). Here too, the estimated levels of admixture were substantial (mean = 0.37, range = 0.16–0.50). Admixture in the history of MR was also detected for the Z chromosome. For the Z chromosome, we additionally found evidence of early admixture from an ancestral *L. melissa* population in the history of all Sierra Nevada populations (admixture proportions of 0.30 and 0.36). Patterns of admixture for graphs with three or more migration edges were increasingly complex and explained little additional allele frequency covariance. Nonetheless, even with many migration edges, the estimated migration weights remained considerable, consistent with major contributions of admixture to the genetic compositions and relationships of populations (e.g., for six migration edges, the mean weight was 0.34 for autosomes, with a range of 0.06–0.61).

### Patterns and causes of genomic variation in evolutionary relationships

We detected 19 unique chromosome-level tree topologies across the 23 *Lycaeides* chromosomes using CASTER (Figure 3); the CASTER-pair model produced similar results (Figure S31). The primary differences among chromosomes involved the placement of the Sierra Nevada (CP, MR, TIC, and CLH) and Warner Mountains (EP and BKM) populations, as well as the positions of GNP and HNV (both *L. idas*) relative to other lineages. The mean Robinson-Foulds distance between chromosome trees was 5.27 (range = 0.00–15.97), with the greatest differences occurring between the Z chromosome and the autosomes (mean = 11.79, range = 7.13–15.97). Overall, tree discordance among chromosomes exceeded null expectations, indicating significant heterogeneity in dominant evolutionary histories across the genome (average generalized Robinson–Foulds distance among chromosomes, null based on 50 permutations: mean = 1.86, range = 1.01–2.38, *P <* 0.02; Figures 3B, S31 and S32).

**Figure 3:**
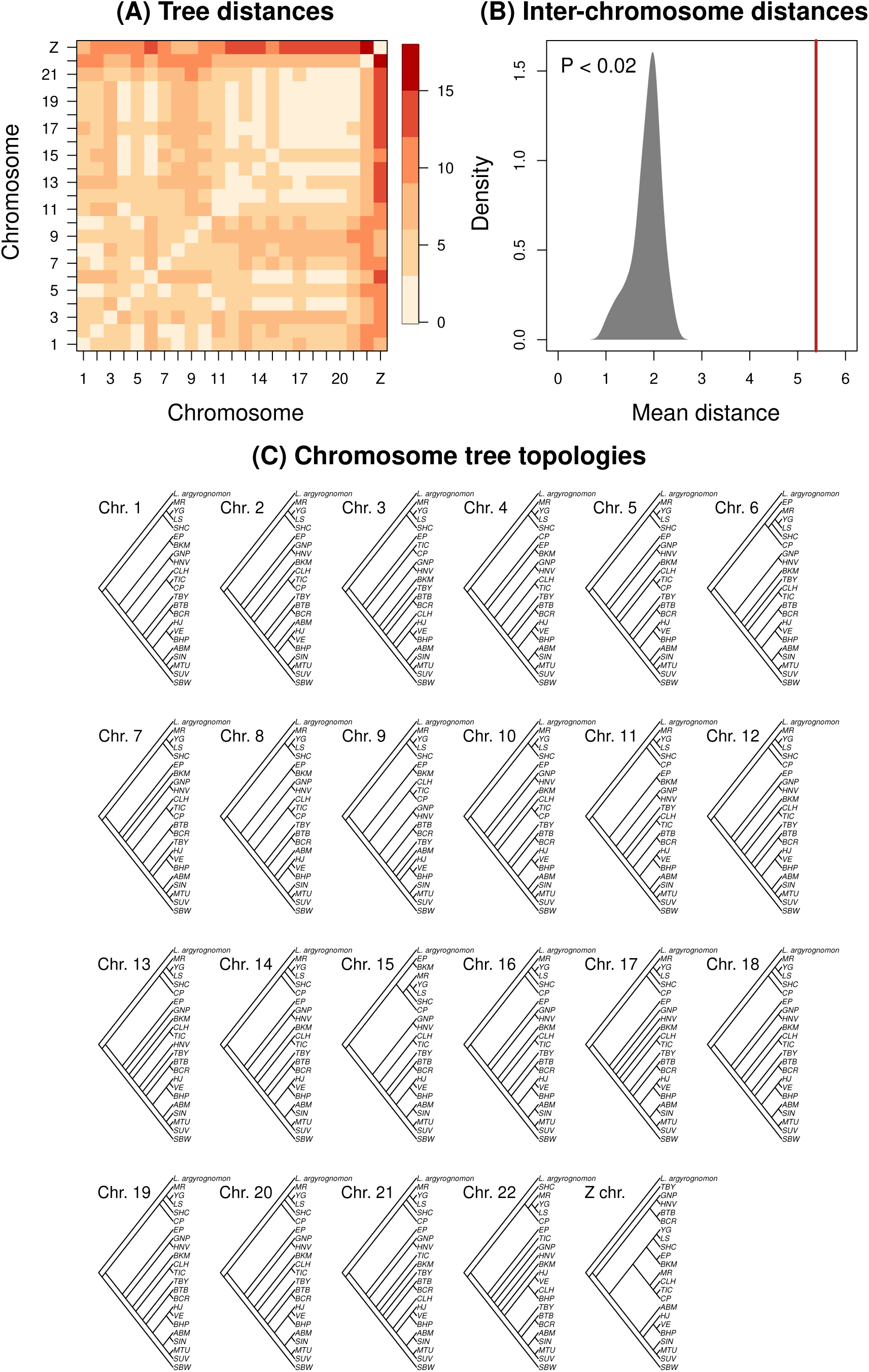
Summary of tree topologies from CASTER. Panel (A) depicts a heatmap showing generalized Robinson-Foulds distances between the tree topology for each pair of chromosomes; this metric is based on differences in the mutual clustering information between trees. Panel (B) shows the mean generalized Robinson-Foulds distance among trees for all pairs of chromosomes (vertical red line) relative to a null distribution (shaded gray region) generated from randomly assigned SNPs to chromosomes (the observed value exceeds null expectations with *P <* 0.02). Example topologies from one permutation are shown in Figure S32. Panel (C) shows the estimated tree topology (as a cladogram) from the CASTER-site model for each chromosome with *L. argyrognomon* as the outgroup. See Table 1 for population IDs.

Patterns of support for alternative four-taxon trees within and among chromosomes provided additional evidence for admixture involving the Sierra Nevada and Warner Mountains populations (Figure 4). Across the seven four-taxon analyses, variation in topology scores within chromosomes was similar in magnitude to variation among chromosome means (mean within = 0.0055, range = 0.0028–0.0098; mean among = 0.0045, range = 0.0019– 0.0083), indicating substantial heterogeneity at both scales. Below, we highlight several representative patterns, including evidence that they cannot be readily explained by ILS and are, in some cases, associated with recombination rate variation.

**Figure 4:**
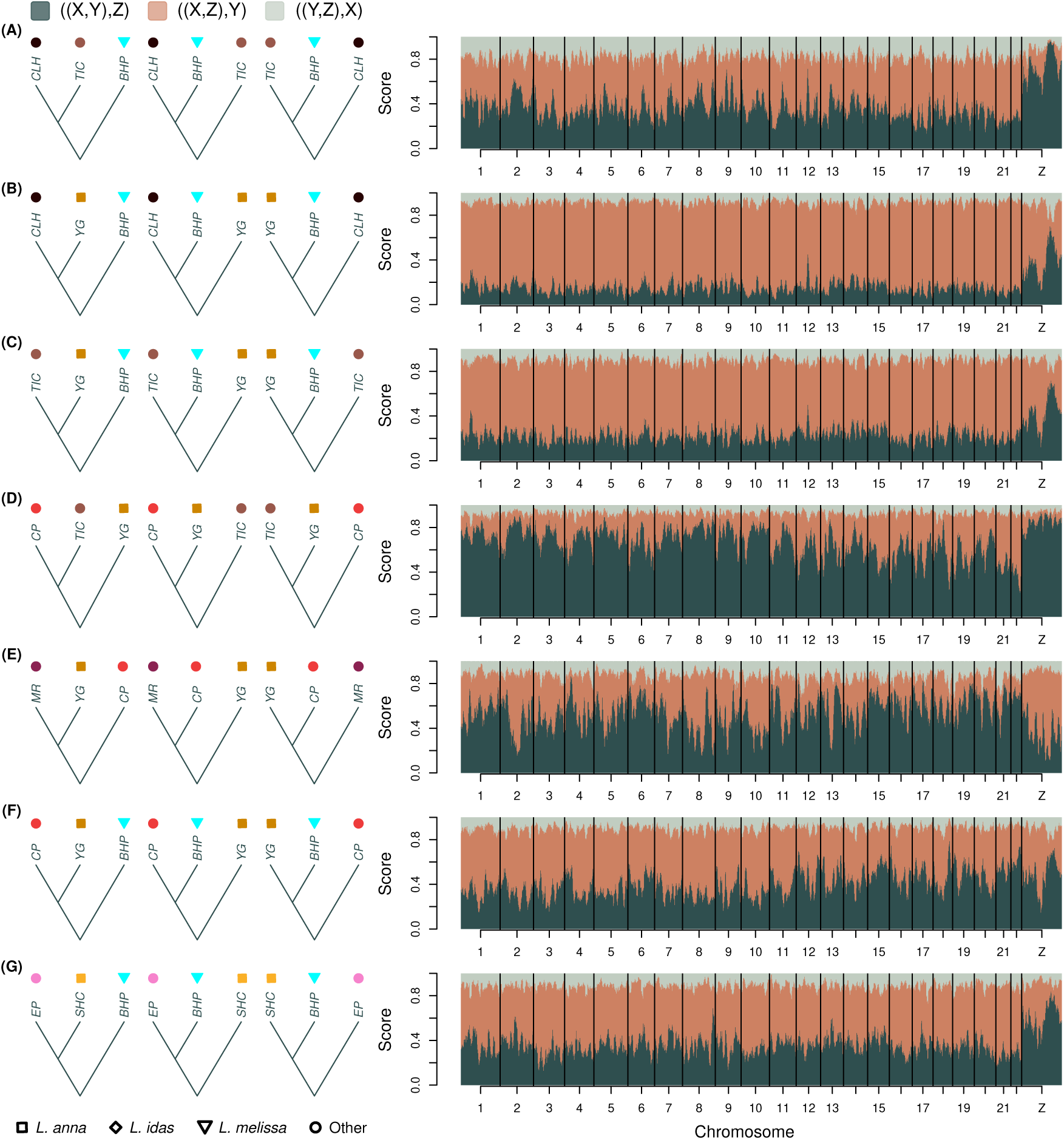
Genomic variation in four-taxon tree topologies for the Sierra Nevada and Warner Mountains populations (SBW, an Alaskan *L. idas* was always used as the outgroup and is not shown). Plots show normalized scores for three tree topologies, which were initially computed in 10 kilobase windows and then averaged over 50 kilobase overlapping sliding windows. Colors denote alternative topologies as shown in the legend, where X, Y and Z denote taxa as depicted to the left of the plots. The ((X,Y),Z) topology was defined to be consistent with the topology of the Bayesian chronogram from Figure 1B. Taxonomic designations for the populations shown in the figure are: *L. anna* = YG and SHC; *L. melissa* = BHP; Sierra Nevada = CP, MR, TIC and CLH; and Warner Mountains = EP.

We first considered seven four-taxon analyses involving Sierra Nevada or Warner Mountains populations (Figure 4). Trees grouping CLH and TIC (two Sierra Nevada populations) with each other, or with *L. anna* (YG) rather than *L. melissa* (BHP), received their strongest support on the Z chromosome, particularly across two large regions, despite strong genome-wide support for trees grouping CLH (and, to a lesser extent, TIC) with BHP (Figure 4A–C). Likewise, trees grouping CP with TIC or MR (all Sierra Nevada populations) were strongly supported across the Z chromosome and, to a somewhat lesser extent, on several large autosomes (Figure 4D,E). Distinct Z-linked patterns were also evident for the Warner Mountains population (EP), with increased support for a clade containing EP and *L. anna* (SHC) across much of the chromosome but weaker support near its center (Figure 4G). Across all seven analyses, support for the two alternative (non-dominant) topologies was significantly asymmetric for the autosomes and for the genome as a whole, and in five of seven cases for the Z chromosome, rejecting expectations under ILS (95% bootstrap confidence intervals; Figure S33A). Most analyses also showed greater phylogenetic signal (Σ) and significantly different asymmetry (Δ) on the Z chromosome than on the autosomes, consistent with lower levels of introgression on the Z chromosome (Table S1; Figures S33A, S34, and S35).

We next examined relationships among the Rocky Mountain populations (BTB and BCR) and nearby *L. melissa* and *L. idas* populations (Figure 5A–C), along with control four-taxon comparisons for which admixture was not suspected (Figure 5D–G). For both Rocky Mountain populations, most of the genome supported recent common ancestry with *L. melissa* (SIN) rather than *L. idas* (GNP) (Figure 5A,B). The primary exception was the Z chromosome, which strongly supported a Rocky Mountain–*L. idas* clade across much of its length; elevated support for this clade was also evident on portions of several autosomes (e.g., chromosome 7). Likewise, the geographically proximate *L. idas* population HNV showed moderate to strong support for a closer relationship with *L. melissa* (SIN) than with GNP across much of the genome, whereas most of the Z chromosome supported the alternative relationship (Figure 5C). Across these four-taxon analyses, asymmetric support for the two alternative topologies was detected only for the genome as a whole, but not for the autosomes or Z chromosome individually. Nevertheless, the Z chromosome consistently exhibited greater phylogenetic signal (Σ) and significantly different asymmetry (Δ) than the autosomes, consistent with the Z retaining a more bifurcating evolutionary history. (Table S1; Figures S33B and S36).

**Figure 5:**
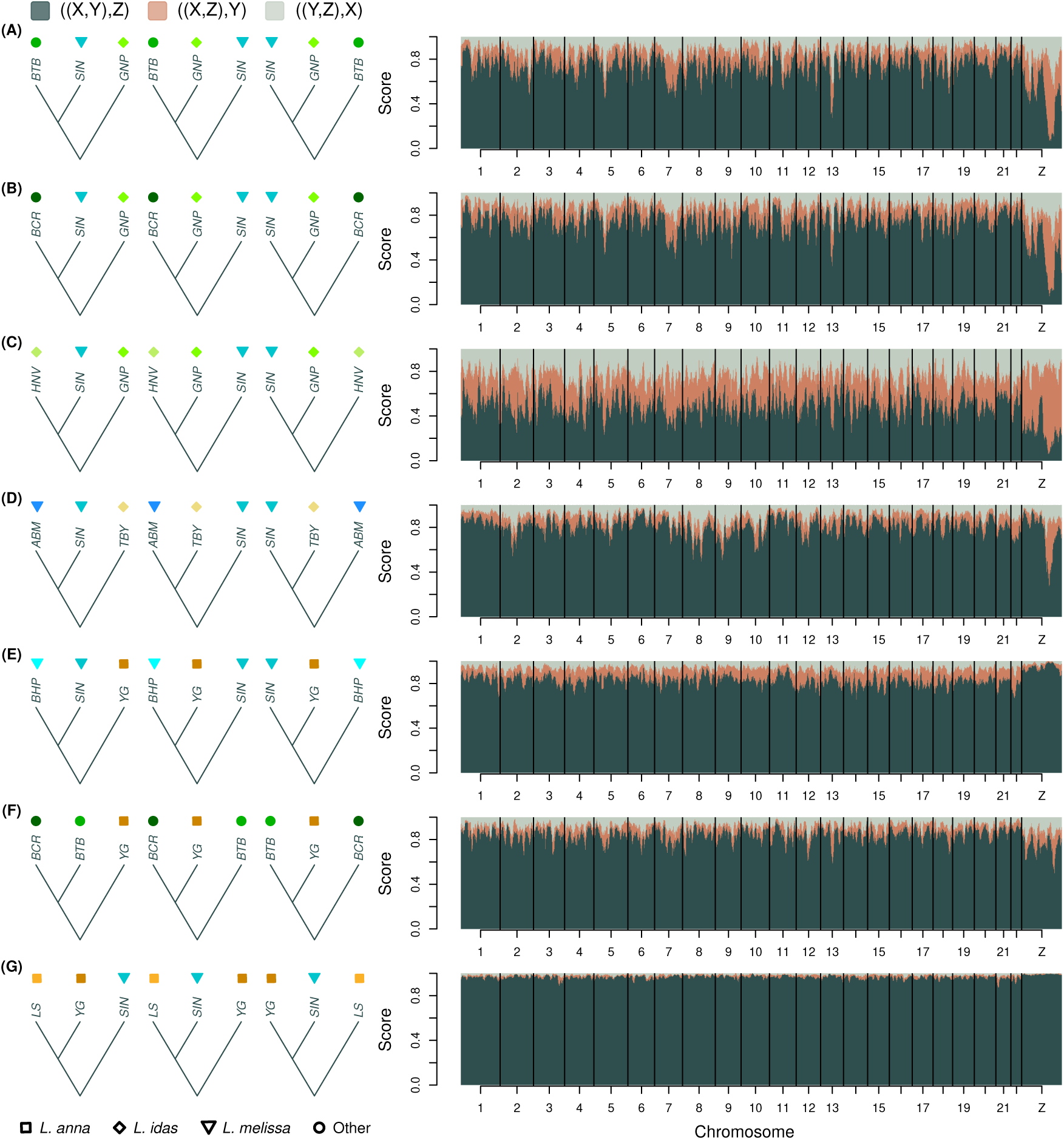
Genomic variation in four-taxon tree topologies for the Rocky Mountains populations and control groups, that is for cases where little or no admixture was expected. SBW, an Alaskan *L. idas* was always used as the outgroup and is not shown. Plots show normalized scores for three tree topologies, which were initially computed in 10 kilobase windows and then averaged over 50 kilobase overlapping sliding windows. Colors denote alternative topologies as shown in the legend, where X, Y and Z denote taxa as depicted to the left of the plots. The ((X,Y),Z) topology was defined to be consistent with the topology of the Bayesian chronogram from Figure 1B. Taxonomic designations for the populations shown in the figure are: *L. anna* = LS and YG; *L. idas* = GNP, HNV and TBY; *L. melissa* = ABM, BHP and SIN; and Rocky Mountains = BCR and BTB.

In contrast, the control four-taxon comparisons, which included either two *L. melissa*, two *L. anna*, or two Rocky Mountain populations together with a more distantly related lineage, showed much more consistent support for a single topology across the genome (Figures 5D–G). In most cases, support for the dominant topology was similar on the autosomes and Z chromosome, and there was little or no evidence of significant asymmetry in support for the alternative topologies, as expected under ILS (Table S1; Figures S33B and S36). The principal exception involved the high-elevation *L. melissa* population ABM, for which trees grouping ABM with the high-elevation *L. idas* population TBY rather than another *L. melissa* population (SIN) received moderate support across part of the Z chromosome. Although the Z chromosome also exhibited significantly greater phylogenetic signal (Σ), asymmetry (Δ) did not differ significantly from zero or from the autosomes (Figure 5D; Figures S33B and S36). Overall, these control analyses show that pronounced chromosome-specific variation in tree support is not an inevitable feature of the data.

Finally, we found only weak to moderate evidence that recombination rate variation was associated with finer-scale variation in phylogenetic signal (Σ) and ancestry asymmetry (Δ) across the 22 autosomes for which we have recombination-rate estimates and across all 14 four-taxon sets analyzed (Figures S37 and S38). Specifically, whether recombination rate was significantly associated with variation in Σ or Δ depended on the recombination map used (i.e., the population from which it was estimated), the four-taxon set analyzed, and whether the analysis controlled for differences in mean recombination rate among autosomes (see Figures S4, S5 and S6). Nonetheless, particularly for Σ and analyses involving the Sierra Nevada and Warner Mountains populations, the proportion of tests showing significant associations exceeded 5% (the expectation under the null hypothesis), and all significant associations with recombination rate were negative. These results suggest that higher recombination rates are, at least sometimes, associated with reduced phylogenetic signal (Σ) and reduced asymmetry in weights for alternative topologies (Δ). However, even when associations were significant, recombination rate always explained less than 1% of the variation in the topology-weight summaries (Σ or Δ).

### Genetic mapping of wing pattern

We detected moderate to substantial contributions of additive genetic variation to wing pattern traits in each of the three *Lycaeides* populations analyzed (mean point estimates of the proportion of variation explained [PVE]: GNP = 0.56, SIN = 0.72, and YG = 0.62; see Figure S39 for estimates and associated uncertainty for each of the 17 traits). Both the autosomes and the Z chromosome contributed to PVE for wing pattern traits, with particularly large contributions from the Z chromosome for a subset of traits in specific populations (e.g., a2, a4, a5, and m in GNP, and a6 in YG) (Figures 6, S2, S40, S41, and S42). The autosomes nevertheless explained most of the additive genetic variation on average (*>*90%). Relative to its genomic representation, however, the contribution of the Z chromosome was disproportionately large in GNP (8.9%) and YG (7.5%) (Table S2); the Z chromosome accounted for only 5.5% of SNPs included in the GWA analyses. Thus, wing pattern variation has a distributed genetic basis spanning both the autosomes and the Z chromosome, with an elevated contribution from the Z in at least two populations.

**Figure 6:**
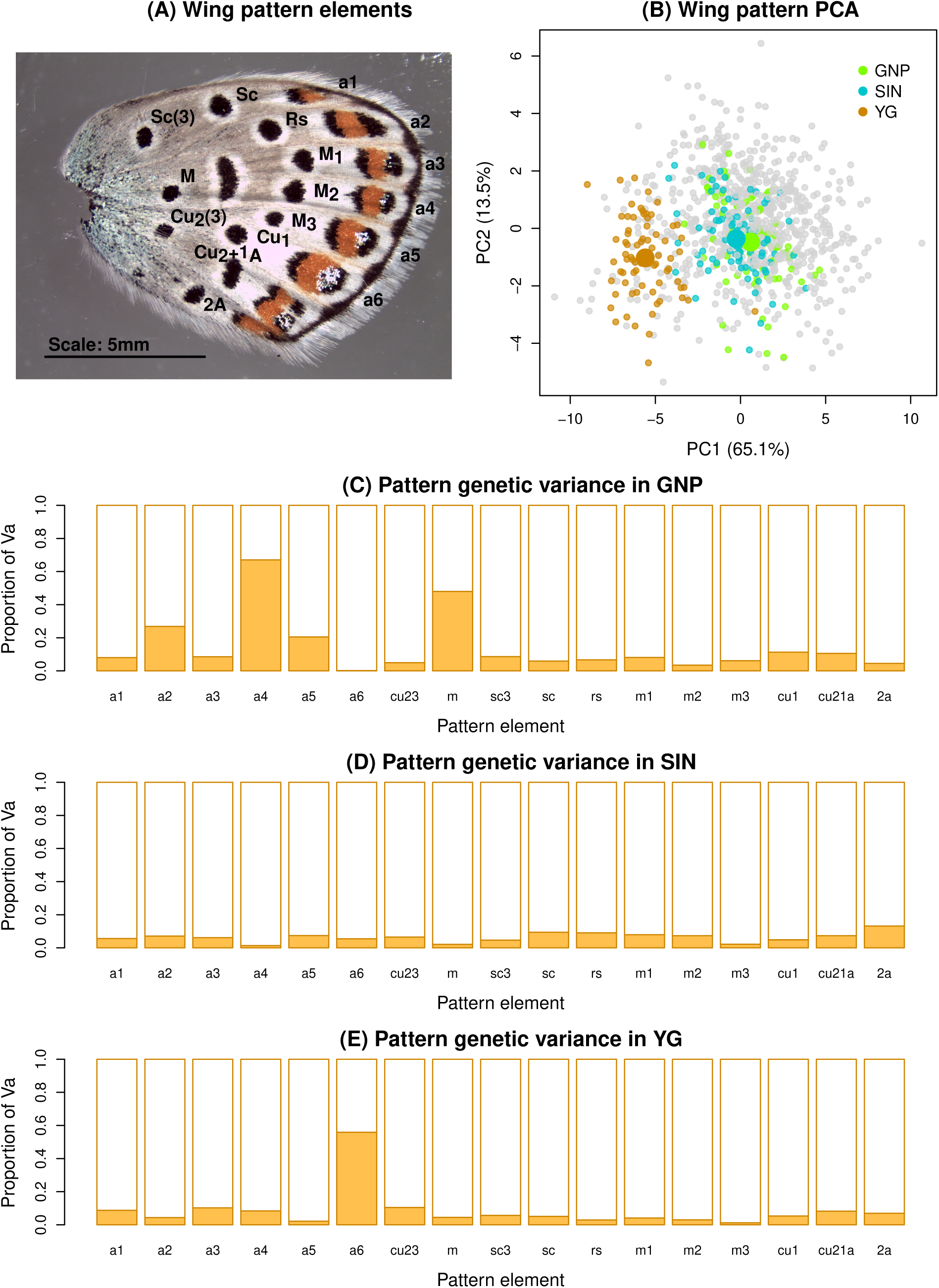
Summary of wing pattern mapping analysis. Panel (A) shows the ventral surface of the hind-wing with the measured black spots and aurorae (a2-a6) labeled. Panel (B) summarizes wing pattern variation in this system based on a principal components analysis. PCs 1 and 2 largely capture overall variation in spot and aurorae size and the relative sizes of spots versus aurorae respectively. Points denote PC scores for measured butterflies with colored points highlighting the specific populations analyzed in the current study. Panels (C), (D) and (E) show the proportion of the estimated additive genetic variation (Va) for each trait attributable to the autosomes (white region) versus the Z chromosome (orange region) for each population. Taxonomic designations for the three populations are: GNP = *L. idas*, SIN = *L. melissa*, and YG = *L. anna*.

## Discussion

To what extent can we describe the evolutionary history of a taxonomic group with a strictly bifurcating tree consisting of well-delineated species? The answer to this question is important for framing our understanding of speciation, species, and the structure of the living world. Our results suggest that, at least for *Lycaeides* butterflies, the answer varies across the genome. We have shown that the genomes of these butterflies comprise a mosaic of genetic regions and chromosomes with distinct ancestries, such that no single tree provides a sufficient summary of their evolutionary history. Variation in evolutionary history was of similar magnitude within individual chromosomes and among chromosomes. That said, the Z sex chromosome was exceptional, exhibiting a distinct and particularly tree-like history across the entire species complex. This genomic heterogeneity cannot be explained by ILS alone. Instead, it primarily reflects admixture and introgression with different outcomes on the autosomes and Z chromosome, with additional, more modest effects of genomic features such as recombination rate. The recent finding of a similarly tree-like evolutionary history for the Z chromosome in *Heliconius* butterflies suggests that this pattern may extend beyond *Lycaeides* (Thawornwattana *et al*., 2023). We discuss these results and their broader implications in more detail below.

### Evolutionary history of *Lycaeides*

Bayesian phylogenetic inference indicates that *Lycaeides* first colonized Alaska and then spread southward roughly one to two million years ago (also see, e.g., Nice *et al*., 2005; Vila *et al*., 2011). The nominal species *L. idas* appears to be paraphyletic and was likely the progenitor of both *L. anna* and *L. melissa* (and probably *L. samuelis*, though this remains to be tested), all of which are endemic to North America, specifically the USA and southern Canada. Our results strongly suggest that admixture has played a central role in the evolutionary history of the Sierra Nevada and Warner Mountain populations (consistent with, e.g., Gompert *et al*., 2006; Nice *et al*., 2013; Gompert *et al*., 2014). These populations received substantial ancestry contributions from *L. melissa* and *L. anna*, as well as from other ancestral populations more closely related to *L. anna* than to *L. melissa*, perhaps representing additional ancestral lineages of *L. idas*. The distinct placement of these admixed populations in some genomic trees, including the Z-chromosome tree, is consistent with these additional ancestry contributions. We found less extensive, but still compelling, evidence of admixture in the Rocky Mountain populations. In particular, these populations harbor predominantly *L. melissa* ancestry on the autosomes but *L. idas* ancestry on the Z chromosome. Past work suggested a larger genome-wide contribution of *L. idas* ancestry based on overall admixture proportions, but because those inferences relied on, or were heavily influenced by, ancestry-informative markers, they were likely disproportionately shaped by the Z chromosome (Gompert *et al*., 2010, 2012; Chaturvedi *et al*., 2020). Our results are also consistent with a north-south cline in the amount of *L. idas* ancestry in these populations. This cline likely reflects a combination of primary clinal divergence and post-Pleistocene secondary contact, together with recent and ongoing hybridization at the geographic margins of the region facilitated by the spread of feral alfalfa (*Medicago sativa*), a host used by *L. melissa* (Gompert *et al*., 2010, 2012; Chaturvedi *et al*., 2020; Forister *et al*., 2020; Gompert *et al*., 2022). Several lines of evidence support our interpretation of these patterns as consequences of gene flow, including admixture following secondary contact as well as primary divergence with gene flow: direct rejection of ILS as a sufficient explanation, the large chromosomal scale of ancestry differences, their clinal geographic structure, and direct evidence of ongoing hybridization (e.g., Gompert *et al*., 2010; Chaturvedi *et al*., 2020). Nevertheless, ILS almost certainly contributes to some of the finer-scale variation in evolutionary history among genomic regions.

### Phenotypic consequences of admixture in *Lycaeides*

One consequence of the mosaic evolutionary histories documented here is that admixture can bring together genomic regions with different ancestry in novel combinations (e.g., Lewontin & Birch, 1966; Le Corre *et al*., 2020; Todesco *et al*., 2020; Meier *et al*., 2023; Rosser *et al*., 2024). Past experiments have shown wing pattern elements have moderate effects on mate preference in *Lycaeides* butterflies (Fordyce *et al*., 2002; Gompert *et al*., 2013b). Because the autosomes and Z chromosome have markedly different ancestry histories in several admixed populations, our finding that both contribute substantially to wing pattern variation implies that admixture can reassort wing-pattern alleles drawn from different ancestral backgrounds. This prediction is supported by experimental crosses, which show that F1 hybrids exhibit mixtures of intermediate and transgressive wing patterns and male genitalic morphologies (another trait strongly distinguishing nominal species in this group) (Lucas *et al*., 2008, 2026). The role of novel combinations of mixed-ancestry autosomes and sex chromosomes for other ecologically important traits remains to be determined. Among such traits, diapause is of particular interest. *Lycaeides* overwinter as diapausing neonate larvae within eggs. Diapause is facultative in *L. melissa*, allowing multiple generations per year, whereas in *L. idas*, *L. anna* and all known admixed populations, it is obligate and only one generation occurs annually (Gompert *et al*., 2013b). In at least some other Lepidoptera, obligate versus facultative diapause is controlled by known clock genes located on the Z chromosome (Jin *et al*., 2024). If diapause has a similar Z-linked genetic basis in *Lycaeides*, retention of *L. idas*- or *L. anna*-like Z ancestry could help explain why admixed populations exhibit obligate diapause despite substantial *L. melissa* ancestry elsewhere in the genome. We do not yet know whether this is true, but the admixed populations analyzed here generally have greater *L. idas* or *L. anna* ancestry on the Z chromosome than *L. melissa* ancestry, consistent with this hypothesis. More broadly, mosaic ancestry provides a mechanism for assembling phenotypic combinations that differ from those associated with any single parental genomic background. This potential is well documented for traits with relatively simple genetic architectures (Mavárez *et al*., 2006; Jiggins *et al*., 2008) but also applies to polygenic traits, such as wing pattern in *Lycaeides* butterflies.

### Implications for understanding species and speciation

This combination of genomic mosaicism and persistent organismal differentiation raises a broader question of what constitutes a species in a system with extensive gene flow. From a whole-organism perspective, our results, combined with past work (Fordyce *et al*., 2002; Nice *et al*., 2002; Lucas *et al*., 2008; Gompert *et al*., 2013b; Lucas *et al*., 2018), suggest that *Lycaeides* are best viewed as a complex of recently diverged taxa in the early stages of the speciation process. Phenotypic and ecological differences exist among these taxa, along with a modest degree of pre- and postzygotic reproductive isolation (Fordyce *et al*., 2002; Gompert *et al*., 2013b; Lucas *et al*., 2026). Yet hybridization has been common in the past, and fertile hybrids are readily produced both in the lab and in natural hybrid zones (Gompert *et al*., 2006; Nice *et al*., 2013; Gompert *et al*., 2014; Chaturvedi *et al*., 2020; Lucas *et al*., 2026). However, speciation need not be viewed solely from a whole-organism perspective; instead, it can proceed at different rates across different regions of the genome, as evidenced by the semipermeable nature of species boundaries and emphasized by the genic view of speciation (Wu, 2001; Harrison & Larson, 2014; Westram *et al*., 2022). We think that such a genic view of speciation provides a useful context for interpreting the distinct histories and patterns of admixture for autosomes versus the Z sex chromosome in *Lycaeides*.

At least when gene flow is possible, progress toward speciation depends on an antagonism between divergent selection, which promotes speciation, and gene flow and recombination, which homogenize allele frequencies and break up locally adaptive gene complexes (Felsenstein, 1981; Barton & Bengtsson, 1986; Flaxman *et al*., 2013). Thus, genomic regions experiencing stronger selection or lower rates of recombination are expected to behave more like species, that is, as distinct, independently evolving lineages. Several mechanisms could therefore cause sex chromosomes to effectively speciate before the rest of the genome, either because they have disproportionate effects on hybrid fitness (stronger selection; i.e., the large X effect) or because of reduced recombination in the heterogametic sex (lower recombination) (Orr, 1997; Carling & Brumfield, 2008; Presgraves, 2008; Lenormand & Roze, 2025). Because recombination does not occur for any chromosomes (autosomes or sex chromosomes) in female butterflies, the heterogametic sex, a greater proportion of Z-chromosome copies than autosomal copies have an opportunity to recombine (∼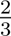 of Z chromosomes are in males, compared with 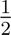 of autosomes). However, assuming, at least as a first approximation, that the effective population size of the Z is 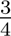 that of the autosomes, its greater opportunity for recombination per chromosome copy is offset by its smaller effective population size. Thus, all else being equal, one would expect similar population recombination rates for the autosomes and Z chromosome. We therefore hypothesize that stronger selection, rather than reduced recombination alone, contributes to the paucity of admixture in the evolutionary history of the Z chromosome in *Lycaeides*.

More generally, the antagonism between selection and recombination suggests that introgression should be rarer on chromosomes or in genomic regions where recombination rates are low or targets of selection are numerous (or selection is stronger). Such patterns have been documented in several systems (see, e.g., Schumer *et al*., 2018) and, all else being equal, should result in more tree-like histories in these genomic regions. For example, recombination per base pair is often lower on longer chromosomes, contributing to a commonly observed negative relationship between chromosome length and minor-parent ancestry or introgression (e.g., Brandvain *et al*., 2014; Schumer *et al*., 2018; Martin *et al*., 2019; Chaturvedi *et al*., 2020; Nouhaud *et al*., 2022). However, these expectations are based largely on relatively simple cases involving pairs of taxa with a clear distinction between major- and minor-parent ancestry. Some of our results are consistent with these expectations. For instance, across the autosomes, we found weak but consistent evidence that support for a single topology was greater in genomic regions with lower recombination rates, as expected if such regions experience less introgression. However, our TreeMix analyses revealed a more complex pattern. Bifurcating trees (and graphs with one migration edge) explained more allele-frequency covariance for smaller chromosomes with higher recombination rates and lower gene densities, whereas graphs with many migration edges explained more covariance for larger chromosomes with lower recombination rates and higher gene densities. One possible explanation is that departures from a bifurcating history are relatively diffuse on high-recombination chromosomes, such that their average covariance structure is reasonably well approximated by a tree but additional migration edges provide relatively little improvement. In contrast, residual covariance on low-recombination chromosomes might be concentrated in a smaller number of strong, coherent signals that are poorly represented by a bifurcating tree but efficiently captured by additional migration edges. Consistent with this interpretation, bifurcating trees explained a greater proportion of allele-frequency covariance for chromosomes whose TreeMix residuals had higher effective rank, that is, chromosomes for which the unexplained covariance was distributed across more effective dimensions, although this relationship was strongly influenced by the Z chromosome. Thus, the genomic consequences of admixture may depend not only on the total amount of introgression, but also on how introgressed ancestry is distributed among loci and populations and how these signals combine with variation arising from ILS. Other factors could also contribute to variation in the outcomes of admixture, including the effective population sizes (and thus genetic loads) of the hybridizing taxa and ecological selection favoring particular ancestry combinations (e.g., Gompert *et al*., 2012; Matute *et al*., 2020; Nouhaud *et al*., 2022; Springer *et al*., 2025).

The same logic may extend to finer genomic scales. At finer genomic scales, chromosomal rearrangements, especially inversions, can reduce effective recombination among linked loci and thereby strengthen the ability of selection to maintain differentiated combinations of alleles (particularly if they lock together co-adapted sets of alleles in supergenes), and thus may in some cases represent the leading edge of the speciation process. Previous work suggests that genomic regions harboring structural variation (in this case, primarily large deletions) are resistant to introgression in a *Lycaeides* hybrid zone, but we do not yet know the contribution of these chromosomal mutations to broader patterns of ancestry and introgression across the species complex (Zhang *et al*., 2023). Regardless of their specific relevance for *Lycaeides*, it is now clear that chromosomal rearrangements often underlie ecologically important differences between species as well as polymorphisms within species (e.g., Lowry & Willis, 2010; Koch *et al*., 2021; Akopyan *et al*., 2022; Nosil *et al*., 2023; Gompert *et al*., 2025). Such rearranged genomic regions can therefore become evolutionarily semi-independent entities before substantial divergence evolves across the remainder of the genome; when rearrangements remain polymorphic, such entities can even coexist within a single organismal population. More generally, these observations support a hierarchical view of biodiversity in which evolutionary entities can exist at multiple genomic and organismal scales, with different boundaries, histories, and degrees of evolutionary independence.

## Data and code availability

The *Lycaeides* pooled whole genome sequence data have been deposited in the NCBI SRA (BioProject PRJNA1375794). Wing pattern data are available from Dryad (https://doi.org/10.5061/dryad.fc827). Code and associated documentation are available at https://github.com/zgompert/LycAdmixMosaic. Final versions of computer code will be deposited on Zenodo when the manuscript is accepted.

## Author contributions

Conceptualization: Z.G., L.K.L., M.L.F., J.A.F., and C.C.N.; Formal analysis: Z.G.; Funding acquisition: Z.G.; Investigation: Z.G., L.K.L., A.D., M.L.F., J.A.F., and C.C.N.; Validation: Z.G.; Visualization: Z.G.; Writing–original draft: Z.G.; Writing–review and editing: Z.G., L.K.L, M.L.F., J.A.F., and C.C.N.

## Acknowledgments

We thank the following for collecting or providing specimens: Alex Buerkle, Samridhi Chaturvedi, Megan Jamison, Vladimir Lukhatanov, Amy Springer, Gerard Talavera, and Roger Vila. Thanks to Natalie Yoho for help with lab work. The support and resources from the Center for High Performance Computing at the University of Utah are gratefully acknowledged. We thank Diana Tataru for comments on an earlier draft of this manuscript.

Online Supplemental Materials

## Supplemental Methods

### Library preparation, sequencing and read filtering

Sequencing libraries were constructed by BGI. In brief, genomic DNA was fragmented with a Covaris ultrasonicator and fragment size distributions were verified on an Agilent 2100 Bioanalyzer. Fragmented DNA underwent end-repair and 3’ dA-tailing followed by bead-based cleanup. Indexed adapters were ligated to A-tailed fragments and ligation products were purified with magnetic beads. Adapter-ligated DNA was then amplified with KAPA HiFi HotStart DNA Polymerase. Amplified libraries were then converted to single-stranded circular DNA using an MGI circularization workflow. DNA nanoballs were generated from the single-stranded circles by rolling-circle amplification. Sequencing was performed using the DNBseq platform with paired-end 100 bp reads and a target read depth of 100× for each library (read depths obtained were ∼114×, see Table 1). After sequencing, the raw reads were filtered. Data filtering was accomplished with SOAPnuke and included removing adaptor sequences, contamination and low-quality reads from raw reads. Specifically, adapters were removed if reads matched 25% or more of the adapter sequence (with a 2 bp maximum mismatch) and reads were discarded if 40% or more of the bases had quality scores less than 10 or if Ns comprised more than 0.1% of the read (i.e., ≥1 N).

### Alignment, variant calling, filtering and allele frequency estimation

We aligned the sequence reads to the *L. melissa* genome (Zhang *et al*., 2023) using bwa-mem2 (version 2.0pre2) with default parameters (Li & Durbin, 2009; Vasimuddin *et al*., 2019). PCR duplicates were then marked and removed with samtools (version 1.16) using the collate, fixmate and markdup commands (Li *et al*., 2009; Ebbert *et al*., 2016). We called genetic variants using bcftools consensus caller (option -c) (version 1.16) (Li, 2011). For this, we skipped alignments with mapping quality less than 20 and bases with quality scores less than 30, ignored insertion-deletion polymorphisms, and only called SNPs if the probability that all populations were fixed for the reference allele given the data was less than 0.01. We then filtered the initial SNP set to retain only SNPs that were bi-allelic, had qualities *>*30, total read coverage *>*1350, and base-quality bias, mapping-quality bias, and read-position bias Z-scores *<* ±3. We used GATK (version 4.1.4.1) (McKenna *et al*., 2010) for filtering.

We extracted the counts of high-quality reads supporting the reference and alternate allele for each SNP and population. We used these counts to obtain maximum likelihood estimates of allele frequencies, which are equivalent to the sample allele frequencies computed from the read counts. This was done in R. We used these allele frequency estimates for principal component analysis (PCA) and for generating input sequences for phylogenetic analyses, as described in the next section. For PCA, we used non-reference allele frequencies of 0.001 for populations with no reads (fully missing data) for a given SNP.

### Data processing, models and priors for BEAST

We used BEAST version 2.7.7 to estimate a time-calibrated phylogeny from a subsample of genome-wide SNPs (Drummond & Rambaut, 2007; Bouckaert *et al*., 2019). As noted in the main text, a concatenated alignment of 5408 SNPs distributed across the genome was used for computational feasibility. Specifically, we retained 0.035% of the original SNP set by randomly sampling SNPs from the full dataset after excluding sites with missing data (zero reads) in any population. To generate a consensus sequence for each population, variable sites were coded as the major allele when its frequency was at least 0.95. Sites that did not meet this criterion were coded as missing data (N).

To account for ascertainment bias associated with SNP-only data, invariant sites were incorporated into the analysis. We first calculated the total number of each nucleotide in the genome and then subtracted the number of SNPs (i.e., variant sites) observed for each nucleotide class. Because only 0.035% of sites were included in the BEAST analysis, these counts were multiplied by 0.00035 to obtain the corresponding numbers of invariant sites represented in the subsampled data: A = 50,190, C = 28,322, G = 28,284, and T = 50,114.

We used the GTR model of sequence evolution with gamma-distributed rate heterogeneity approximated using four discrete categories. Substitution-model parameters were estimated from the data using the default priors (gamma distributions) implemented in BEAST. We used the Coalescent Bayesian Skyline tree prior to allow effective population size to vary through time, with Jeffreys-type priors on the effective population sizes.

We employed the Random Local Clock (RLC) model, which can be viewed as a generalization of both strict and relaxed molecular-clock models because the number and placement of rate shifts are estimated as part of the analysis (Drummond & Suchard, 2010). We specified two calibration nodes, each with an informative prior based on divergence-time estimates from the literature. Specifically, we placed a normal prior on the age of the most recent common ancestor of the North American *Lycaeides* ingroup with a mean of 1.29 million years (standard deviation [SD] = 0.167), based on the calibrated global butterfly phylogeny of Kawahara *et al*. (2023). We also placed a normal prior on the divergence between *L. argyrognomon* and the North American *Lycaeides* lineage with a mean of 2.125 million years (SD = 0.167), corresponding to the average of four estimates reported by Vila *et al*. (2011). This divergence is thought to coincide with the colonization of North America by *Lycaeides*. The overall molecular clock rate was estimated from the data, but the starting value was set to 0.0029 based on a mutation-rate estimate from *Heliconius* butterflies (*µ* = 2.9 × 10*^−^*9) (Keightley *et al*., 2015).

We used coupled MCMC to facilitate mixing and convergence, with four heated chains and one cold chain (delta temperature = 0.025) (Müller & Bouckaert, 2020). We performed six independent coupled-MCMC analyses, each consisting of five chains and 500,000,000 iterations. Parameter values were logged every 1000 iterations, whereas trees were sampled every 5000 iterations.

We used LogCombiner (version 2.7.5) to combine log and tree files and to assess mixing and convergence. A 20% burn-in was removed from each run prior to combining, and the log files were further thinned by a factor of five. This yielded 480,006 posterior samples of parameter values and 120,006 sampled trees. We then used TreeAnnotator to generate a maximum clade credibility tree with median node heights (Heled & Bouckaert, 2013), and visualized the resulting tree using FigTree (version 1.4.4) (Rambaut, 2018).

### Residual structure for TreeMix

We examined the extent to which residuals from TreeMix population trees (graphs with 0 migration edges) exhibited similar structure across chromosomes, which would suggest consistency in the populations that deviated most strongly from expectations under a strictly bifurcating tree. For each chromosome, we used PCA to summarize the structure of the residual matrix from the population tree. Specifically, we analyzed the centered, but not standardized, residual matrix using the prcomp function in R. This matrix contains residuals for every pair of populations, with positive residuals indicating population pairs that are more genetically similar than expected from the tree and negative residuals indicating pairs that are less genetically similar than expected.

We then used several complementary approaches to compare residual structure among chromosomes. First, we computed Pearson correlation coefficients between the PC1 scores from each pair of chromosome-specific PCAs. Second, we conducted a PCA on a matrix of PC1 scores across chromosomes, where chromosomes constituted the observations and the PC1 scores for the 23 populations constituted the variables. This allowed us to visualize similarities among chromosomes in a lower-dimensional space and identify groups of chromosomes with similarly structured residuals. Finally, we computed the effective rank (also known as the participation ratio) of each residual matrix from the original PCA analyses, which provides a measure of the effective number of dimensions contributing to variation in the residual matrix. Residual matrices with lower effective rank have fewer important dimensions, that is, fewer dimensions contributing substantially to the total variation in the residuals. We calculated the effective rank as 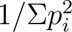, where *p_i_* denotes the normalized eigenvalue (*λ_i_*), *p_i_* = *λ_i_/*Σ*_j_λ_j_*.

### Estimating recombination-rate landscapes

We used iSMC to estimate population recombination-rate landscapes for five *Lycaeides* lineages, each represented by moderate-coverage (∼8×) whole-genome sequence data (HiSeq 2000, 2×100-bp paired-end reads, *>*20 million high-quality [≥Q30] reads per individual) from a single female butterfly: *L. anna* (Yuba Gap, CA), *L. idas* (Trout Lake, WY), *L. melissa* (Bonneville Shoreline Trail, UT), the Sierra Nevada lineage (Carson Pass, CA), and the Warner Mountain lineage (Buck Mountain, CA) (Barroso *et al*., 2019). These data were originally generated for *de novo* genome assembly and are described in Chaturvedi *et al*. (2020). Because iSMC infers recombination rates from patterns of heterozygosity and females are heterogametic in butterflies (resulting in little or no heterozygosity on the Z chromosome), we restricted this analysis to autosomes.

We first aligned the sequence data to the *L. melissa* reference genome (Zhang *et al*., 2023) using bwa-mem2 (version 2.0pre2), as described above for the pooled whole-genome sequence data (Li & Durbin, 2009; Vasimuddin *et al*., 2019). Likewise, we used samtools (version 1.16) to remove PCR duplicates and bcftools (version 1.16) for variant calling (Li *et al*., 2009; Li, 2011; Ebbert *et al*., 2016). We excluded alignments with mapping quality scores less than 20 and bases with quality scores less than 30, ignored insertion-deletion polymorphisms, and called SNPs only when the probability that the site was invariant for the reference allele given the data was less than 0.01. Variant calling was performed separately for each lineage, resulting in five distinct VCF files. We filtered each VCF file to retain only SNPs supported by at least five sequence reads. Each VCF file also included invariant sites, which provide information on sequence coverage across the genome and are needed to distinguish homozygosity from missing data.

We then used iSMC (version 1.0.0) to estimate population recombination rates (*ρ* = 4*N_e_r*) from each filtered VCF file (Barroso *et al*., 2019). Estimates of *ρ* from iSMC are based on genome-wide patterns of heterozygosity in individual genomes. We assumed five *ρ* categories and 30 intervals, consistent with default recommendations of Barroso *et al*. (2019). We estimated *ρ* at three genomic scales using window sizes of 25 kbp, 250 kbp, and 1 Mbp. For each 25-kbp window, we calculated a composite recombination-rate estimate as the average of the estimates for the corresponding 25-kbp, 250-kbp, and 1-Mbp regions containing that window. These estimates were then averaged across lineages and, where relevant, across entire chromosomes for downstream analyses.

### Bayesian regression methods

We conducted a series of Bayesian regression analyses to determine whether, and to what extent, chromosomal features explained differences among chromosomes in the proportion of allele frequency covariance explained by a population tree (no migration edges) or graph (one or more migration edges) inferred with TreeMix. We first considered three chromosome-level covariates: chromosome length, mean recombination rate, and gene density. For recombination rate, we used the mean estimate of the population recombination parameter (*ρ*) for each chromosome, averaging across lineages and chromosomal regions (see “Estimating recombination-rate landscapes” above). Chromosome length, measured in base pairs, was obtained from the *L. melissa* reference genome (Zhang *et al*., 2023), and gene density was calculated from an existing genome annotation as the proportion of each chromosome annotated as coding sequence (CDS) (Goodwin *et al*., 2026).

We fit Bayesian general linear models with these covariates and their pairwise interactions as independent variables and the proportion of allele frequency covariance explained by the TreeMix tree or graph as the dependent variable. Analyses were conducted separately for graphs with 0 to 8 migration edges; in each case, the sample size was *n* = 22 autosomes. We compared models containing different subsets of covariates using approximate leave-one-out cross-validation, with predictive performance summarized by the expected log pointwise predictive density (ELPD) (Vehtari *et al*., 2017). The best-fitting model, defined as the model with the highest ELPD, varied among graphs with different numbers of migration edges. However, chromosome length was included most frequently, appearing in the best model for five of the nine migration-edge values considered. Because the covariates were highly correlated (absolute Pearson correlations among variable pairs ranged from 0.39 to 0.76), we also evaluated a model using PC1 from a principal component analysis (PCA) of the three (centered and standardized) covariates. PC1 explained 68.6% of the variation in the three covariates. Higher PC1 values corresponded to larger chromosomes with higher gene density and lower recombination rates per base pair, as indicated by loadings of 0.62, 0.47, and −0.63, respectively. The model with PC1 as the only covariate performed well overall (median rank across the nine migration-edge values = 2) and was therefore used for the analyses described in the main text.

We fit all models using the rstan interface to Stan and Hamiltonian Monte Carlo (HMC) sampling (Betancourt, 2017; Neal, 2011; Stan Development Team, 2019). Each model was run with four chains, each consisting of 4000 iterations, including a burn-in period of 2000 iterations. We used the loo package (version 2.8.0) to calculate ELPD values from the HMC output (Vehtari *et al*., 2024). We placed normal priors (mean = 0, SD = 10) on all regression coefficients and a half-normal prior on the residual standard deviation (SD = 2).

### Tests of incomplete lineage sorting and its alternatives

We summarized and analyzed the window-based unnormalized topology scores from CASTER to formally test whether heterogeneity in topologies across the genome could be explained by incomplete lineage sorting (ILS) alone. We focused on two main summaries. First, we used the total (summed) unnormalized score (denoted Σ), that is, the sum of the scores for the three possible four-taxon topologies, as a measure of phylogenetic signal (consistent support for a single topology) for each window (see equation S12 in Zhang *et al*., 2025). Windows with strong support for a single genealogy will have high values of Σ, whereas windows with weak or ambiguous phylogenetic resolution will have lower values. Under a simple ILS model, discordant gene genealogies are expected to be associated with lower phylogenetic signal on average, whereas windows that support alternative topologies and exhibit strong phylogenetic signal are more likely to reflect additional processes, such as gene flow or introgression (Zhang *et al*., 2025).

Second, under the multispecies coalescent with ILS, the expected proportions of loci (windows) supporting the primary (species-tree) topology versus the two alternative four-taxon topologies (where one species is the outgroup) are 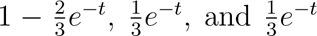, where *t* is the internal branch length in coalescent units (Rosenberg, 2002; Degnan & Rosenberg, 2009). Most importantly, regardless of the value of *t*, the two non-species-tree (i.e., alternative) topologies should occur with equal frequency if ILS is the sole cause of genomic heterogeneity in evolutionary histories. Substantial asymmetry therefore requires additional processes such as hybridization, gene flow, ancestral population structure, or selection (Degnan & Rosenberg, 2009; Zhang *et al*., 2025). Similar logic underlies ABBA–BABA tests of introgression (Green *et al*., 2010; Durand *et al*., 2011; Martin *et al*., 2015). Thus, deviations from equal frequencies of the alternative topologies, particularly when combined with strong phylogenetic signal supporting those deviations, can provide evidence of processes other than ILS, especially ancient or ongoing hybridization or complex patterns of gene flow. Selection could also contribute to such patterns, although its effects are more likely to be localized to specific genomic regions rather than genome-wide.

Our focus was not on detecting introgression (or non–ILS more broadly) for individual windows but rather on testing whether ILS alone could explain differences in ancestry patterns between autosomes and the Z chromosome. To this end (for each four-taxon set), we calculated the total score (Σ) for each window along with the difference in scores for the alternative topologies (denoted Δ). Alternative topologies were defined relative to the primary topology, which was determined either from the species tree inferred using BEAST or from the sum of scores for each topology across the entire genome (these were generally the same, although they differed in a few cases where the autosomes and Z supported different topologies). We then calculated differences in Σ and Δ between the Z chromosome and autosomes.

We used a permutation procedure to test whether phylogenetic signal was greater on the Z than on the autosomes (Σ*_Z_ >* Σ*_A_*, where *Z* and *A* denote the Z chromosome and autosomes, respectively) and whether the symmetry of the alternative topologies differed between the Z and autosomes (Δ*_Z_* ≠ Δ*_A_*). Evidence of Σ*_Z_ >* Σ*_A_* combined with Δ*_Z_* ≠ Δ*_A_* would be consistent with different evolutionary histories for the Z chromosome versus autosomes. One plausible explanation for such a pattern is differential introgression of the Z chromosome relative to autosomes. For example, elevated Σ*_Z_* (phylogenetic signal) could arise if the Z chromosome is resistant to introgression or if the Z has experienced adaptive introgression accompanied by more extensive introgression of autosomes. The analysis was conducted on thinned data sets (retaining every 5th window with different starting points) to reduce the effects of non-independence among neighboring 10 kb windows; 1000 permutations were used to generate null distributions.

As an additional assessment of ILS as the cause of heterogeneity in tree topologies, we computed bootstrap confidence intervals for Δ for the Z chromosome, autosomes, and the genome as a whole. In each case, ILS predicts Δ ≈ 0, and thus that confidence intervals should overlap 0 (Degnan & Rosenberg, 2009; Zhang *et al*., 2025). For this analysis, the primary and alternative topologies were defined based on the sum of scores for (i) the Z chromosome only, (ii) the autosomes only, and (iii) the whole genome. We then calculated the mean Δ across windows for the corresponding sets of windows. This analysis tests for variation in topologies within the Z chromosome, autosomes, or genome as a whole that cannot be readily explained by ILS. For example, introgression involving the entire Z chromosome could produce signals at the genome level but not necessarily lead to deviations from ILS when considering variation in topology along the Z chromosome itself. As above, bootstrap confidence intervals for Δ were estimated using thinned (every 5th) 10 kb windows to minimize the effects of non-independence among neighboring loci. Thinned windows were resampled with replacement 1000 times to compute bootstrap confidence intervals (the starting point for thinning, that is, the 1st, 2nd, 3rd, 4th, or 5th window, was varied among bootstrap replicates).

Finally, we fit a series of regression models to determine whether recombination-rate variation was associated with finer-scale variation in phylogenetic signal (Σ) and ancestry asymmetry (Δ) across the 22 autosomes for which we had recombination-rate estimates. We did this for all 14 four-taxon sets described in the main text, using each of the five recombination maps as well as an average map. We fit regressions relating window-based estimates of Σ and Δ (50-kb windows) to overlapping window-based estimates of *ρ* (25-kb windows). For each of the 14 four-taxon sets and six recombination maps (including the average map), we fit both ordinary linear regressions, with recombination rate as the sole predictor variable, and linear mixed models with chromosome included as a random effect. The latter allowed us to ask whether recombination rate explained variation in ancestry patterns after accounting for differences among chromosomes. These models were fit in R using the lm and lmer functions, respectively; lmer was implemented in lme4 version 2.0.1 (Bates *et al*., 2015).

### Wing pattern analyses

We conducted a PCA of wing pattern measurements (the sizes of 17 wing pattern elements) for the full set of *Lycaeides* butterflies from Lucas *et al*. (2018) (*n* = 1000 butterflies, including the 295 butterflies used for genetic mapping) (see Figure S2 for representative wing images). Methods for obtaining and measuring wing patterns are provided in Lucas *et al*. (2018), but in brief, the ventral surface of hind wings were dissected and photographed under a Leica light microscope; the size (area) and position of wing pattern elements were then measured using ImageJ (Abramoff *et al*., 2004) (here, we use only the size data). Element sizes were standardized by wing size and differences between males and females. We conducted the PCA in R using the prcomp function after centering and standardizing the wing pattern measurements.

For wing-pattern genetic mapping, we reanalyzed the same 17 wing pattern measurements and corresponding genetic data from three *Lycaeides* populations: YG (*L. anna*, *n* = 100), GNP (*L. idas*, *n* = 98), and SIN (*L. melissa*, *n* = 97) (Lucas *et al*., 2018). We aligned genotyping-by-sequencing (GBS) reads from these butterflies to the *L. melissa* reference genome (Zhang *et al*., 2023) using the aln and samse algorithms implemented in bwa (version 0.7.17-r1198) (Li & Durbin, 2009). For alignment, we allowed a maximum of five mismatches per read, no more than two mismatches in the first 20-bp seed, set the quality threshold for read trimming to 10, and retained only reads with a unique best alignment. We then used samtools (version 1.16) to compress, sort, and index the alignments (Li *et al*., 2009). SNPs were identified using bcftools mpileup and call (Li, 2011). We considered only bases and reads with minimum base and mapping quality scores of 20 and 30, respectively, ignored indels, and retained SNPs only when the probability that the site was invariant given the data was less than 0.01, using the default prior probability of 0.001 that a nucleotide site was variable. We further filtered the SNP set to retain only biallelic variants with a mean coverage of at least 2× per butterfly, support from at least 10 reads, a quality score of at least 30, base-quality, mapping-quality, and read-position bias statistics with absolute values less than four, missing data for no more than 20% of individuals, sequence coverage not exceeding three standard deviations above the mean (to reduce the inclusion of paralogous loci), and a minor allele frequency greater than 0.001 (which primarily removes singleton variants that are uninformative for genetic mapping). These filters resulted in a final dataset of 137,865 SNPs.

We next estimated genotypes for each individual at each locus. To do so, we first obtained maximum-likelihood estimates of allele frequencies for each population using the expectation-maximization algorithm described by Li (2011) and implemented in estpEM (version 0.1) (Soria-Carrasco *et al*., 2014). This method accounts for genotype uncertainty, as represented by genotype likelihoods, when estimating allele frequencies. The version of the software we used also accommodates the hemizygous state of females for Z-linked loci. We set the convergence tolerance to 0.001 and used 40 independent starting values. We then obtained empirical Bayesian point estimates of individual genotypes by combining the estimated population-specific allele frequencies (used as priors) with genotype likelihoods from bcftools. The maximum posterior genotype was used as the point estimate and coded as 0, 1, or 2 copies of the non-reference allele. For Z-linked SNPs in females, genotypes were coded as 0 or 2.

We then used a polygenic genome-wide association mapping approach to estimate the proportion of additive genetic variance explained by each chromosome for each trait in each population (Zhou *et al*., 2013). Specifically, we fit Bayesian sparse linear mixed models using gemma (version 0.95a) (Zhou & Stephens, 2012). For each trait-population combination, we ran 20 independent Markov chain Monte Carlo chains of 1,000,000 iterations, including a burn-in period of 500,000 iterations. We then estimated additive genetic variance from model-averaged SNP effect estimates and population allele frequencies (*p*) as *V_a_* = 2*p*(1 − *p*)*β̂*^2^, where *β̂* is the posterior mean SNP effect estimated by BSLMM, including contributions from both the polygenic and sparse effect components (see, e.g., Lucas *et al*., 2018; Gompert *et al*., 2019).

## Supplemental Tables and Figures

**Table S1:** Summary of statistical tests concerning the symmetry of unnormalized weights for the two less common topologies and phylogenetic signal for each window from each four-taxon, windowed **CASTER** analysis. Results are shown for the four-taxon sets from Figures 4 and 5. For this analysis, we defined the primary (species-tree) topology as the topology with the highest mean score across all windows, and the less common topologies as the two alternatives. The statistics reported are: Σ*_Z_* = the mean sum of weights for Z chromosome windows; Σ*_A_* = the mean sum of weights for autosomal windows; *P*_Σ_ = the *P*-value for the alternative hypothesis that Σ*_Z_ >* Σ*_A_*; Δ*_Z_* = the mean difference in weights between the two less common topologies for Z chromosome windows; Δ*_A_* = the mean difference in weights between the two less common topologies for autosomal windows; and *P*_Δ_ = the *P*-value for the alternative hypothesis that Δ*_Z_* ≠ Δ*_A_*. Permutation tests (1000 permutations) of datasets thinned to minimize the effects of non-independence between adjacent windows were used to evaluate these hypotheses.

| Taxon set | $\Sigma_A$ | $\Sigma_Z$ | $P_\Sigma$ | $\Delta_A$ | $\Delta_Z$ | $P_\Delta$ |
| --- | --- | --- | --- | --- | --- | --- |
| ((CLH,TIC),BHP) | 0.47 | 0.78 | 0.001 | 0.08 | 0.49 | 0.002 |
| ((CLH,YG),BHP) | 0.60 | 0.51 | 0.956 | 0.04 | 0.13 | 0.002 |
| ((TIC,YG),BHP) | 0.49 | 0.46 | 0.731 | 0.06 | 0.14 | 0.020 |
| ((CP,TIC),YG) | 0.45 | 0.69 | 0.001 | 0.09 | 0.02 | 0.004 |
| ((MR,YG),CP) | 0.61 | 0.76 | 0.001 | 0.11 | 0.36 | 0.002 |
| ((CP,YG),BHP) | 0.46 | 0.45 | 0.665 | 0.15 | 0.16 | 0.380 |
| ((EP,SHC),BHP) | 0.37 | 0.41 | 0.243 | 0.09 | 0.21 | 0.032 |
| ((BTB,SIN),GNP) | 0.30 | 0.35 | 0.031 | 0.01 | 0.04 | 0.028 |
| ((BCR,SIN),GNP) | 0.30 | 0.35 | 0.027 | 0.01 | 0.04 | 0.078 |
| ((HNV,SIN),GNP) | 0.21 | 0.25 | 0.042 | 0.00 | 0.09 | 0.002 |
| ((ABM,SIN),TBY) | 0.55 | 0.86 | 0.001 | 0.00 | -0.01 | 0.588 |
| ((BHP,SIN),YG) | 0.62 | 1.00 | 0.001 | 0.02 | 0.01 | 0.380 |
| ((BCR,BTB),YG) | 0.38 | 0.33 | 0.920 | 0.00 | -0.00 | 0.924 |
| ((LS,YG),SIN) | 1.17 | 1.57 | 0.001 | 0.00 | 0.00 | 0.690 |

**Table S2:**
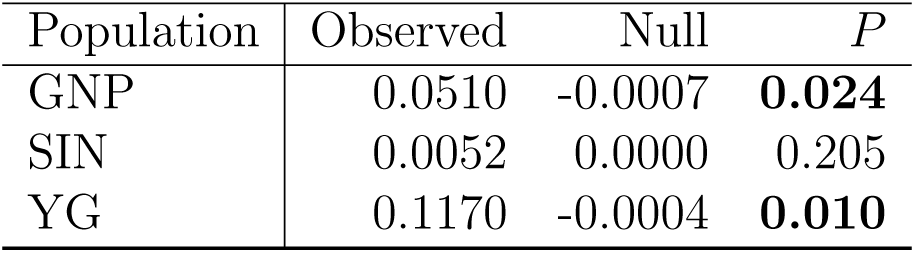
Summary of tests of an excess contribution of the Z chromosome to wing pattern traits. Tests are based on the residuals from regressing the proportion of variation explained by each chromosome for each trait on the number of SNPs per chromosome. The observed value for each population is then the mean of the residuals for the Z across traits. Null expectations were then obtained by permuting residuals across chromosomes and computing the mean of the residuals for the permuted residuals assigned to the Z (1000 permutations were performed). We report the mean of the null (“Null”) and the permutation *P*-value, that is the proportion of null simulations with test statistics equal to or exceeding the observed value. Significant *P*-values at *α* = 0.05 are in bold.

**Figure S1:**
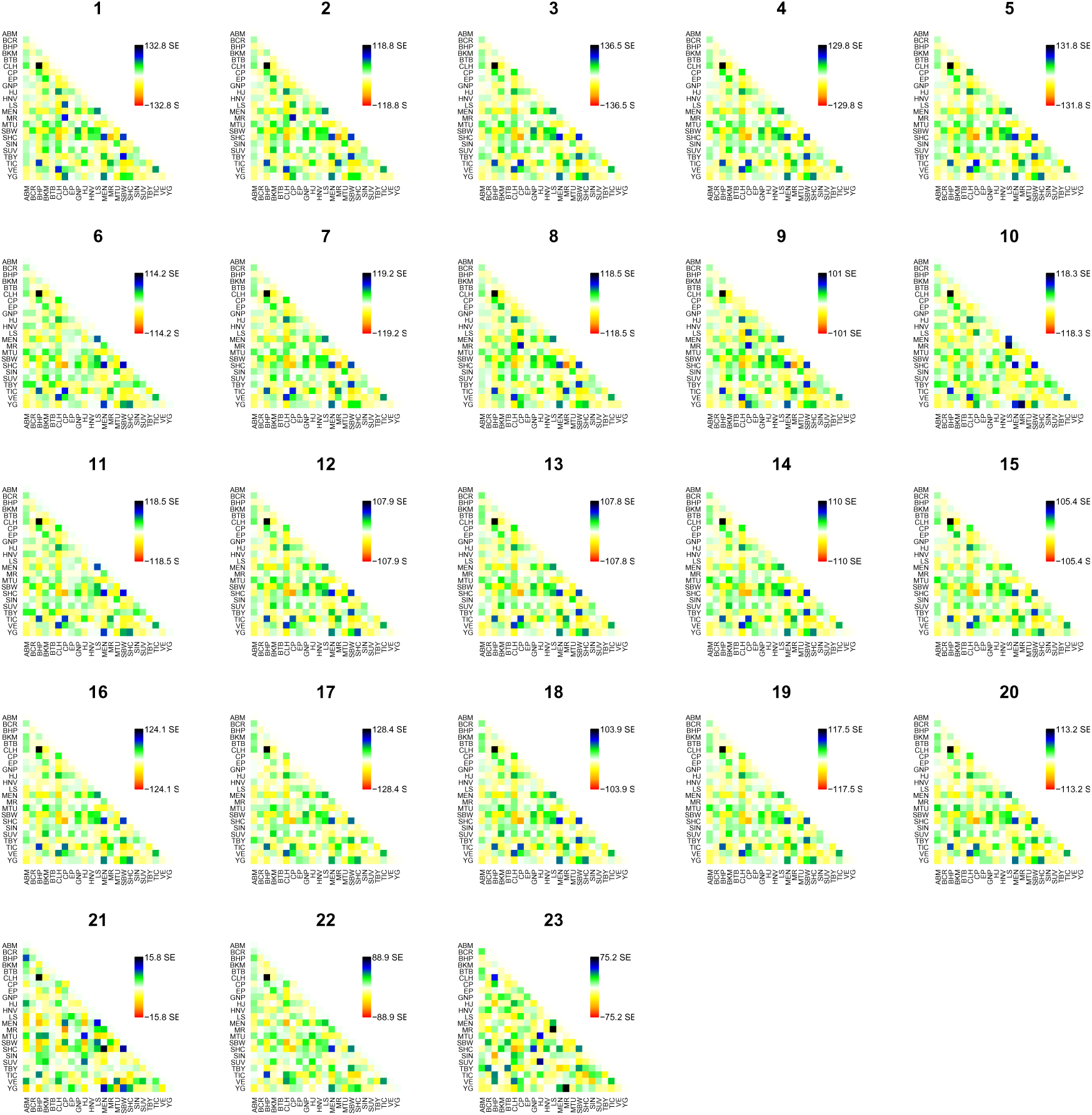
Residuals from population trees with no migration edges. Each panel shows residuals for one of the 23 chromosomes (indicated by the number above each plot). In each plot, colored boxes denote residuals for population pairs. Residuals have been scaled by the standard error across all pairs. Positive values denote pairs that are more closely related than expected from the tree, which is suggestive of admixture events.

**Figure S2:**
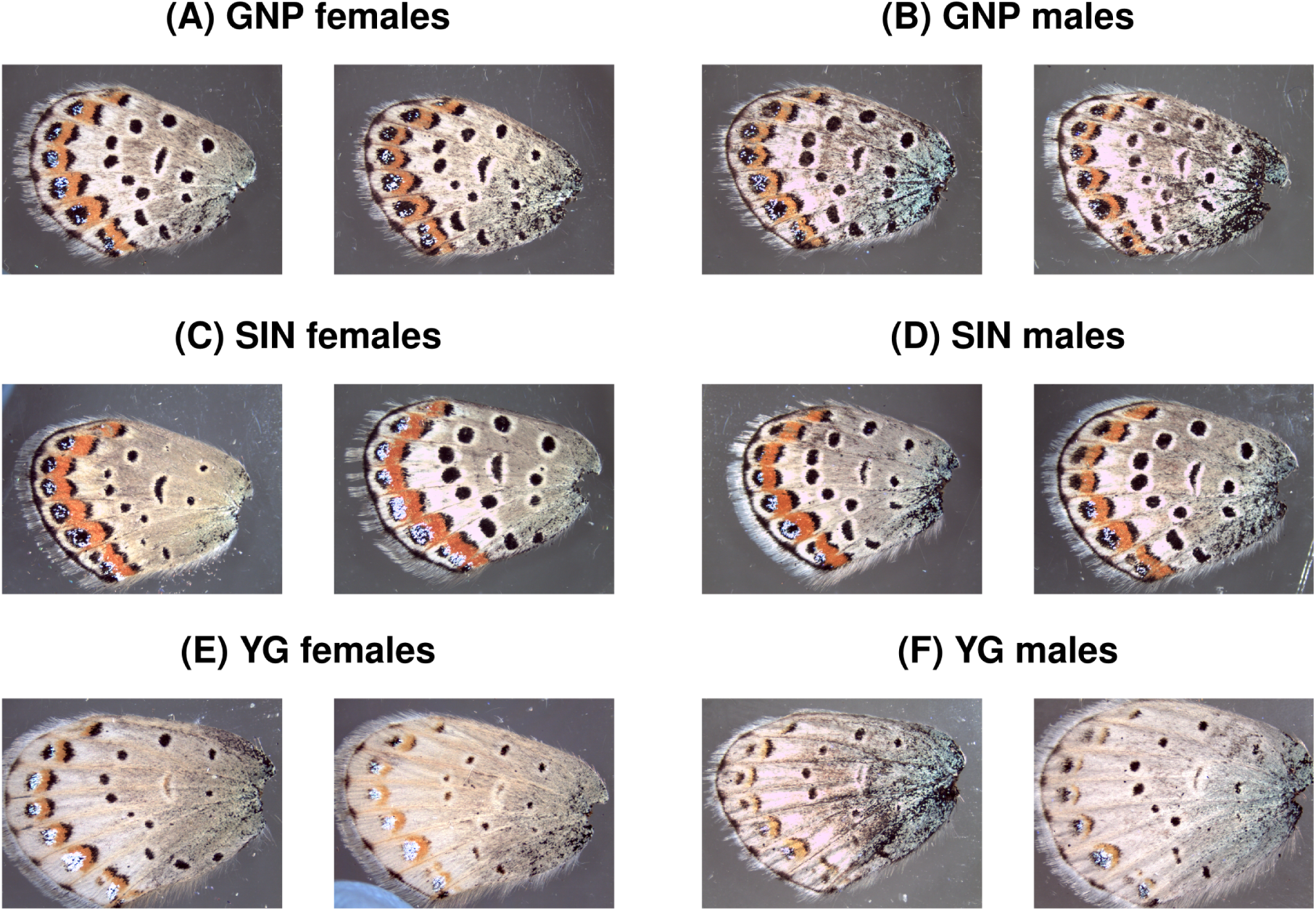
Example images of ventral hindwing surface of 12 *Lycaeides* butterflies. Two images each are shown for females and males from GNP (panels A and B, *L. idas*), SIN (panels C and D, *L. melissa*) and YG (panels E and F, *L. anna*). All images were taken at the same scale.

**Figure S3:**
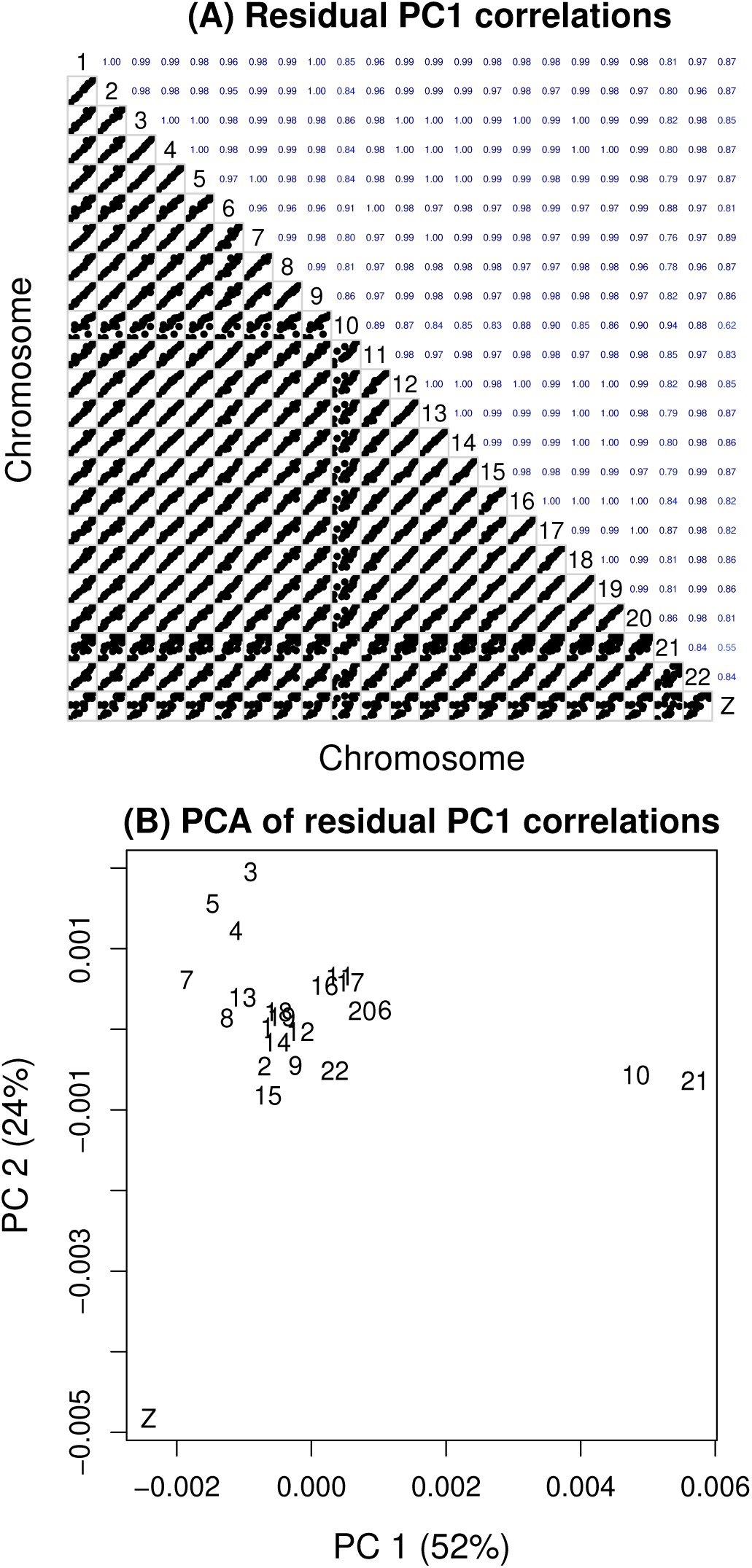
Comparison of residual structure across chromosomes. Residuals from the fit of a bifurcating tree (no migration edges) for each chromosome were first summarized using PCA. Panel (A) shows pairwise correlations for PC1 scores (across populations) for each pair of chromosomes, with scatterplots on the lower triangle, Pearson correlation coefficients on the upper triangle and chromosome numbers along the diagonal. In panel (B), results of a second PCA of the residual PC1 for each chromosome are shown, which effectively summarizes the information in panel (A) in a two-dimensional space. The resulting scatterplot shows PC1 and PC2 scores from this PCA for each chromosome (numbered autosomes and the Z in the plot). The percent variance explained by each PC is given.

**Figure S4:**
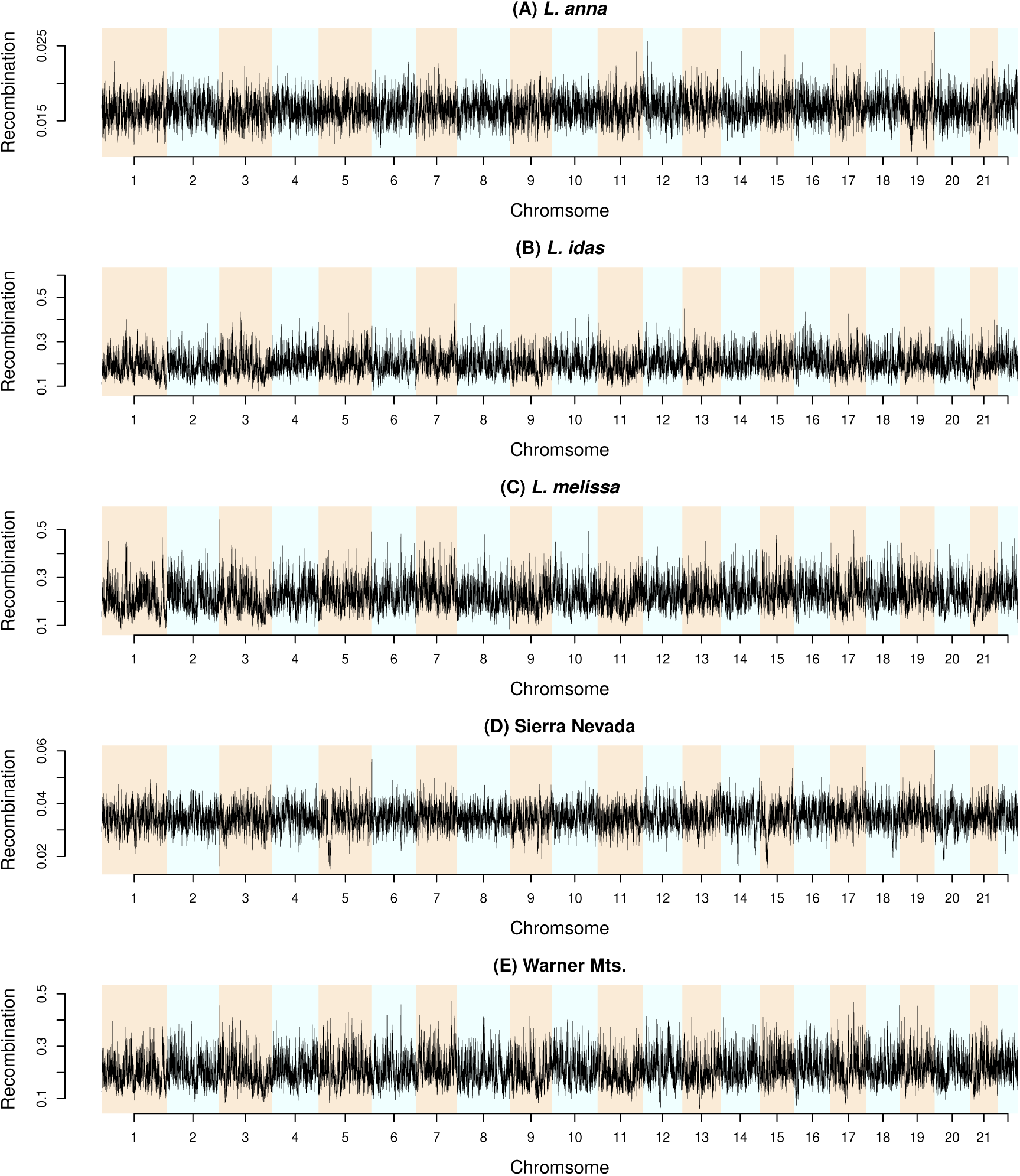
Recombination-rate landscapes for five *Lycaeides* lineages. Estimated population recombination rates (*ρ* = 4*N_e_r*) are shown across the 22 autosomes for each lineage. For each 25-kbp window, we calculated a composite recombination-rate estimate as the mean of the estimates for the corresponding 25-kbp, 250-kbp, and 1-Mbp regions containing that window.

**Figure S5:**
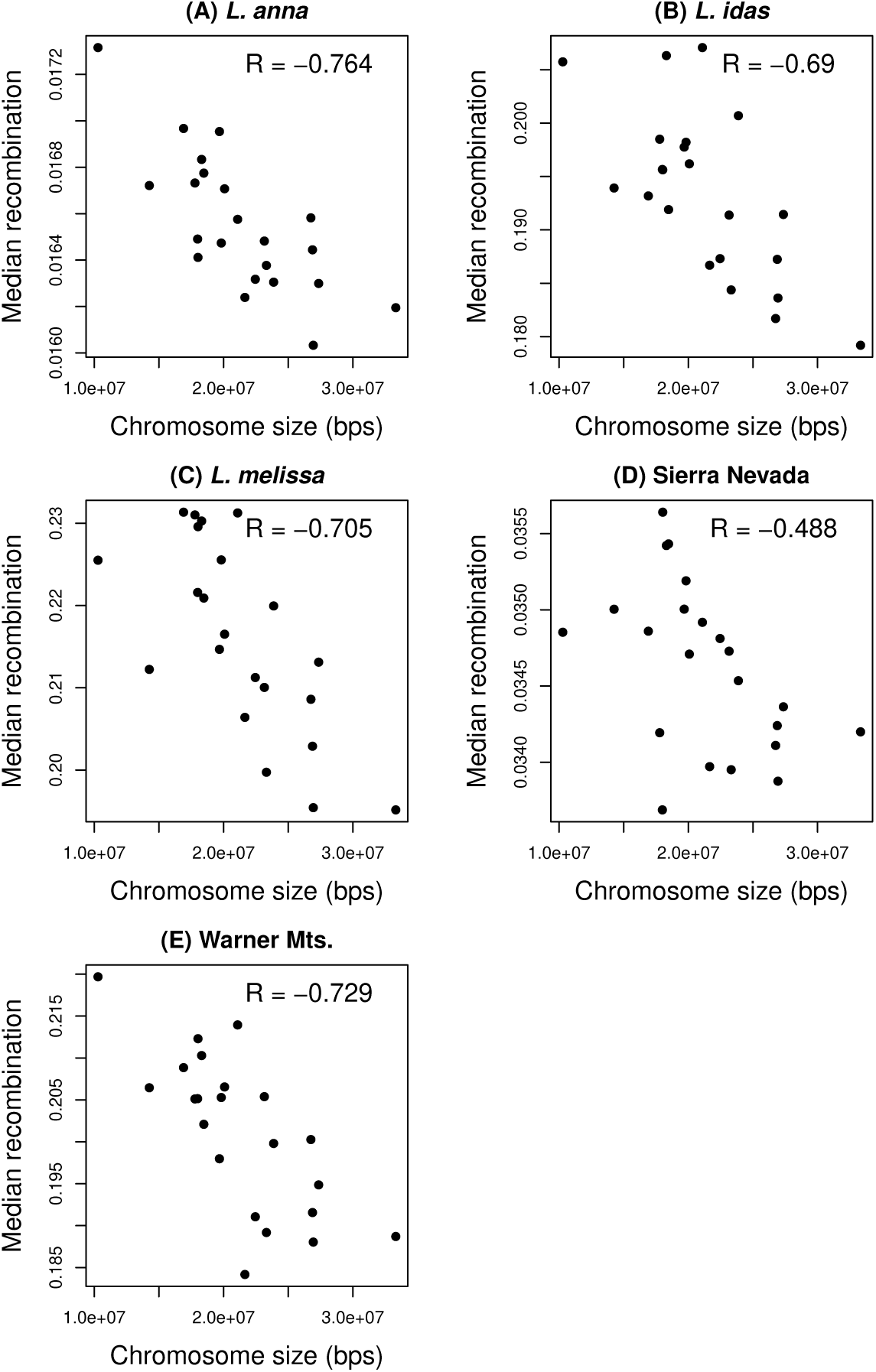
Relationship between chromosome size and mean recombination rate across five *Lycaeides* lineages. Scatterplots show the relationship between chromosome size and mean recombination rate (*ρ* = 4*N_e_r*) for each of the 22 autosomes in each of five *Lycaeides* lineages. Each point represents a chromosome. Pearson correlation coefficients for the relationship between chromosome size and recombination rate are reported in each panel.

**Figure S6:**
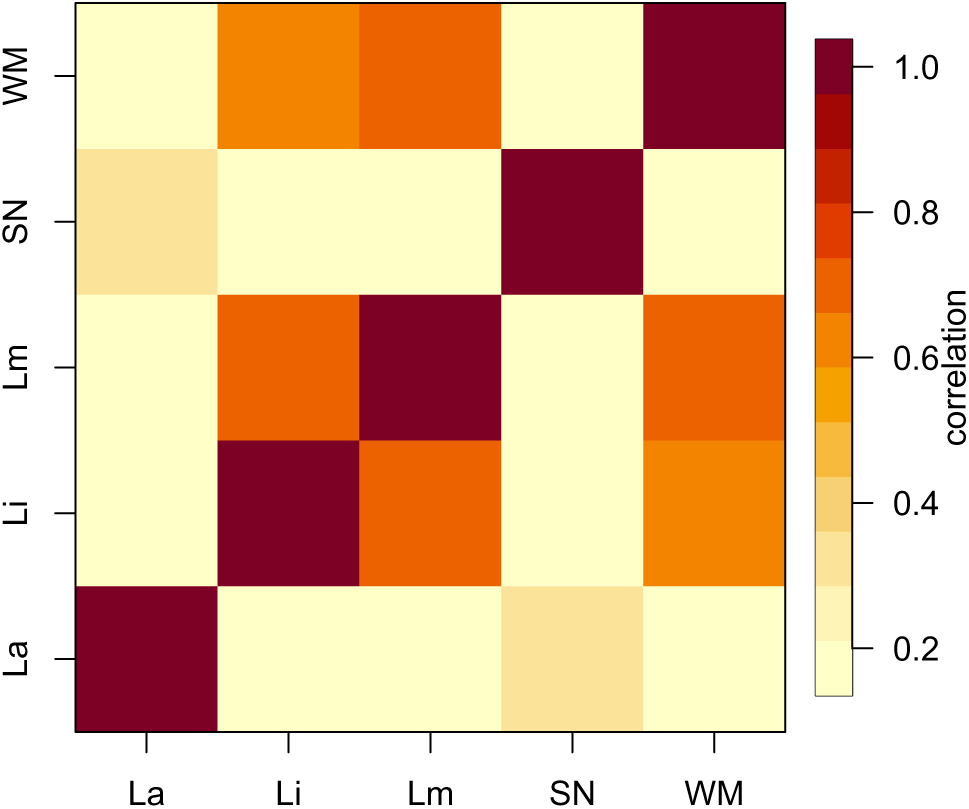
Relationships among recombination landscapes across five *Lycaeides* lineages. The heatmap summarizes pairwise correlations in recombination-rate landscapes between each pair of *Lycaeides* lineages. The scale bar indicates the strength of the Pearson correlation. Abbreviations are as follows: La = *L. anna*, Li = *L. idas*, Lm = *L. melissa*, SN = Sierra Nevada, and WM = Warner Mountain.

**Figure S7:**
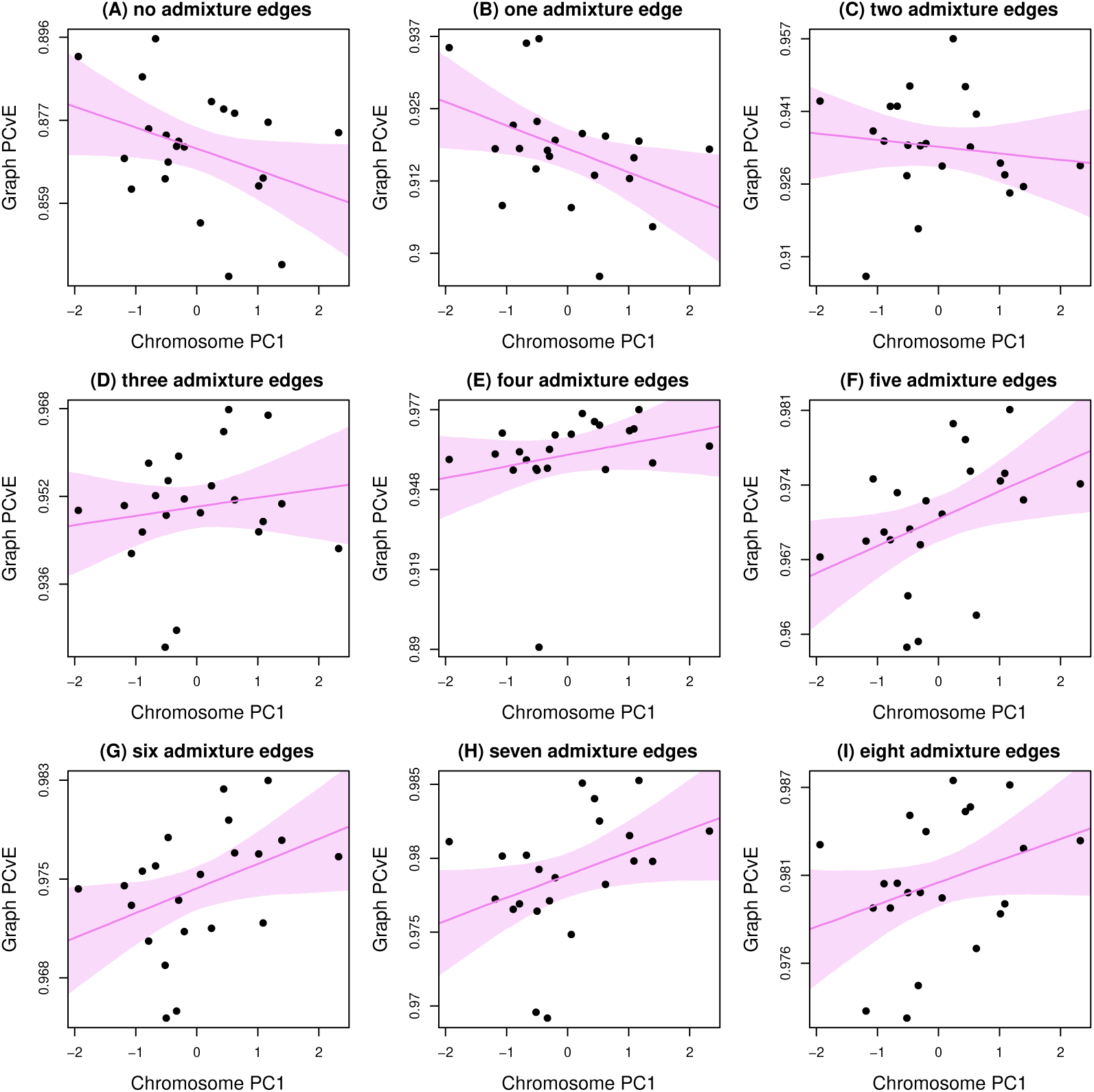
Scatterplots depict the relationships between chromosomal features and the proportion of allele frequency covariance explained (PCvE) for graphs with different numbers of admixture edges. Only the 22 autosomes were included, each denoted by a black point. Three chromosome features–size, mean recombination rate, and gene density–were summarized using PCA for these regression analyses; larger positive values of PC1 (which captured 69% of the variation in these features) denote larger chromosomes with lower recombination rates per bp and higher gene density. In each panel, the solid line and shaded region show the fitted regression line and 90% equal-tail probability interval (ETPI) from a Bayesian regression model. Positive slopes indicate higher PCvE for chromosomes that are larger, experience less recombination, and have higher gene density (and vice versa for negative slopes). Slope estimates were: (A) no admixture edges, *β* = −0.36, 95% ETPI = −0.71 – 0.00, posterior probability *β >* 0 (pp) = 0.05; (B) one admixture edge, *β* = −0.42, 95% ETPI = −0.78 – −0.05, pp = 0.03; (C) two admixture edges, *β* = −0.13, 95% ETPI = −0.52 – 0.26, pp = 0.29; (D) three admixture edges, *β* = 0.16, 95% ETPI = −0.24 – 0.53, pp = 0.75; (E) four admixture edges, *β* = 0.24, 95% ETPI = −0.15 – 0.62, pp = 0.85; (F) five admixture edges, *β* = 0.42, 95% ETPI = 0.07 – 0.79, pp = 0.97; (G) six admixture edges, *β* = 0.40, 95% ETPI = 0.02 – 0.77, pp = 0.96; (H) seven admixture edges, *β* = 0.37, 95% ETPI = 0.01 – 0.75, pp = 0.95; (I) eight admixture edges, *β* = 0.33, 95% ETPI = −0.05 – 0.71, pp = 0.92.

**Figure S8:**
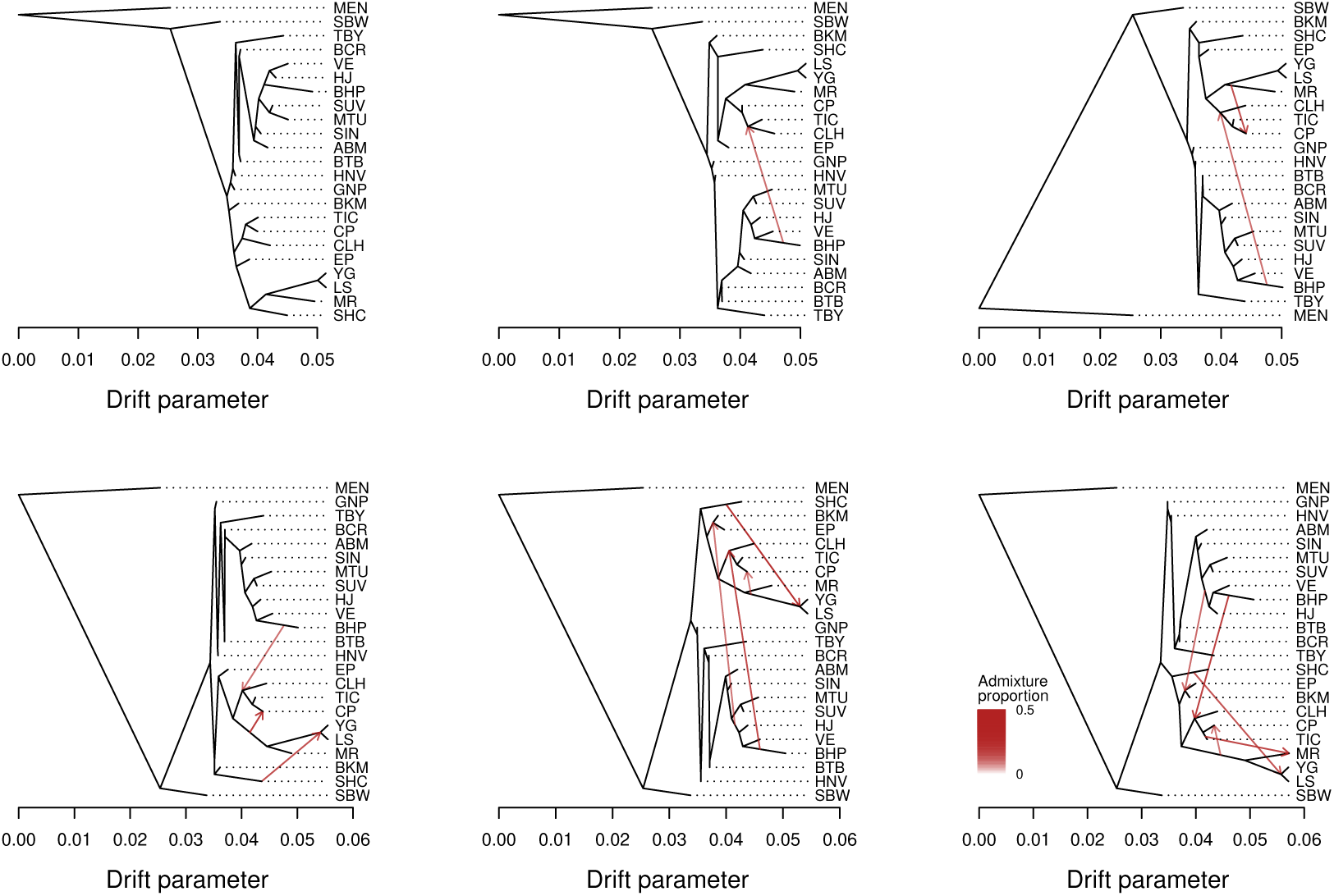
Population graphs for chromosome 1, with units in terms of a drift parameter proportional to evolutionary change and with the *L. argyrognomon* population (MEN) included as an outgroup. Graphs are shown for m = 0 (bifurcating tree) to five migration edges (red arrows). The admixture proportions associated with each are indicated by the intensity of each red arrow.

**Figure S9:**
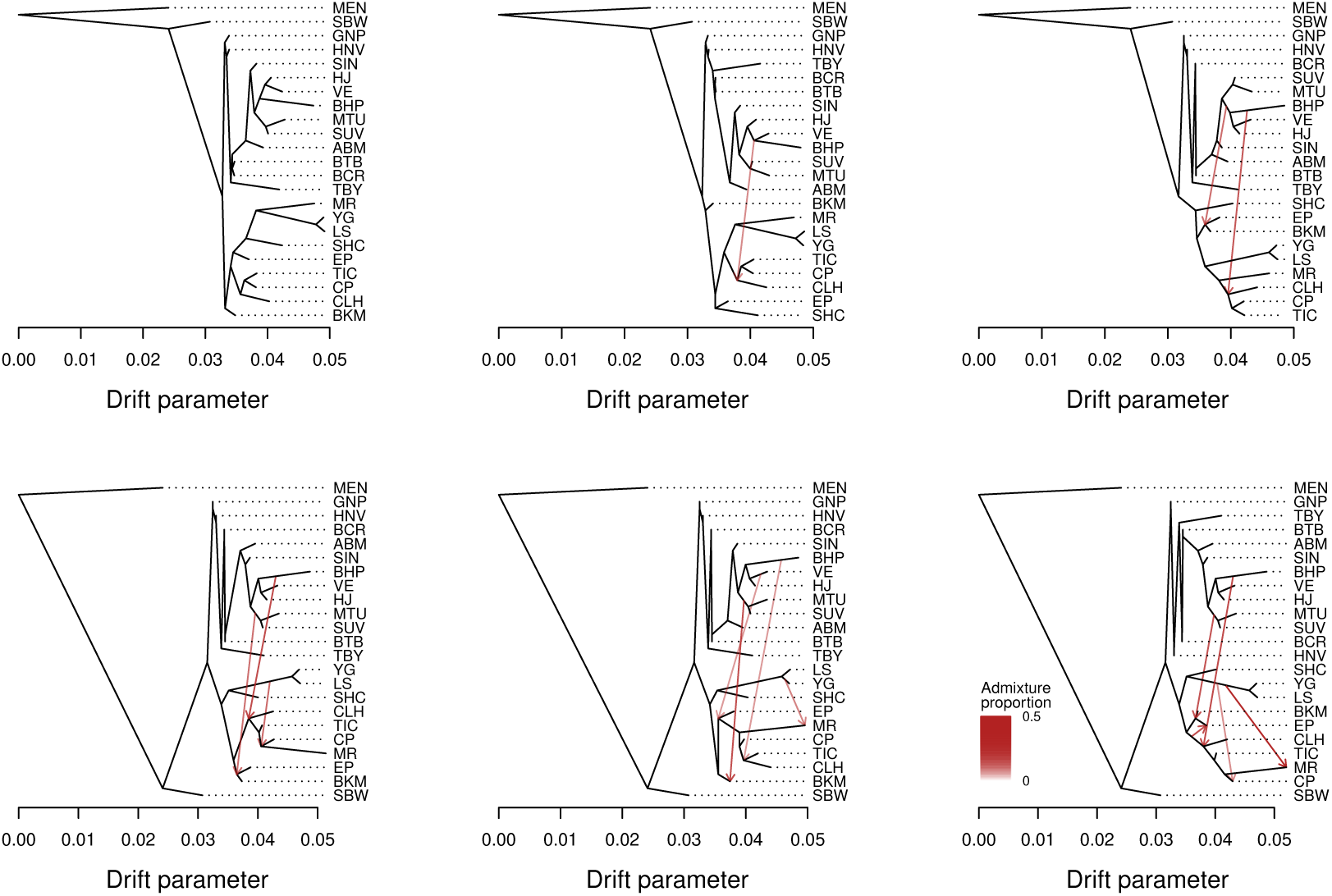
Population graphs for chromosome 2, with units in terms of a drift parameter proportional to evolutionary change and with the *L. argyrognomon* population (MEN) included as an outgroup. Graphs are shown for m = 0 (bifurcating tree) to five migration edges (red arrows). The admixture proportions associated with each are indicated by the intensity of each red arrow.

**Figure S10:**
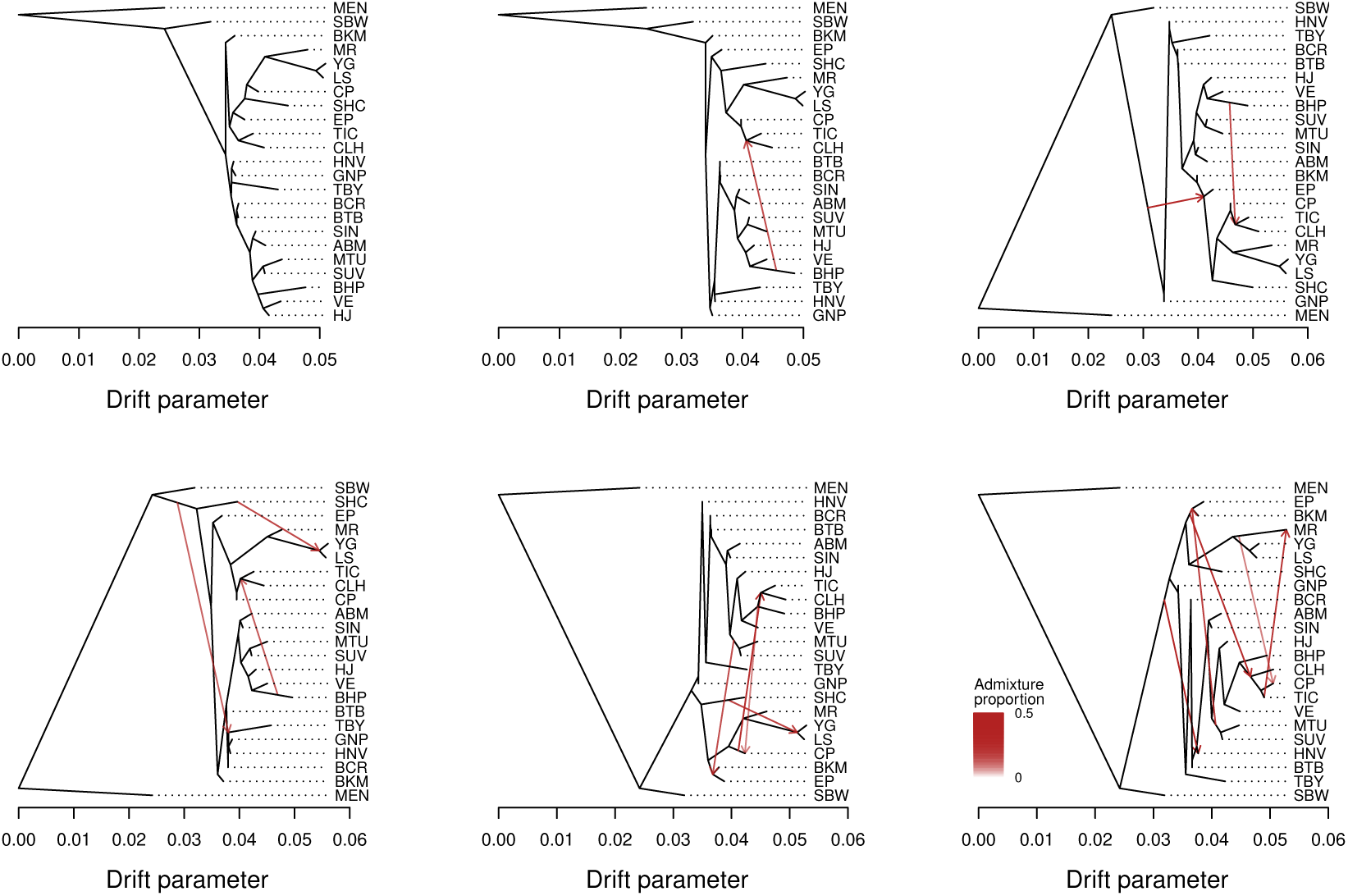
Population graphs for chromosome 3, with units in terms of a drift parameter proportional to evolutionary change and with the *L. argyrognomon* population (MEN) included as an outgroup. Graphs are shown for m = 0 (bifurcating tree) to five migration edges (red arrows). The admixture proportions associated with each are indicated by the intensity of each red arrow.

**Figure S11:**
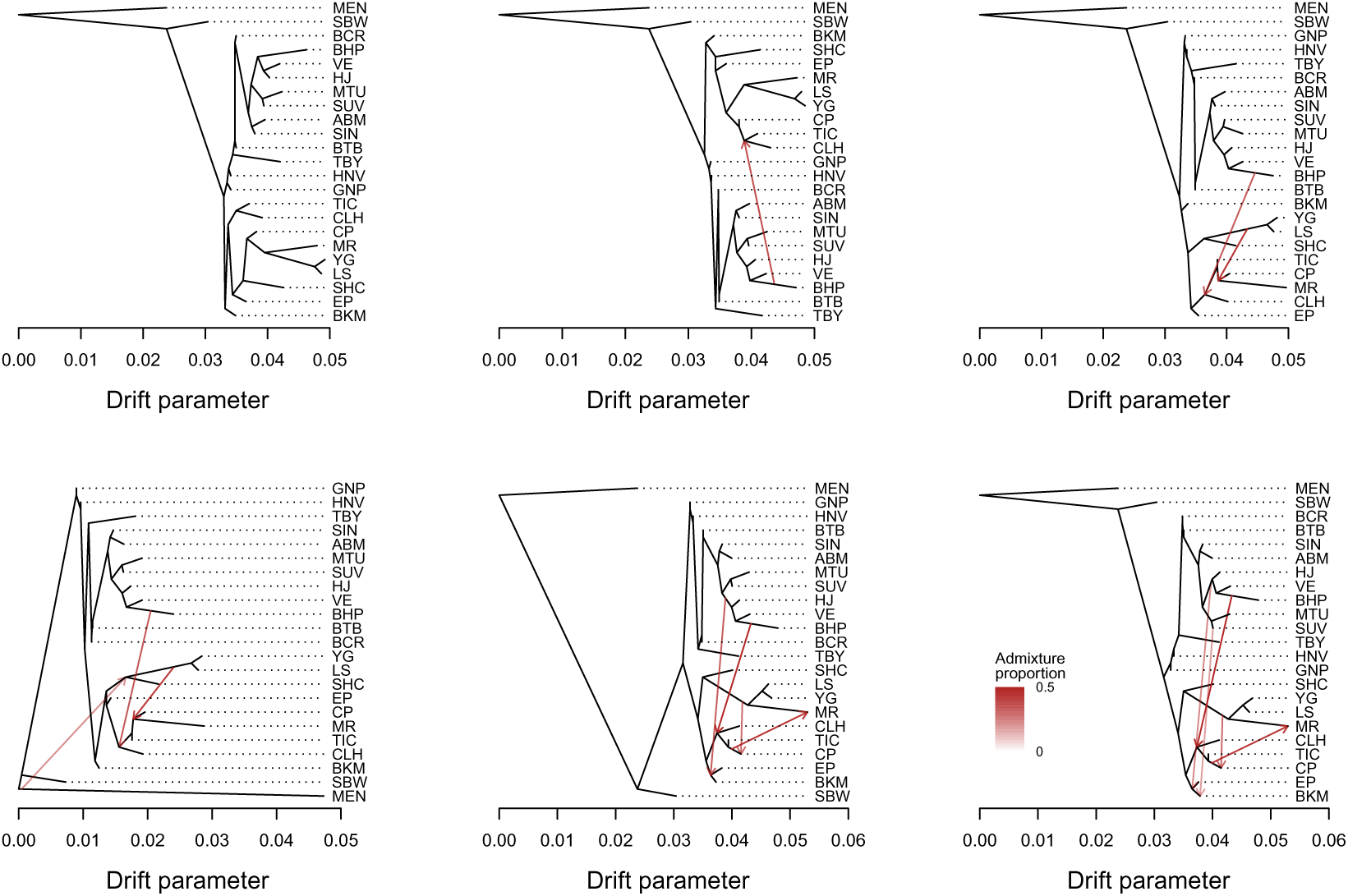
Population graphs for chromosome 4, with units in terms of a drift parameter proportional to evolutionary change and with the *L. argyrognomon* population (MEN) included as an outgroup. Graphs are shown for m = 0 (bifurcating tree) to five migration edges (red arrows). The admixture proportions associated with each are indicated by the intensity of each red arrow.

**Figure S12:**
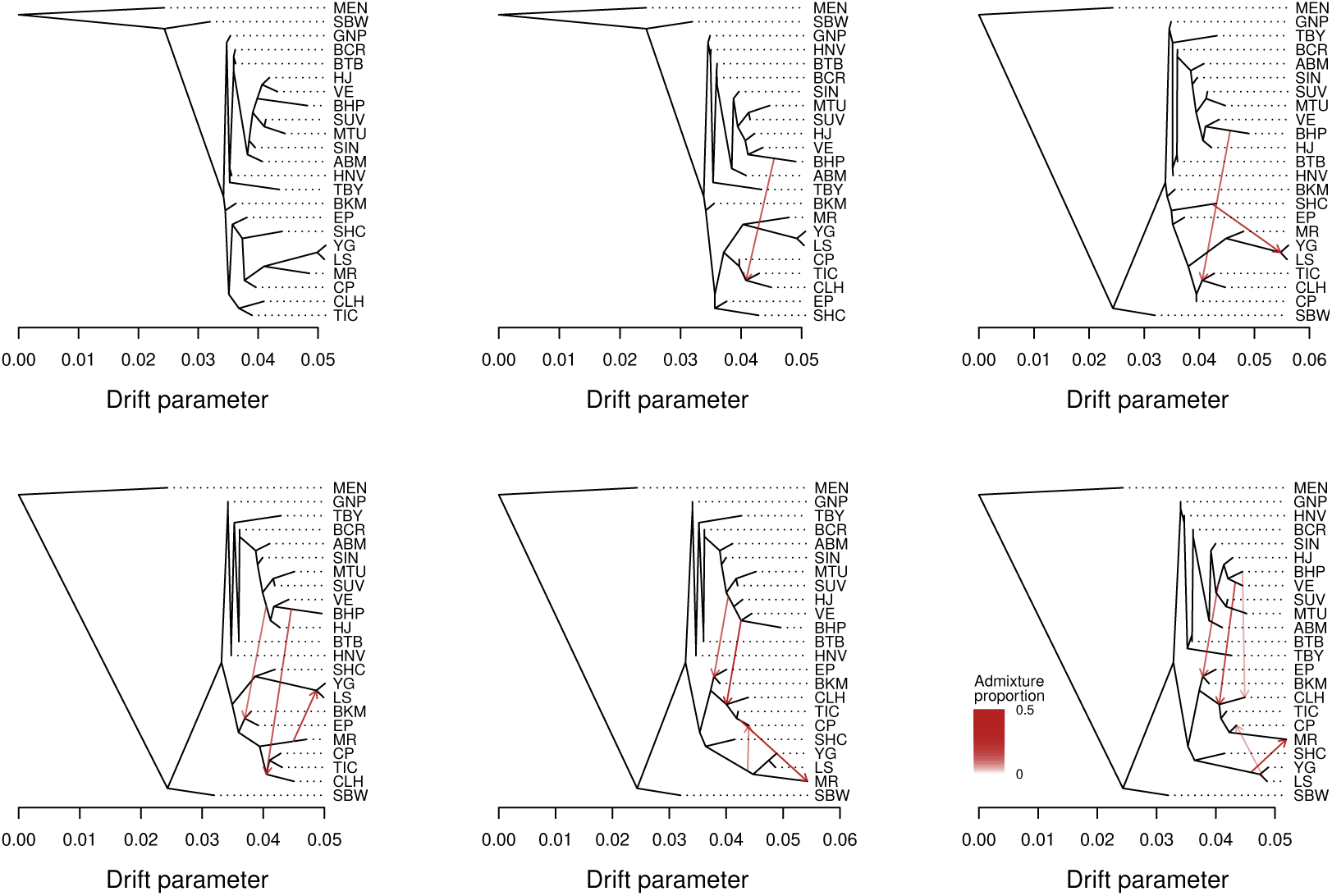
Population graphs for chromosome 5, with units in terms of a drift parameter proportional to evolutionary change and with the *L. argyrognomon* population (MEN) included as an outgroup. Graphs are shown for m = 0 (bifurcating tree) to five migration edges (red arrows). The admixture proportions associated with each are indicated by the intensity of each red arrow.

**Figure S13:**
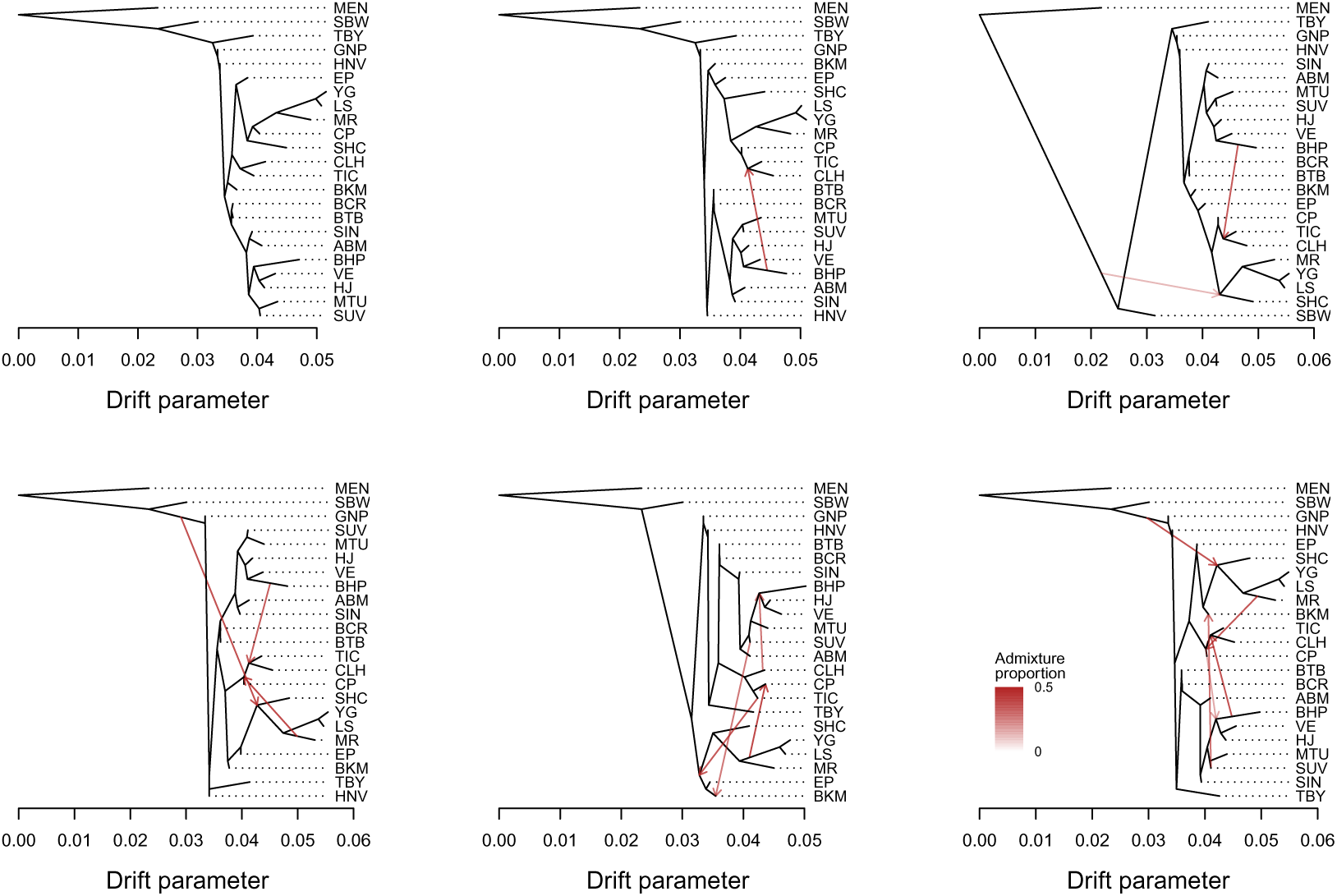
Population graphs for chromosome 6, with units in terms of a drift parameter proportional to evolutionary change and with the *L. argyrognomon* population (MEN) included as an outgroup. Graphs are shown for m = 0 (bifurcating tree) to five migration edges (red arrows). The admixture proportions associated with each are indicated by the intensity of each red arrow.

**Figure S14:**
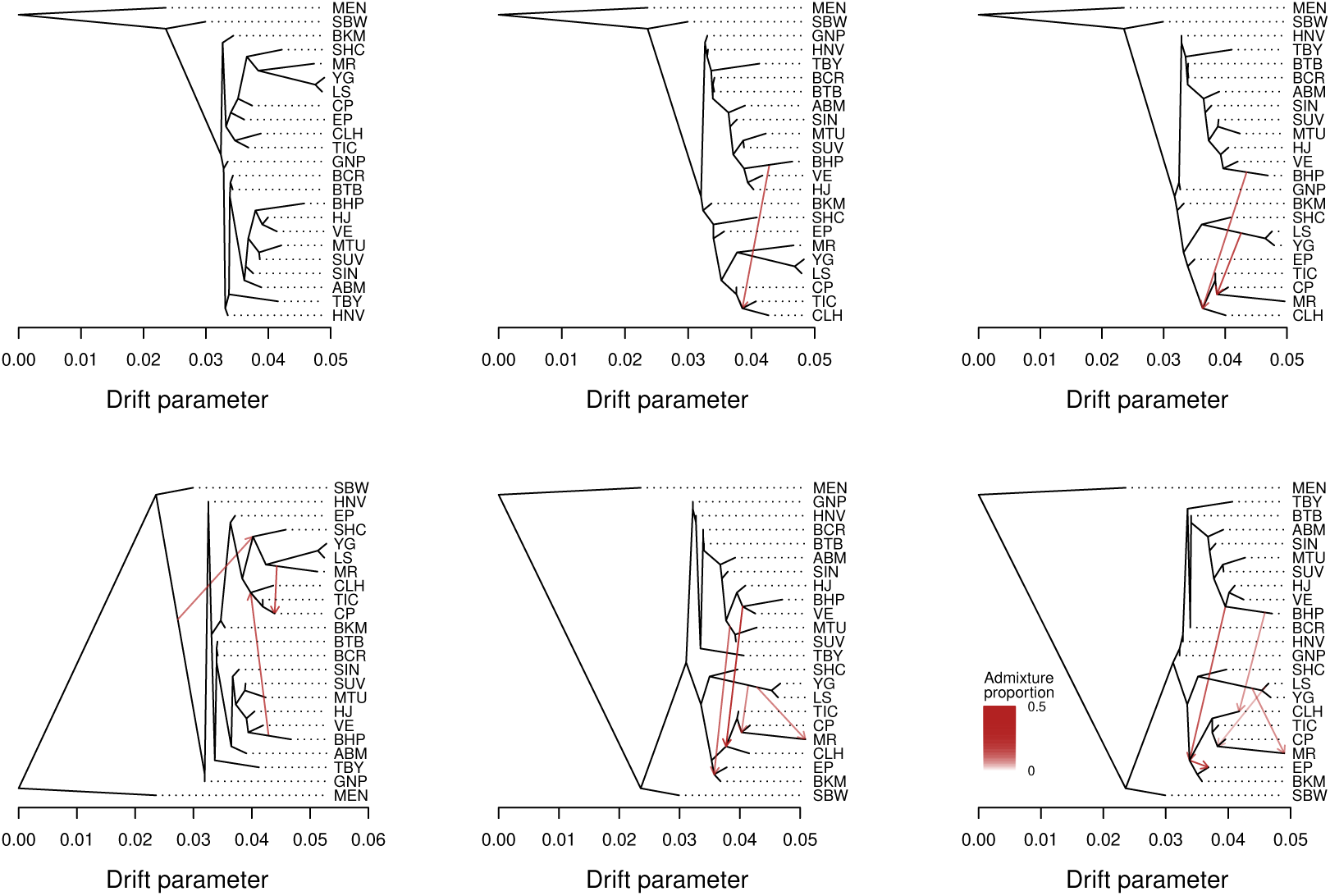
Population graphs for chromosome 7, with units in terms of a drift parameter proportional to evolutionary change and with the *L. argyrognomon* population (MEN) included as an outgroup. Graphs are shown for m = 0 (bifurcating tree) to five migration edges (red arrows). The admixture proportions associated with each are indicated by the intensity of each red arrow.

**Figure S15:**
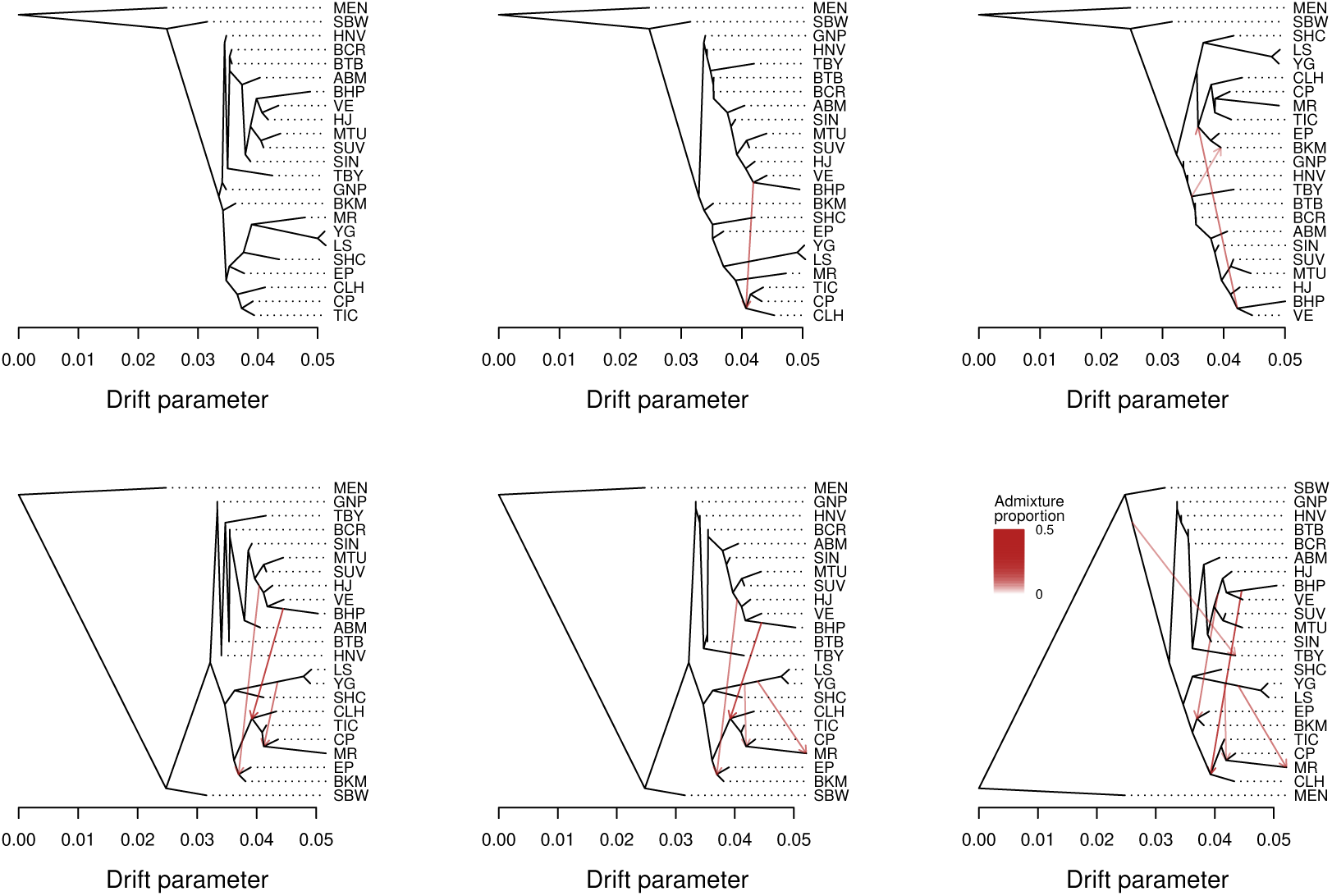
Population graphs for chromosome 8, with units in terms of a drift parameter proportional to evolutionary change and with the *L. argyrognomon* population (MEN) included as an outgroup. Graphs are shown for m = 0 (bifurcating tree) to five migration edges (red arrows). The admixture proportions associated with each are indicated by the intensity of each red arrow.

**Figure S16:**
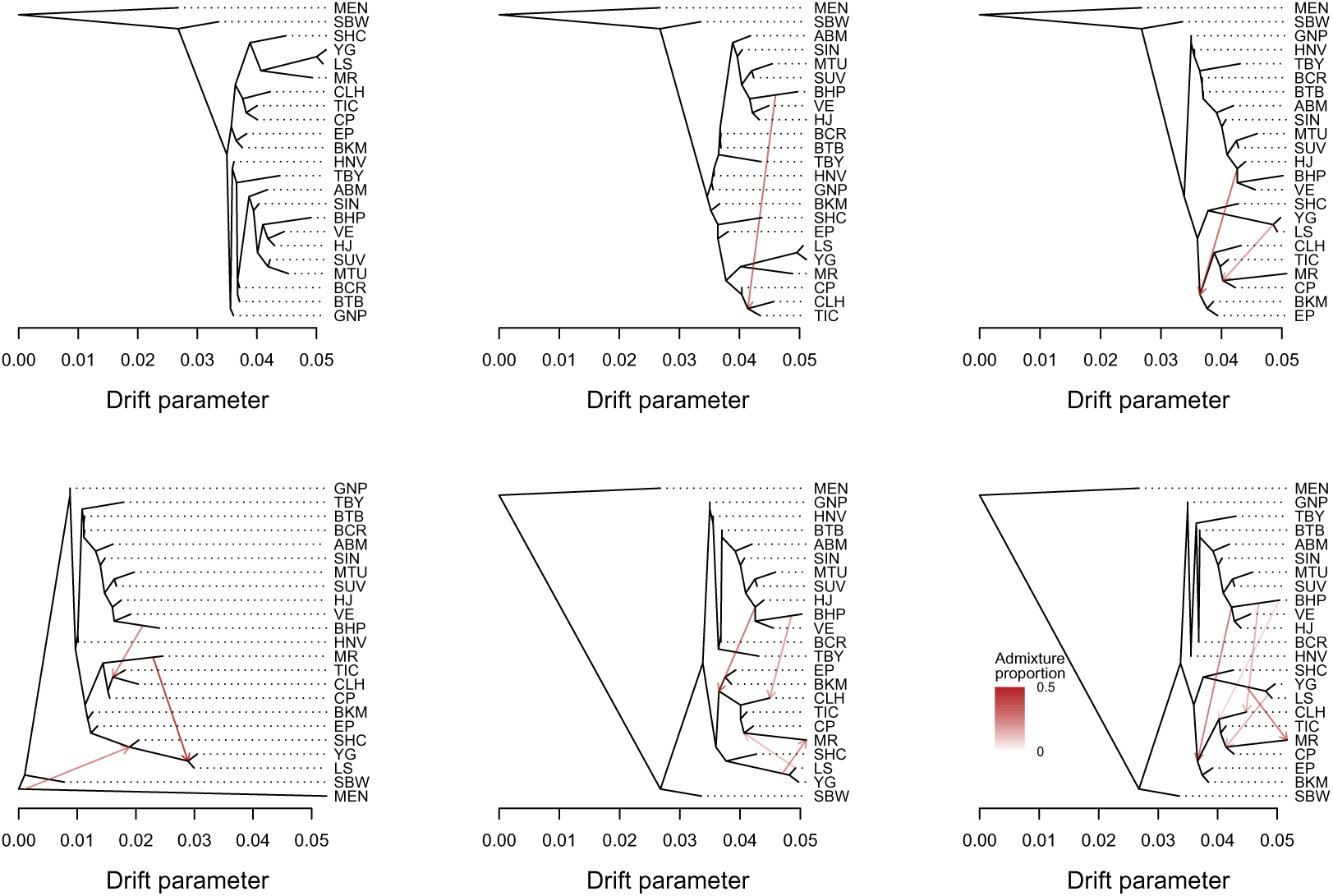
Population graphs for chromosome 9, with units in terms of a drift parameter proportional to evolutionary change and with the *L. argyrognomon* population (MEN) included as an outgroup. Graphs are shown for m = 0 (bifurcating tree) to five migration edges (red arrows). The admixture proportions associated with each are indicated by the intensity of each red arrow.

**Figure S17:**
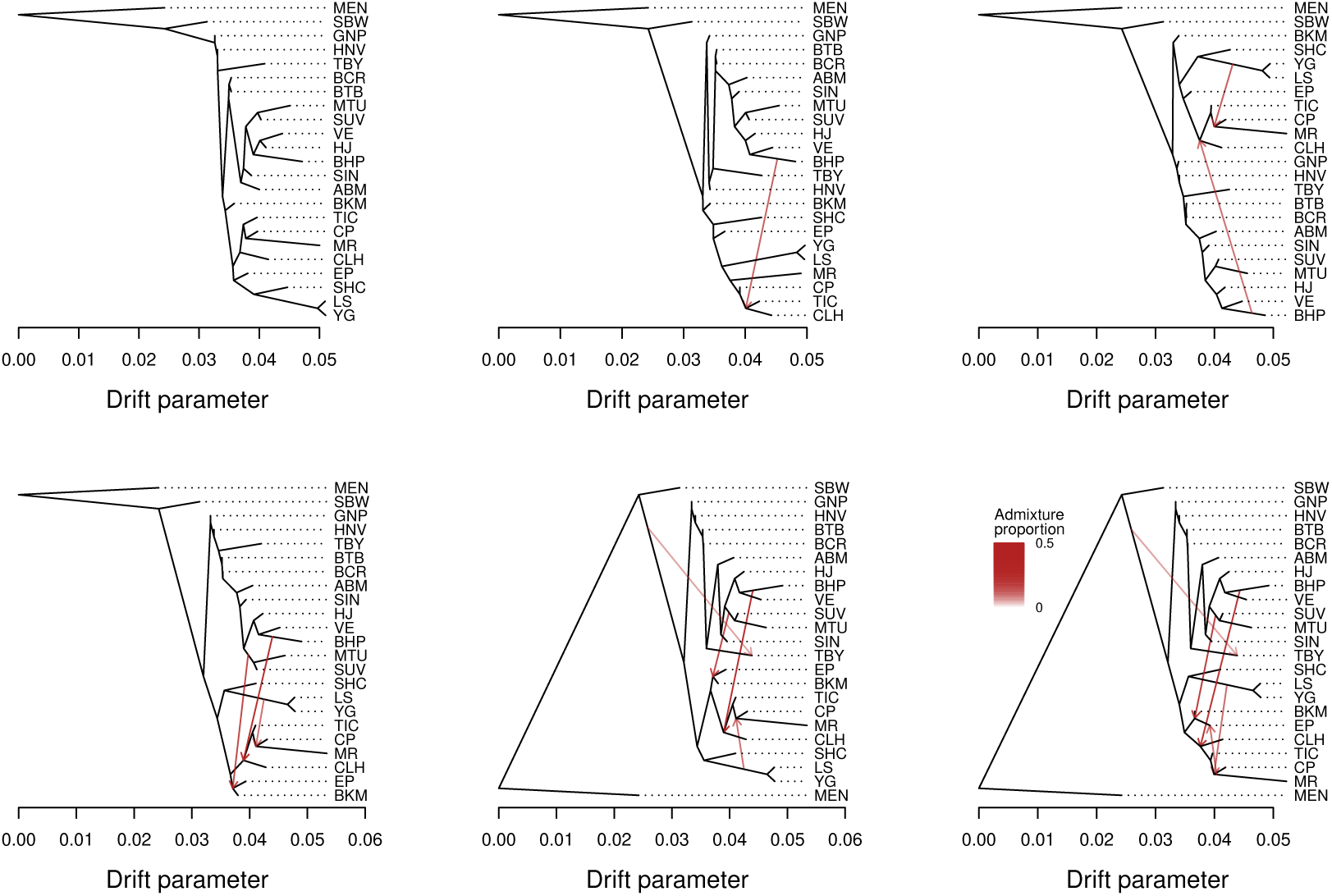
Population graphs for chromosome 10, with units in terms of a drift parameter proportional to evolutionary change and with the *L. argyrognomon* population (MEN) included as an outgroup. Graphs are shown for m = 0 (bifurcating tree) to five migration edges (red arrows). The admixture proportions associated with each are indicated by the intensity of each red arrow.

**Figure S18:**
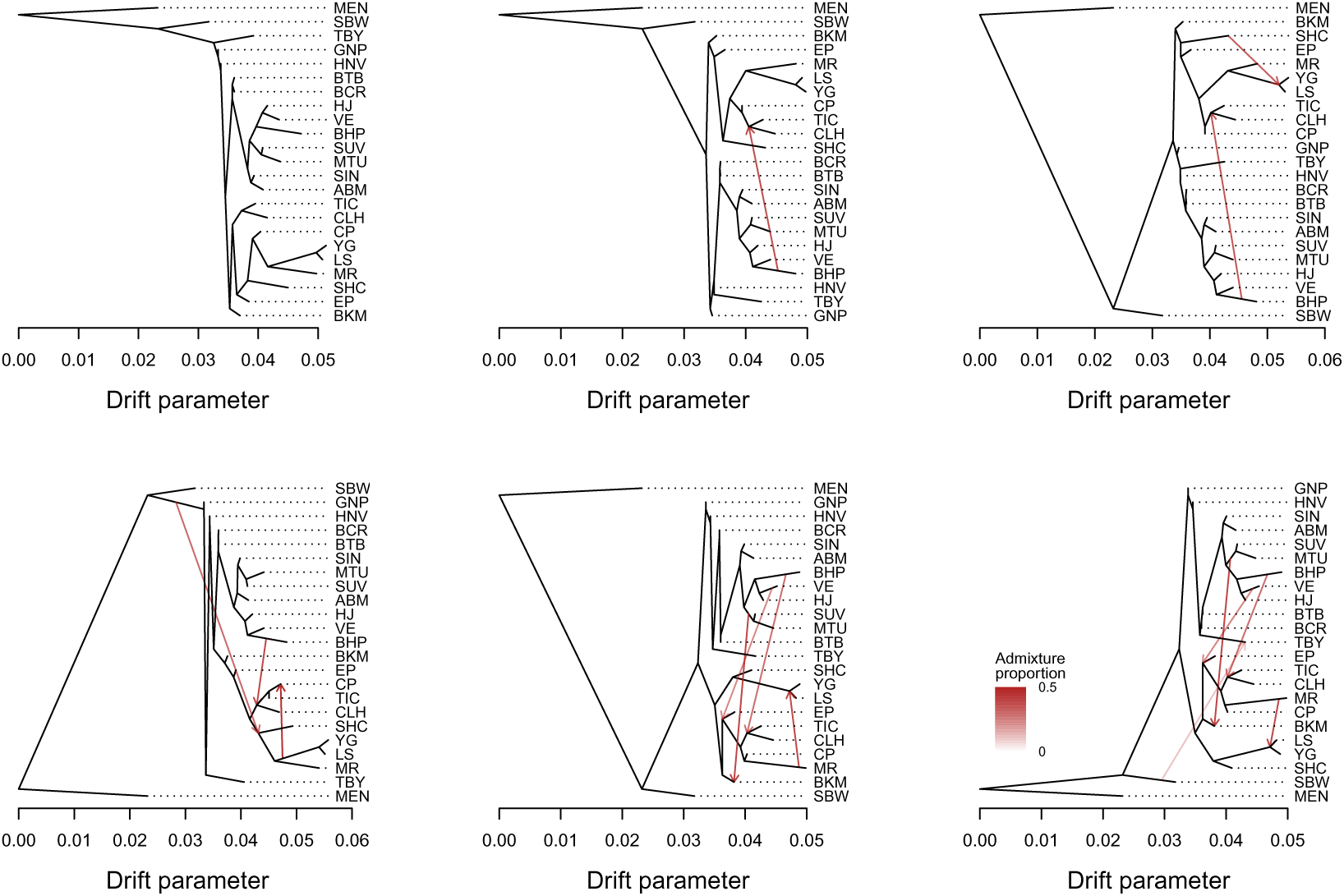
Population graphs for chromosome 11, with units in terms of a drift parameter proportional to evolutionary change and with the *L. argyrognomon* population (MEN) included as an outgroup. Graphs are shown for m = 0 (bifurcating tree) to five migration edges (red arrows). The admixture proportions associated with each are indicated by the intensity of each red arrow.

**Figure S19:**
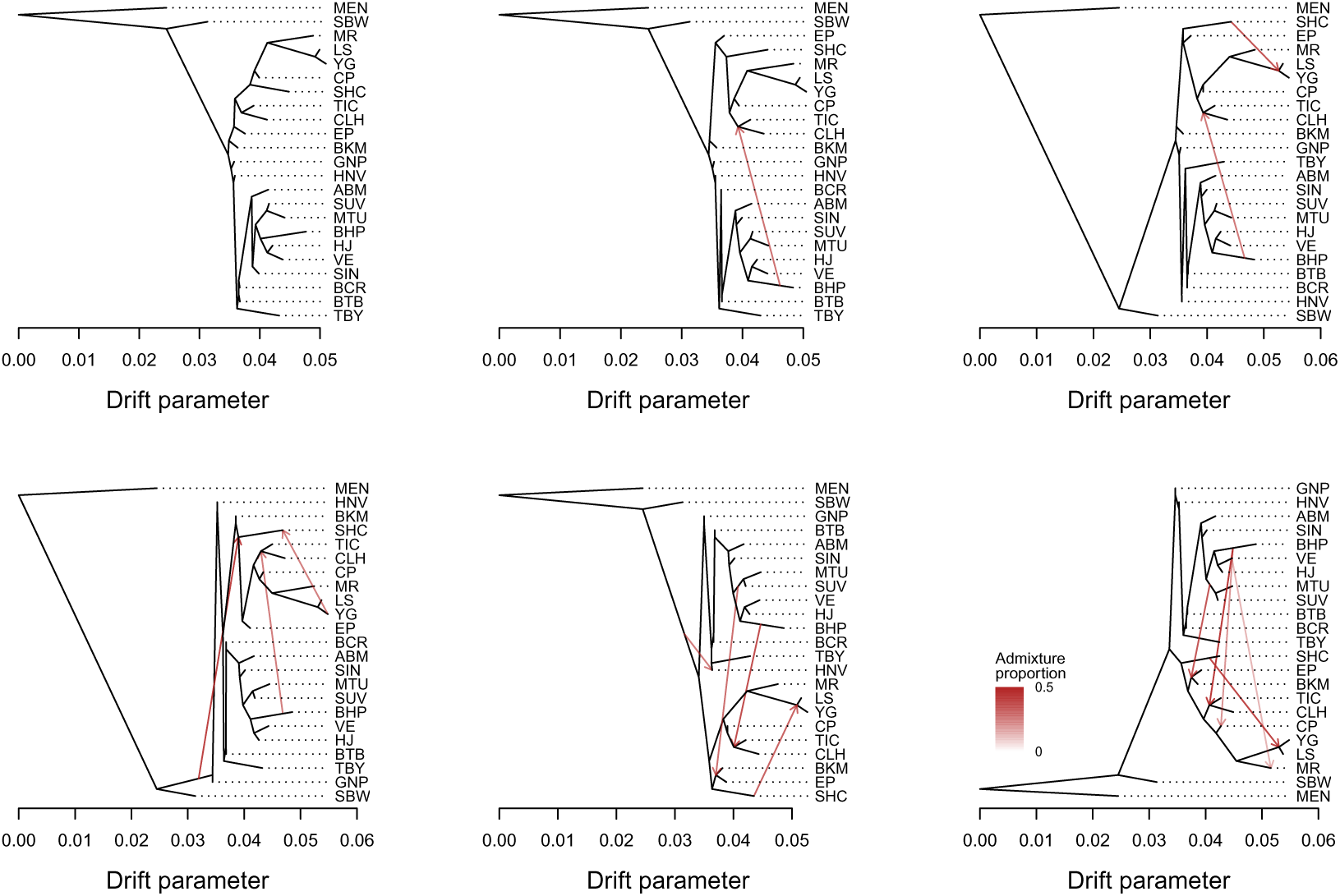
Population graphs for chromosome 12, with units in terms of a drift parameter proportional to evolutionary change and with the *L. argyrognomon* population (MEN) included as an outgroup. Graphs are shown for m = 0 (bifurcating tree) to five migration edges (red arrows). The admixture proportions associated with each are indicated by the intensity of each red arrow.

**Figure S20:**
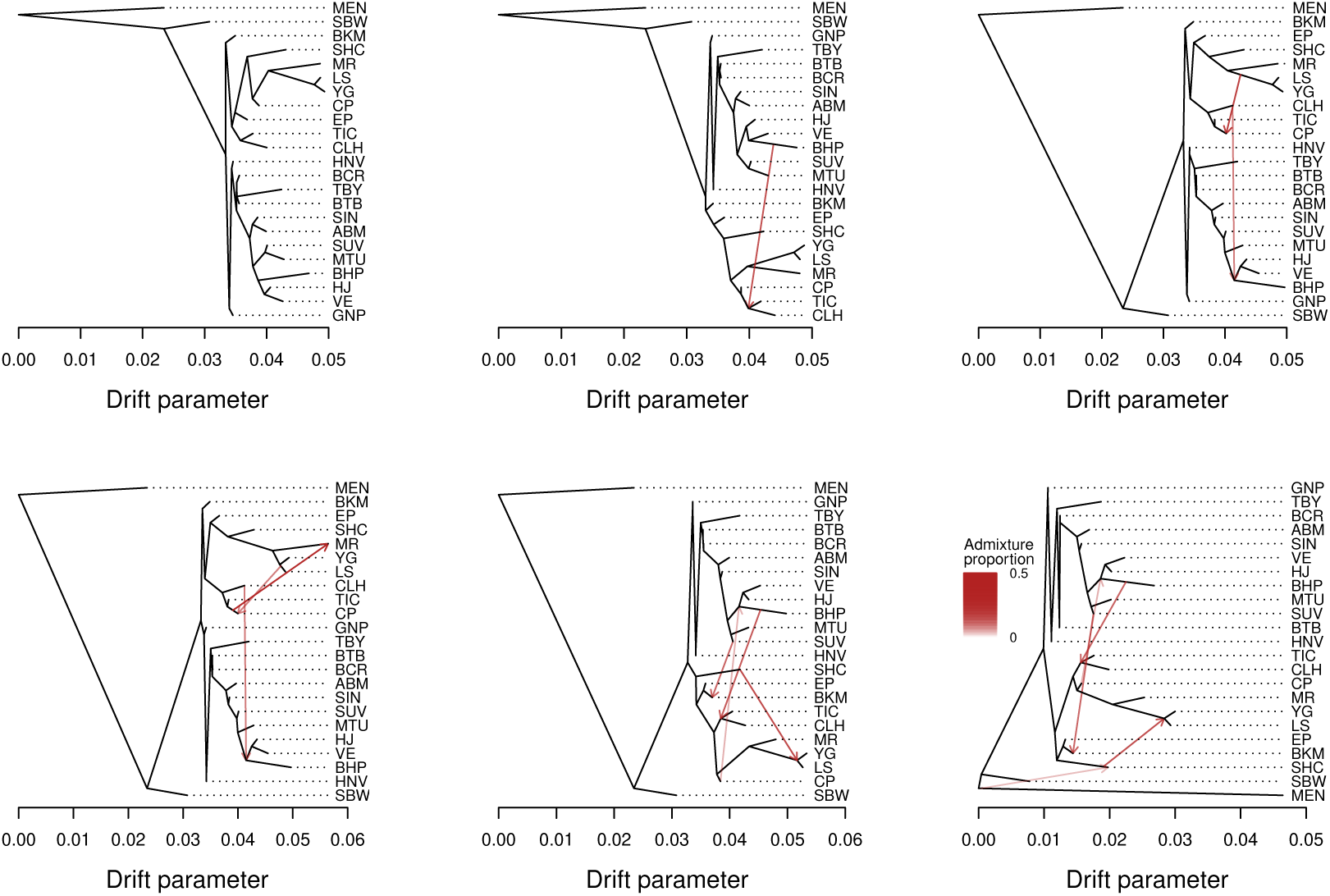
Population graphs for chromosome 13, with units in terms of a drift parameter proportional to evolutionary change and with the *L. argyrognomon* population (MEN) included as an outgroup. Graphs are shown for m = 0 (bifurcating tree) to five migration edges (red arrows). The admixture proportions associated with each are indicated by the intensity of each red arrow.

**Figure S21:**
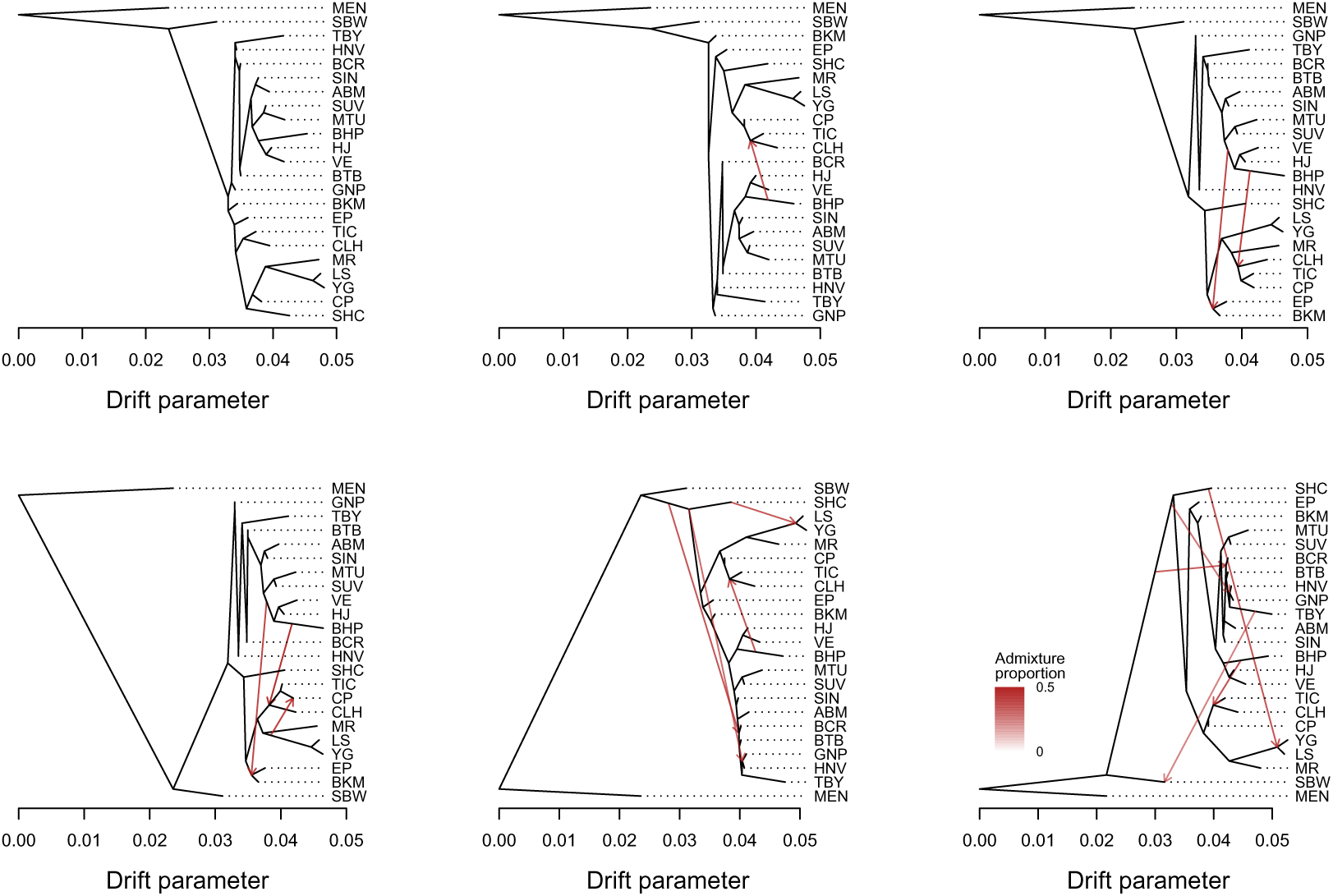
Population graphs for chromosome 14, with units in terms of a drift parameter proportional to evolutionary change and with the *L. argyrognomon* population (MEN) included as an outgroup. Graphs are shown for m = 0 (bifurcating tree) to five migration edges (red arrows). The admixture proportions associated with each are indicated by the intensity of each red arrow.

**Figure S22:**
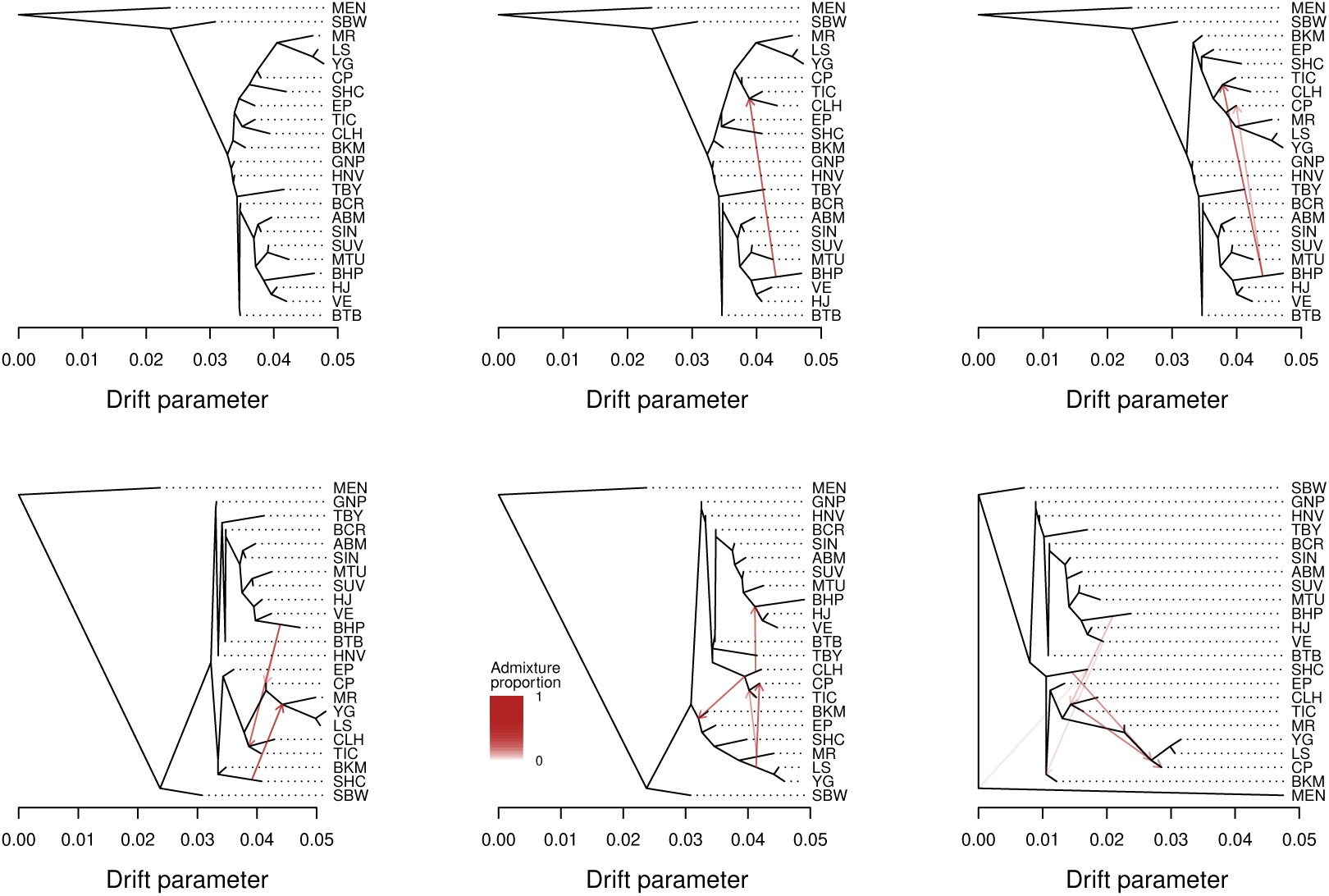
Population graphs for chromosome 15, with units in terms of a drift parameter proportional to evolutionary change and with the *L. argyrognomon* population (MEN) included as an outgroup. Graphs are shown for m = 0 (bifurcating tree) to five migration edges (red arrows). The admixture proportions associated with each are indicated by the intensity of each red arrow.

**Figure S23:**
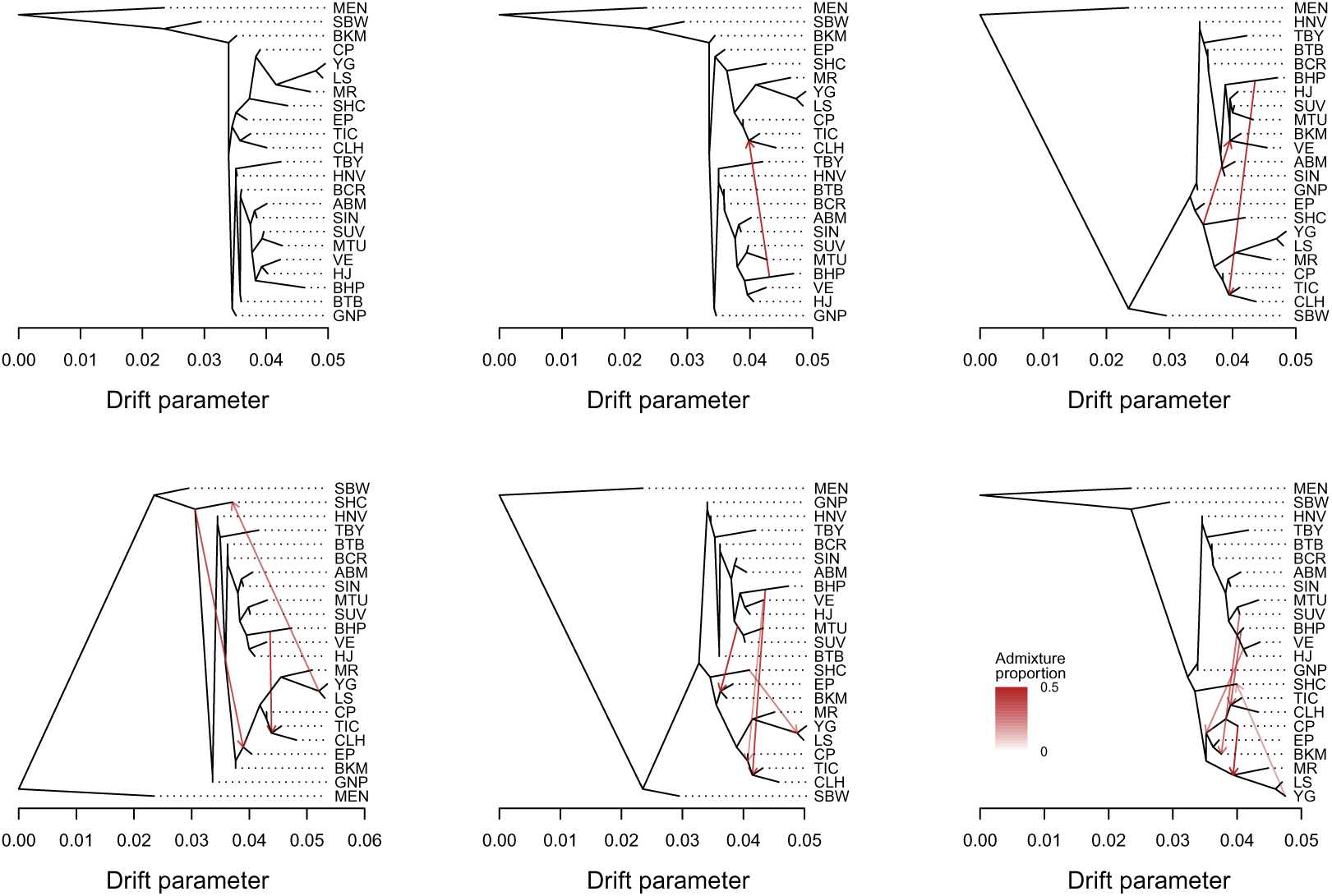
Population graphs for chromosome 16, with units in terms of a drift parameter proportional to evolutionary change and with the *L. argyrognomon* population (MEN) included as an outgroup. Graphs are shown for m = 0 (bifurcating tree) to five migration edges (red arrows). The admixture proportions associated with each are indicated by the intensity of each red arrow.

**Figure S24:**
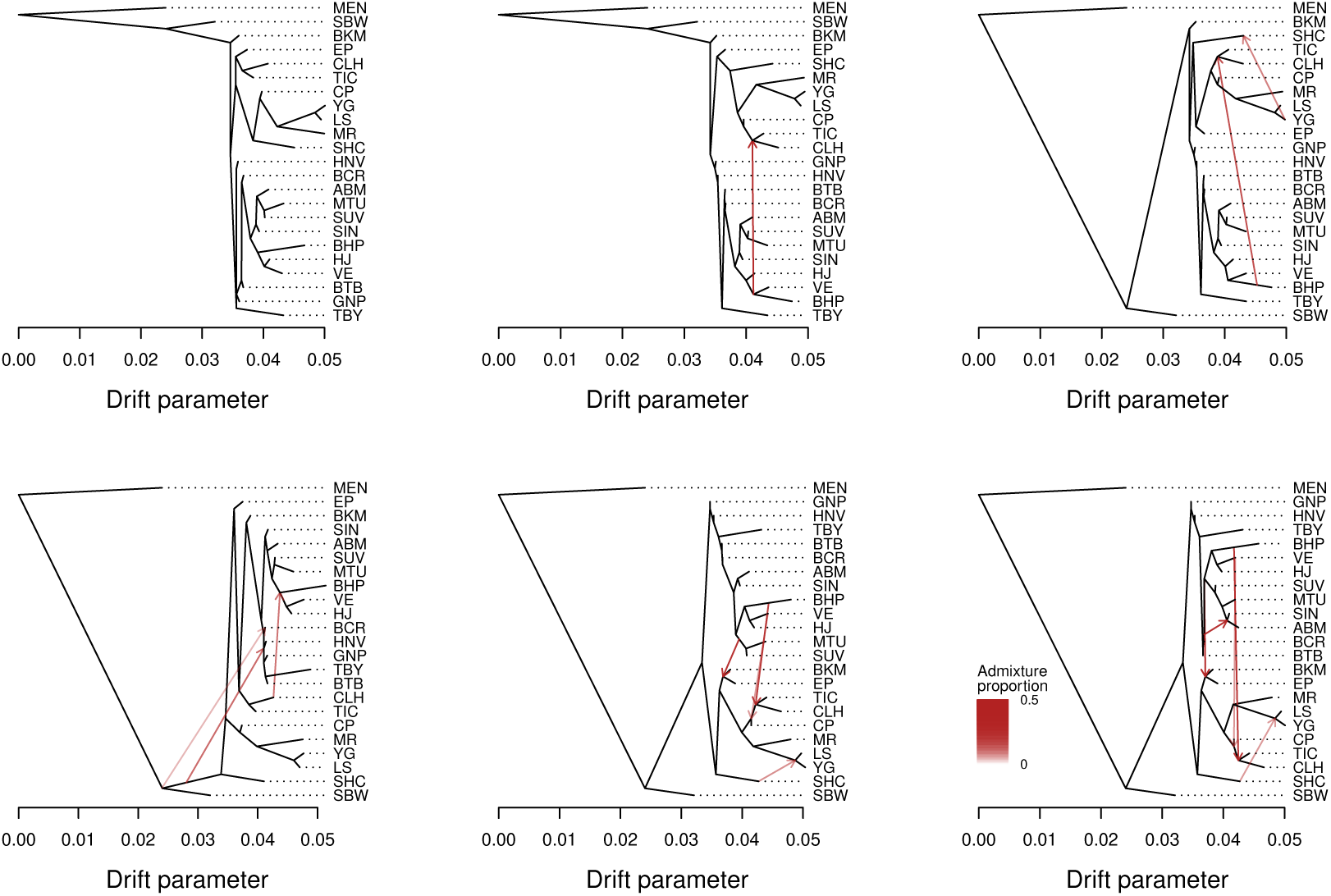
Population graphs for chromosome 17, with units in terms of a drift parameter proportional to evolutionary change and with the *L. argyrognomon* population (MEN) included as an outgroup. Graphs are shown for m = 0 (bifurcating tree) to five migration edges (red arrows). The admixture proportions associated with each are indicated by the intensity of each red arrow.

**Figure S25:**
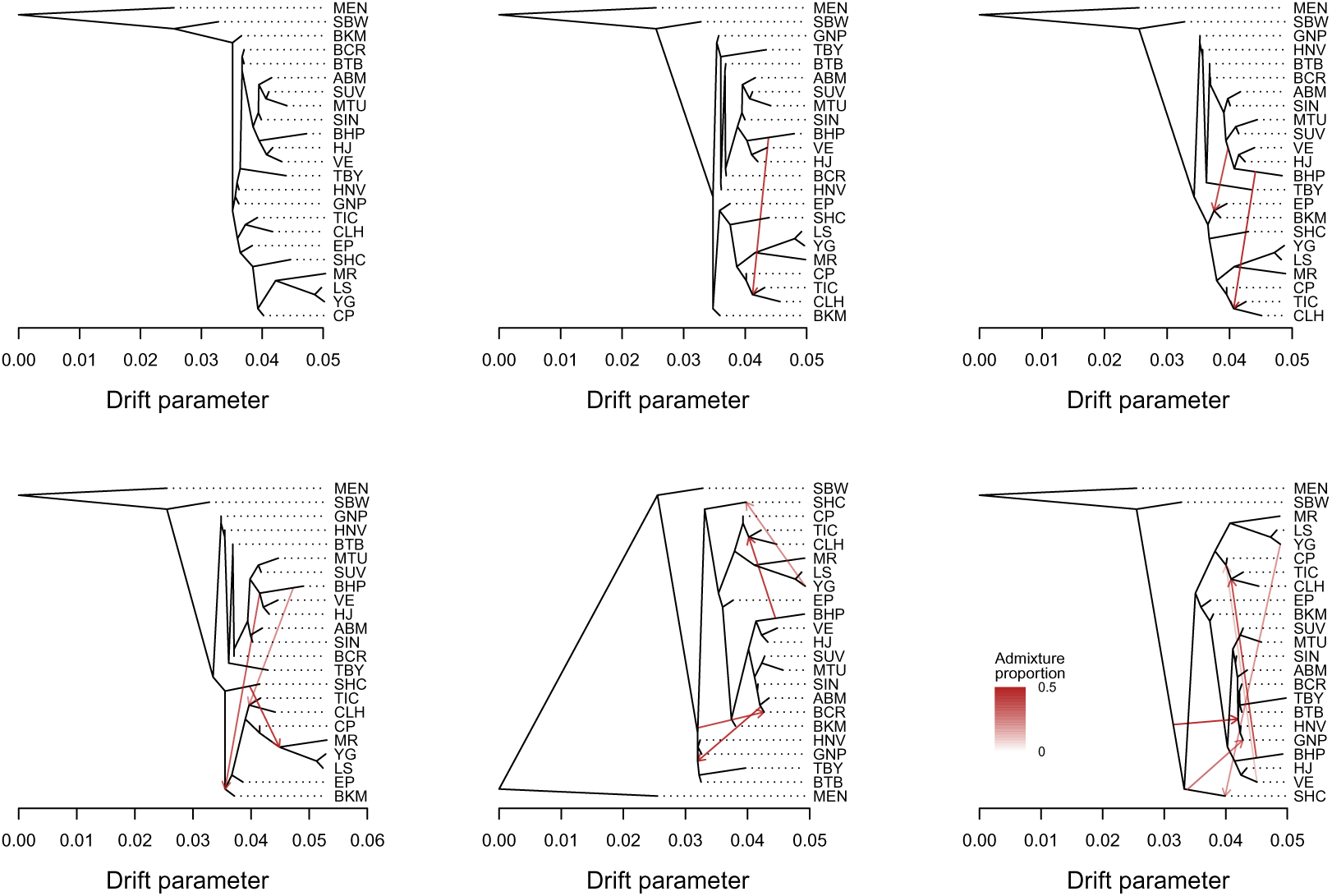
Population graphs for chromosome 18, with units in terms of a drift parameter proportional to evolutionary change and with the *L. argyrognomon* population (MEN) included as an outgroup. Graphs are shown for m = 0 (bifurcating tree) to five migration edges (red arrows). The admixture proportions associated with each are indicated by the intensity of each red arrow.

**Figure S26:**
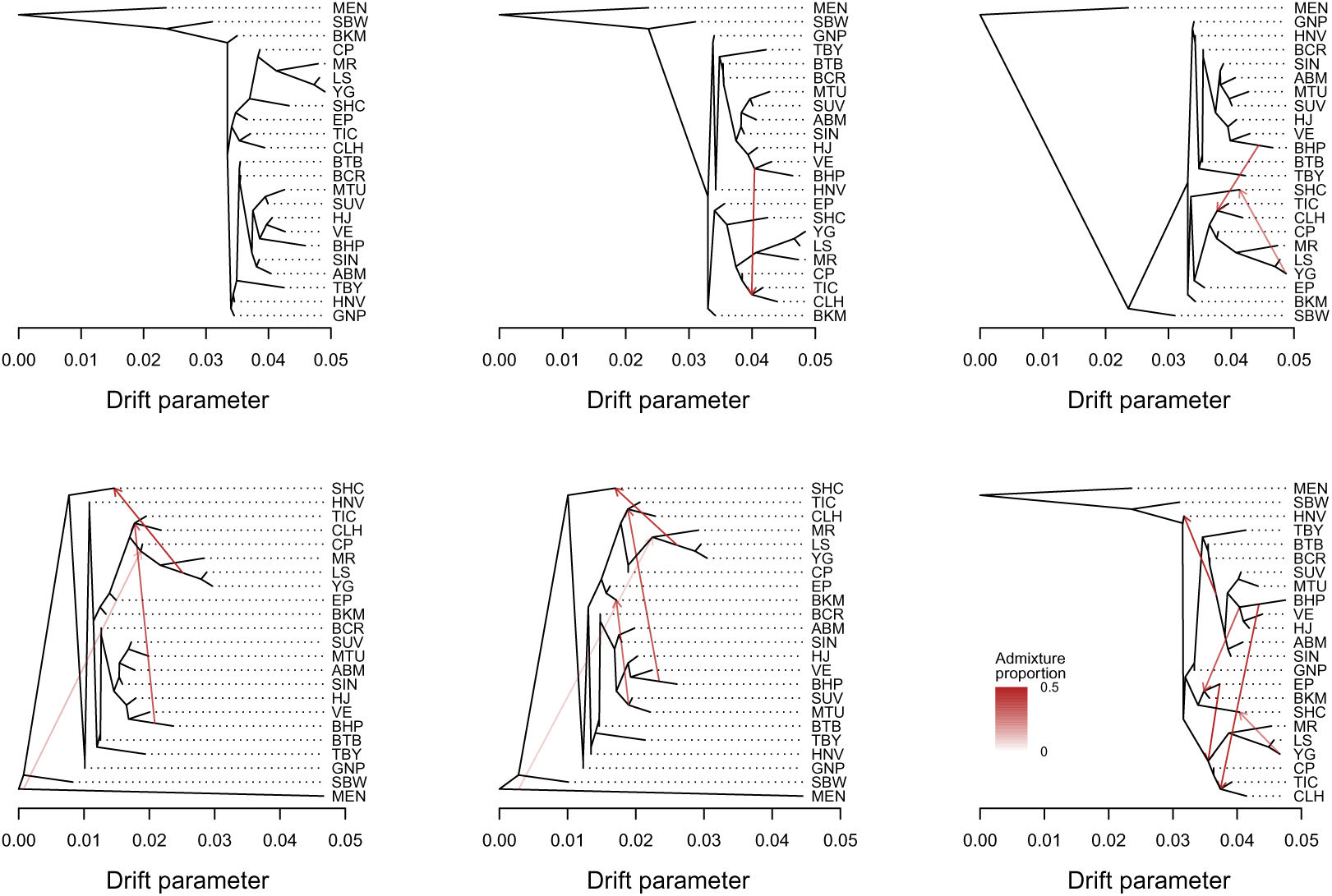
Population graphs for chromosome 19, with units in terms of a drift parameter proportional to evolutionary change and with the *L. argyrognomon* population (MEN) included as an outgroup. Graphs are shown for m = 0 (bifurcating tree) to five migration edges (red arrows). The admixture proportions associated with each are indicated by the intensity of each red arrow.

**Figure S27:**
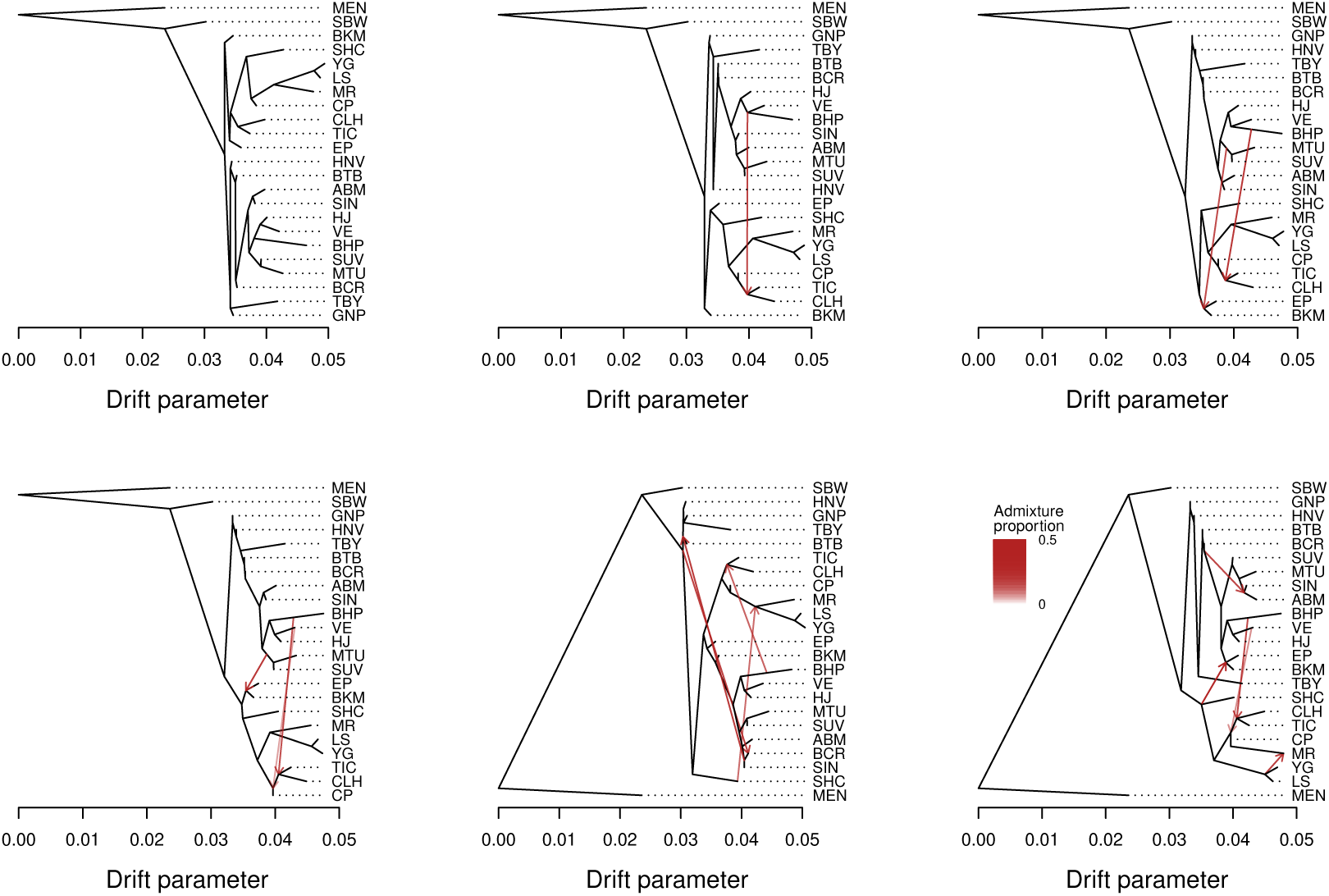
Population graphs for chromosome 20, with units in terms of a drift parameter proportional to evolutionary change and with the *L. argyrognomon* population (MEN) included as an outgroup. Graphs are shown for m = 0 (bifurcating tree) to five migration edges (red arrows). The admixture proportions associated with each are indicated by the intensity of each red arrow.

**Figure S28:**
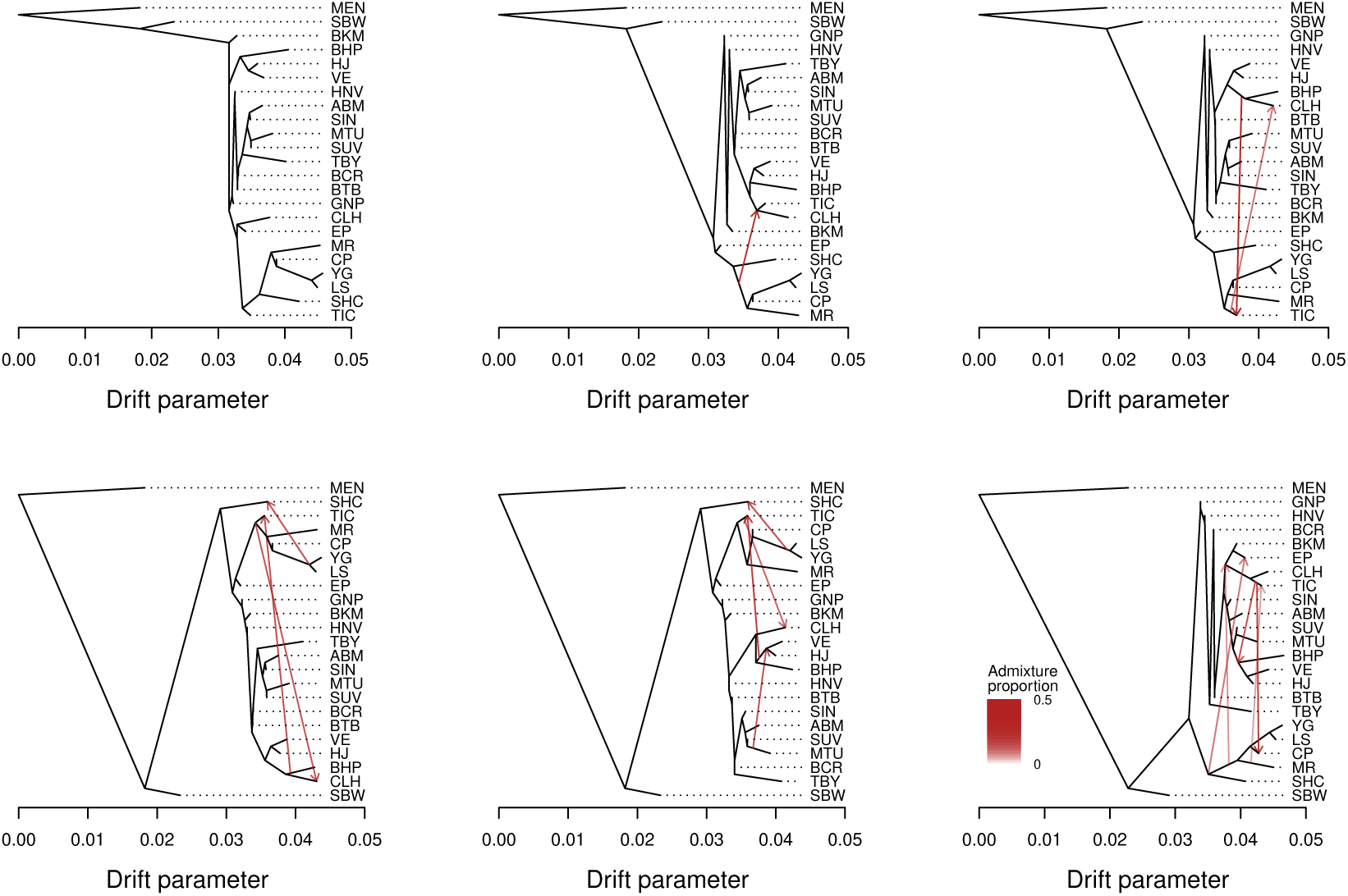
Population graphs for chromosome 21, with units in terms of a drift parameter proportional to evolutionary change and with the *L. argyrognomon* population (MEN) included as an outgroup. Graphs are shown for m = 0 (bifurcating tree) to five migration edges (red arrows). The admixture proportions associated with each are indicated by the intensity of each red arrow.

**Figure S29:**
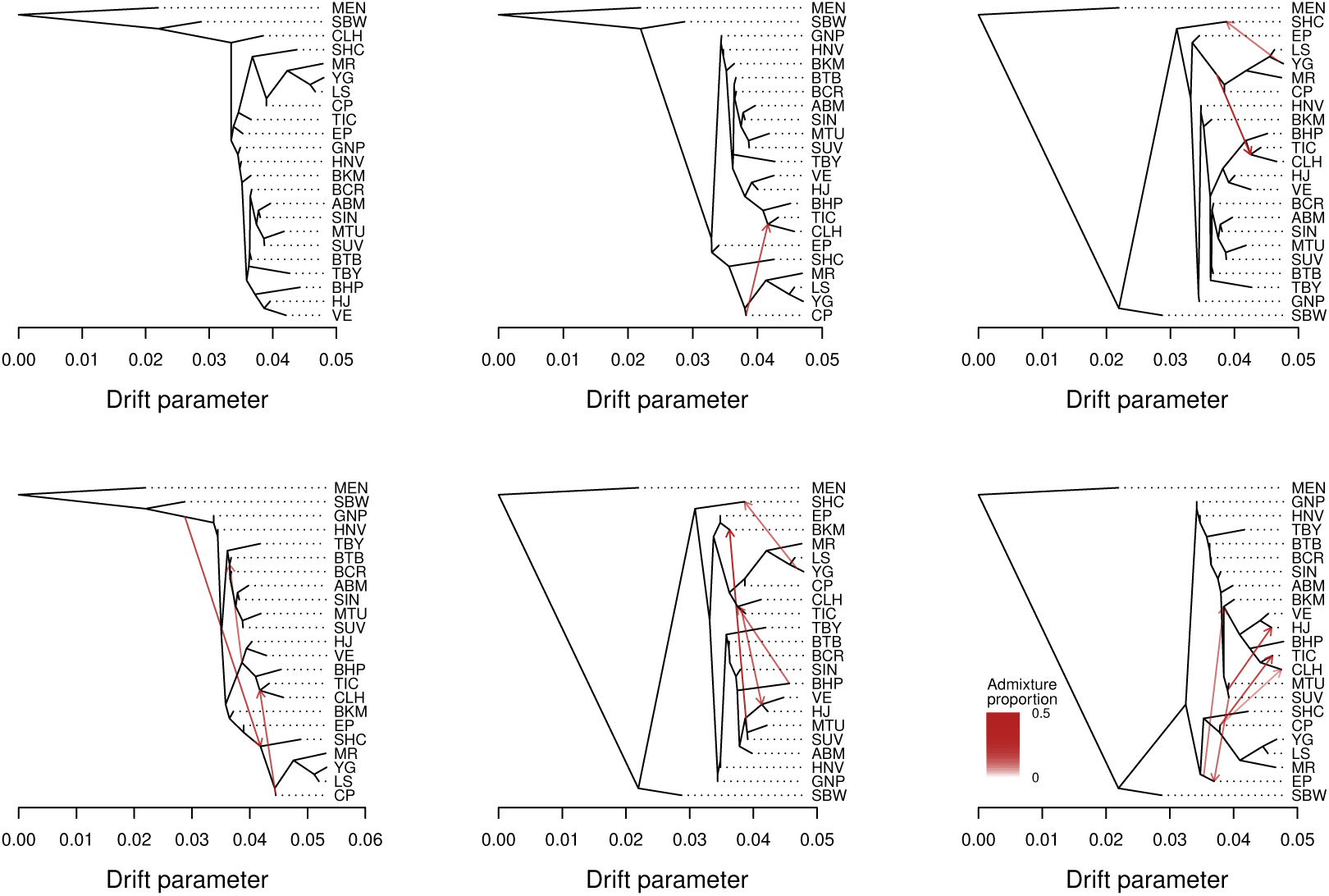
Population graphs for chromosome 22, with units in terms of a drift parameter proportional to evolutionary change and with the *L. argyrognomon* population (MEN) included as an outgroup. Graphs are shown for m = 0 (bifurcating tree) to five migration edges (red arrows). The admixture proportions associated with each are indicated by the intensity of each red arrow.

**Figure S30:**
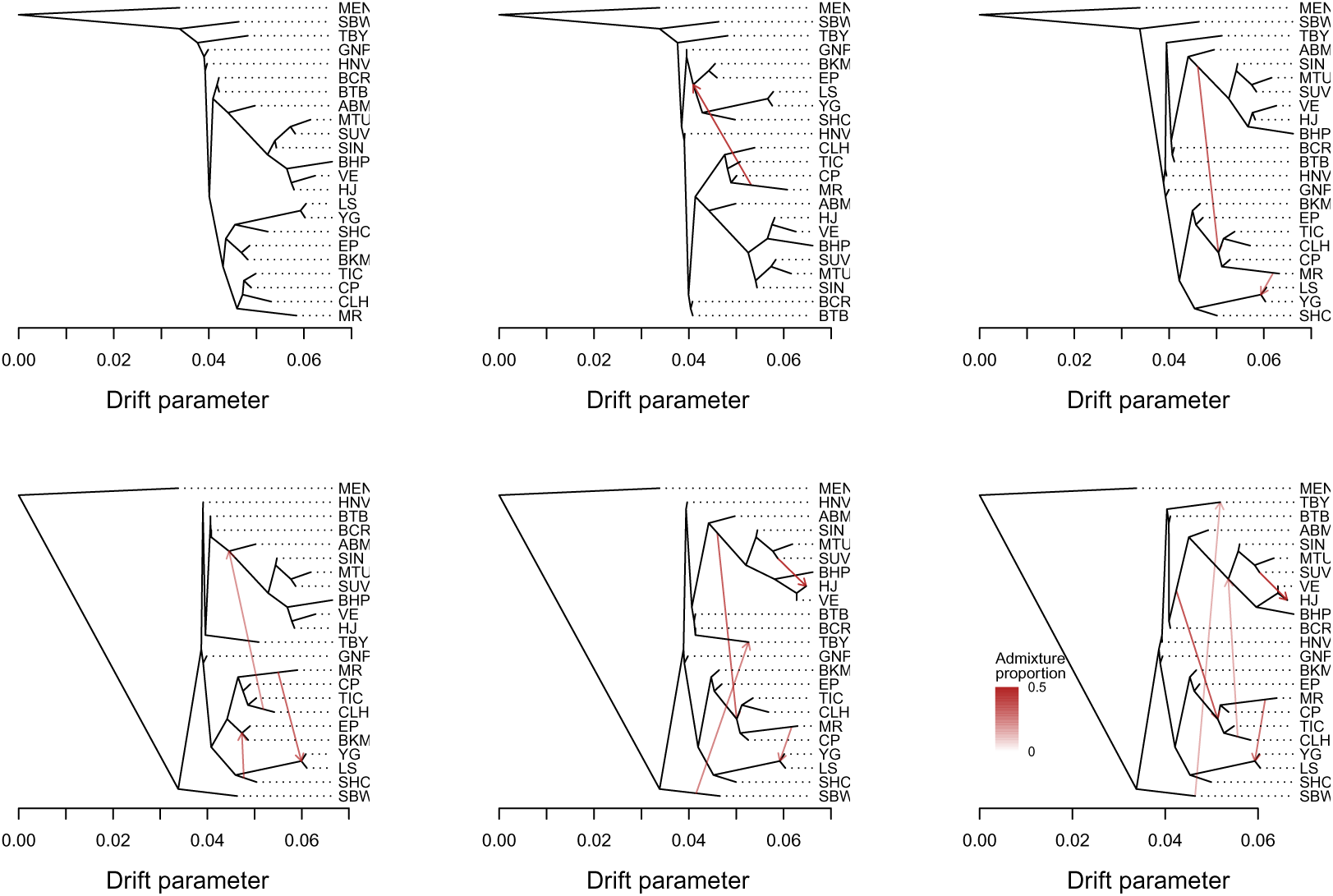
Population graphs for the Z chromosome, with units in terms of a drift parameter proportional to evolutionary change and with the *L. argyrognomon* population (MEN) included as an outgroup. Graphs are shown for m = 0 (bifurcating tree) to five migration edges (red arrows). The admixture proportions associated with each are indicated by the intensity of each red arrow.

**Figure S31:**
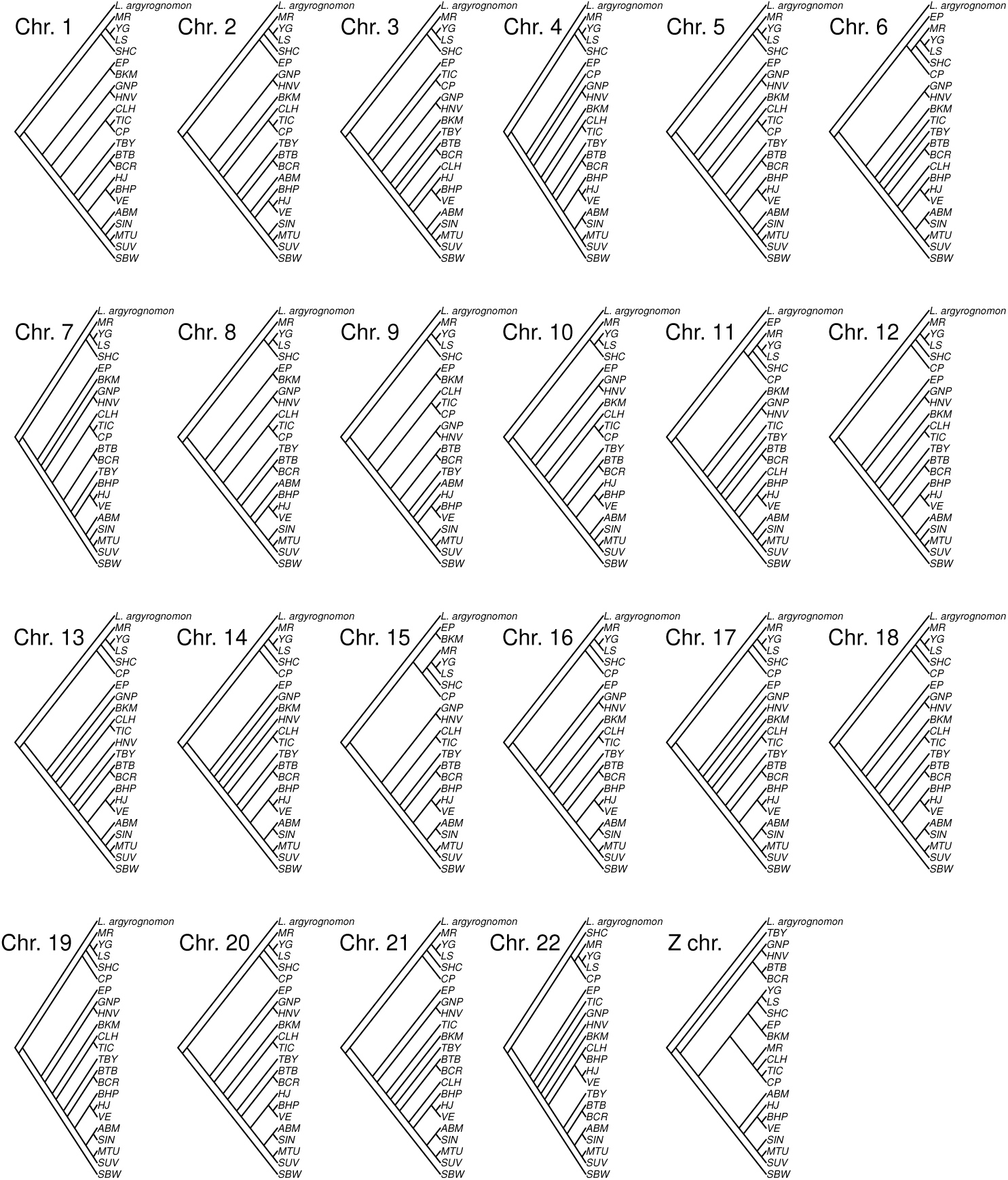
Tree topologies (as a cladogram) for each chromosome based on the CASTER-pair model. *Lycaeides argyrognomon* (population = MEN) was used as the outgroup.

**Figure S32:**
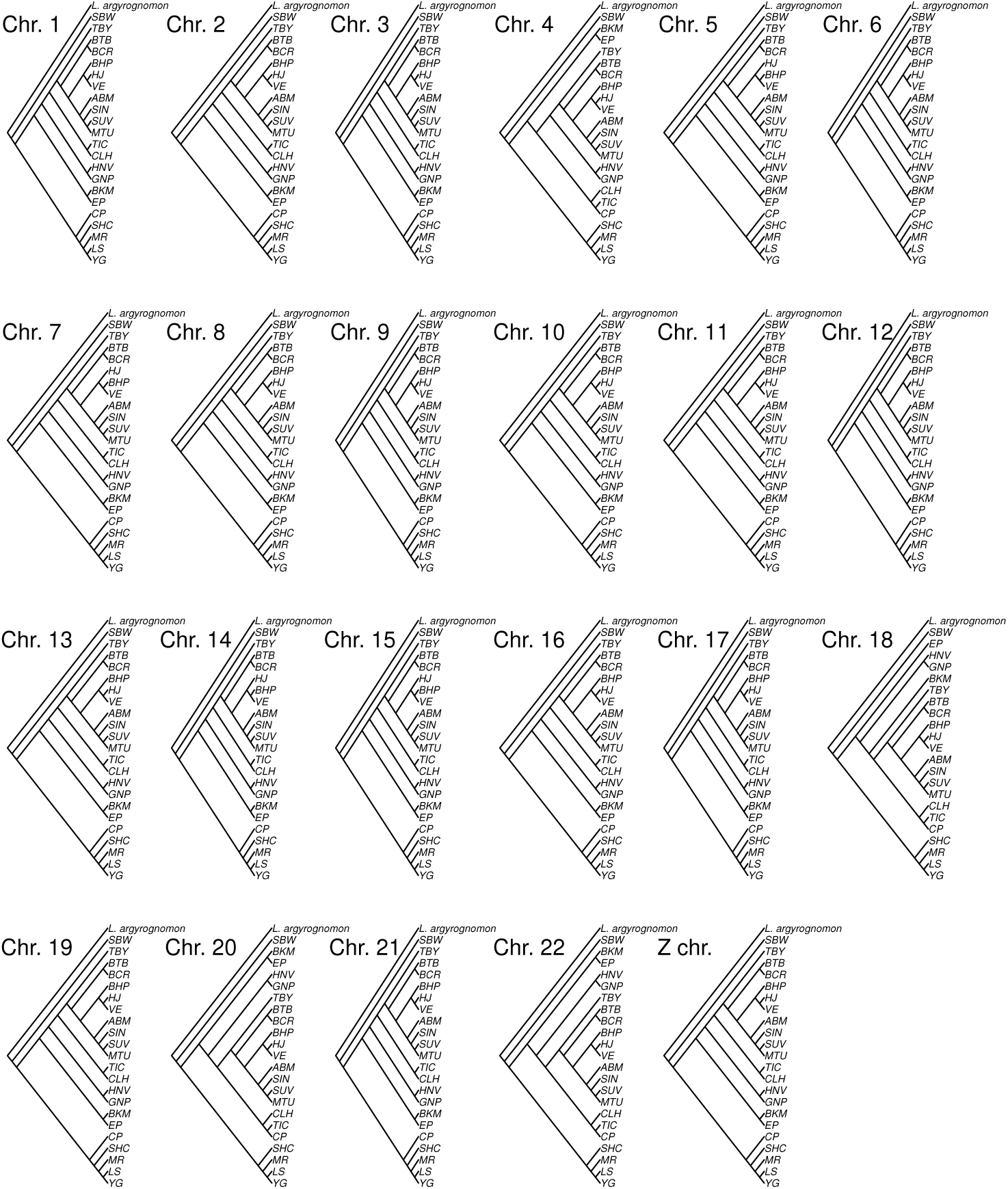
Example tree topologies (as a cladogram) based on the CASTER-site model with SNPs permuted among chromosomes. *Lycaeides argyrognomon* (population = MEN) was used as the outgroup. Results are shown for one permutation/randomization.

**Figure S33:**
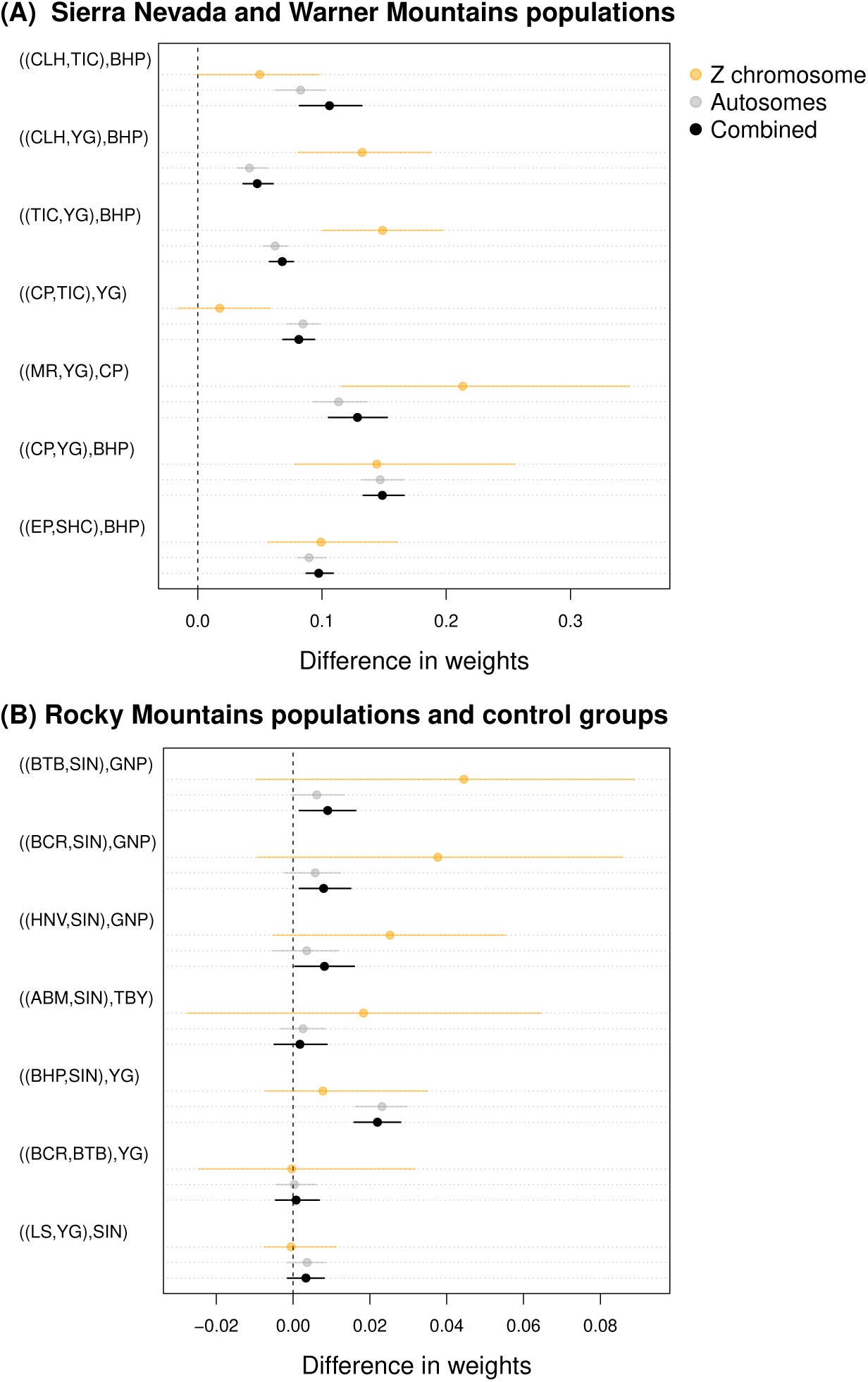
Dot plots summarizing the symmetry of unnormalized weights for the two less common topologies from each four-taxon, windowed CASTER analysis. Results are shown for the Z chromosome only, the autosomes only, and the Z chromosome and autosomes combined. In each case, the dominant topology was defined as the topology with the highest mean weight across the relevant set of windows (Z only, autosomes only, or all windows). The mean difference was then computed between support for the two alternative topologies; a mean difference of 0 is expected under the multispecies coalescent with only incomplete lineage sorting (ILS) (denoted by a horizontal line). Point estimates (dots) and 95% confidence intervals (horizontal lines) for these differences were obtained by bootstrap resampling of datasets thinned to minimize the effects of non-independence between adjacent windows. Panels (A) and (B) show the four-taxon sets from Figures 4 and 5, respectively.

**Figure S34:**
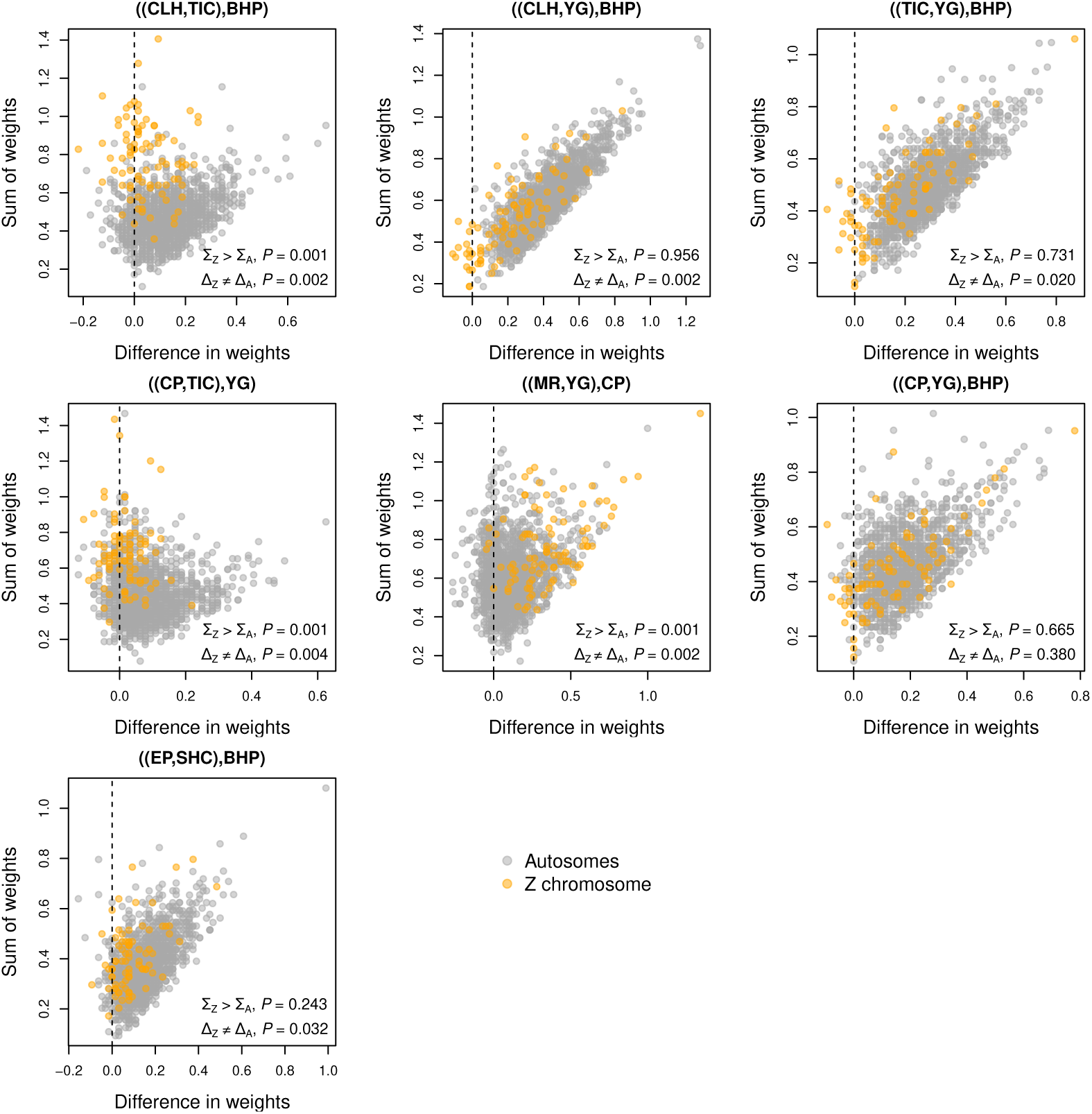
Scatterplots summarizing the symmetry of unnormalized weights for the two less common topologies (difference in weights) versus phylogenetic signal (sum of weights) for each window from each four-taxon, windowed CASTER analysis. Results are shown for the four-taxon sets from Figure 4. For this analysis, we defined the primary (species-tree) topology as the topology recovered by BEAST and the less common topologies as the two alternatives (compare this to Figure S35). Points denote windows and are colored to indicate autosomal (gray) versus Z chromosome (orange) windows. *P*-values are reported for tests of the statistical hypotheses that (i) the mean sum of weights for Z windows (Σ*_Z_*) exceeds the mean sum of weights for autosomal windows (Σ*_A_*), and (ii) the mean difference in weights between the two less common topologies for Z windows (Δ*_Z_*) and autosomal windows (Δ*_A_*) is not zero. Permutation tests (1000 permutations) of datasets thinned to minimize the effects of non-independence between adjacent windows were used to evaluate these hypotheses.

**Figure S35:**
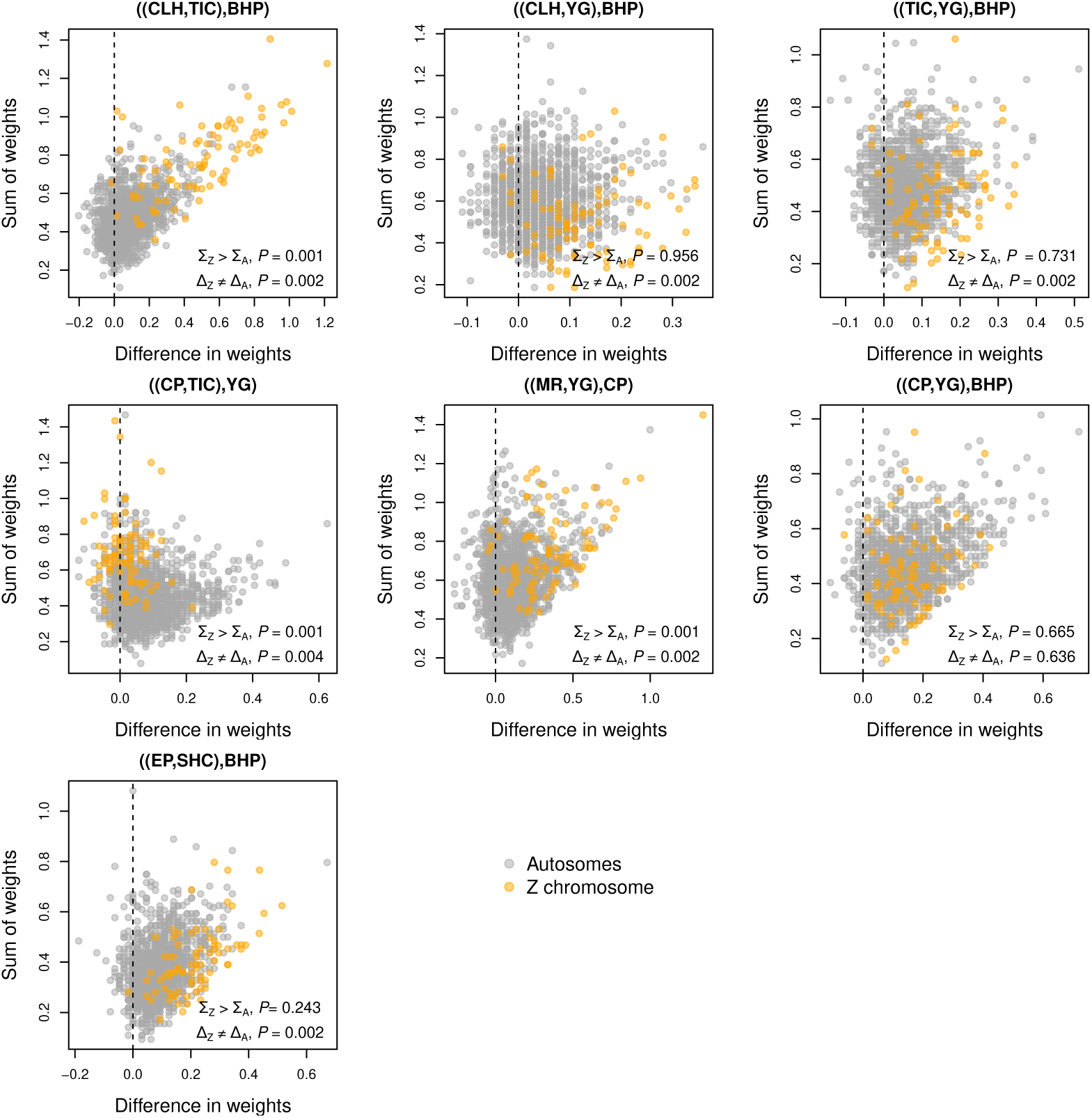
Scatterplots summarizing the symmetry of unnormalized weights for the two less common topologies (difference in weights) versus phylogenetic signal (sum of weights) for each window from each four-taxon, windowed CASTER analysis. Results are shown for the four-taxon sets from Figure 4. For this analysis, we defined the primary (species-tree) topology as the topology with the highest mean score across all windows and the less common topologies as the two alternatives (compare this to Figure S34). Points denote windows and are colored to indicate autosomal (gray) versus Z chromosome (orange) windows. *P*-values are reported for tests of the statistical hypotheses that (i) the mean sum of weights for Z windows (Σ*_Z_*) exceeds the mean sum of weights for autosomal windows (Σ*_A_*), and (ii) the mean difference in weights between the two less common topologies for Z windows (Δ*_Z_*) and autosomal windows (Δ*_A_*) is not zero. Permutation tests (1000 permutations) of datasets thinned to minimize the effects of non-independence between adjacent windows were used to evaluate these hypotheses.

**Figure S36:**
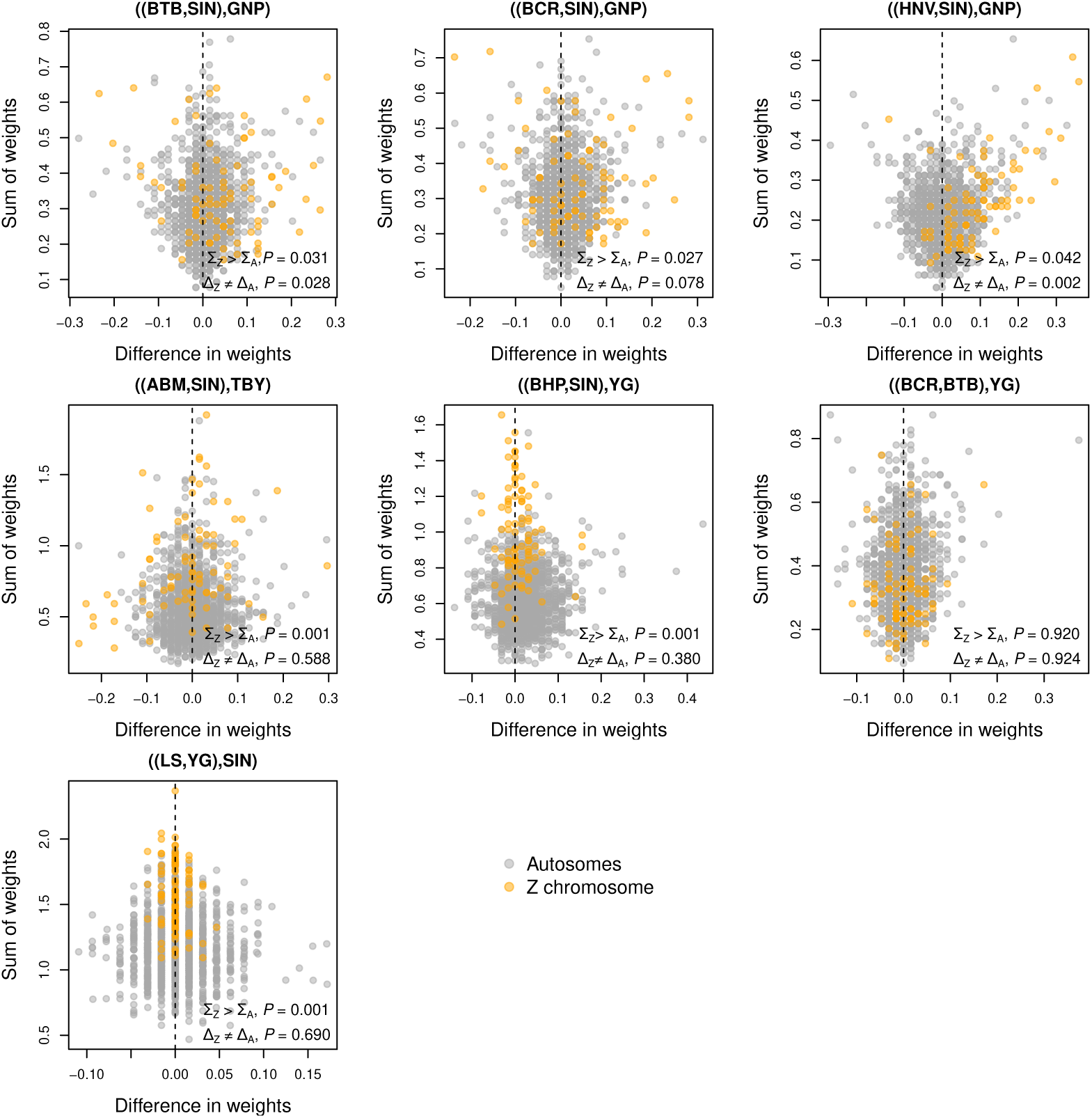
Scatterplots summarizing the symmetry of unnormalized weights for the two less common topologies (difference in weights) versus phylogenetic signal (sum of weights) for each window from each four-taxon, windowed CASTER analysis. Results are shown for the four-taxon sets from Figure 5. For this analysis, we defined the primary (species-tree) topology as the topology with the highest mean score across all windows, which is also the topology recovered by BEAST, and the less common topologies as the two alternatives. Points denote windows and are colored to indicate autosomal (gray) versus Z chromosome (orange) windows. *P*-values are reported for tests of the statistical hypotheses that (i) the mean sum of weights for Z windows (Σ*_Z_*) exceeds the mean sum of weights for autosomal windows (Σ*_A_*), and (ii) the mean difference in weights between the two less common topologies for Z windows (Δ*_Z_*) and autosomal windows (Δ*_A_*) is not zero. Permutation tests (1000 permutations) of datasets thinned to minimize the effects of non-independence between adjacent windows were used to evaluate these hypotheses.

**Figure S37:**
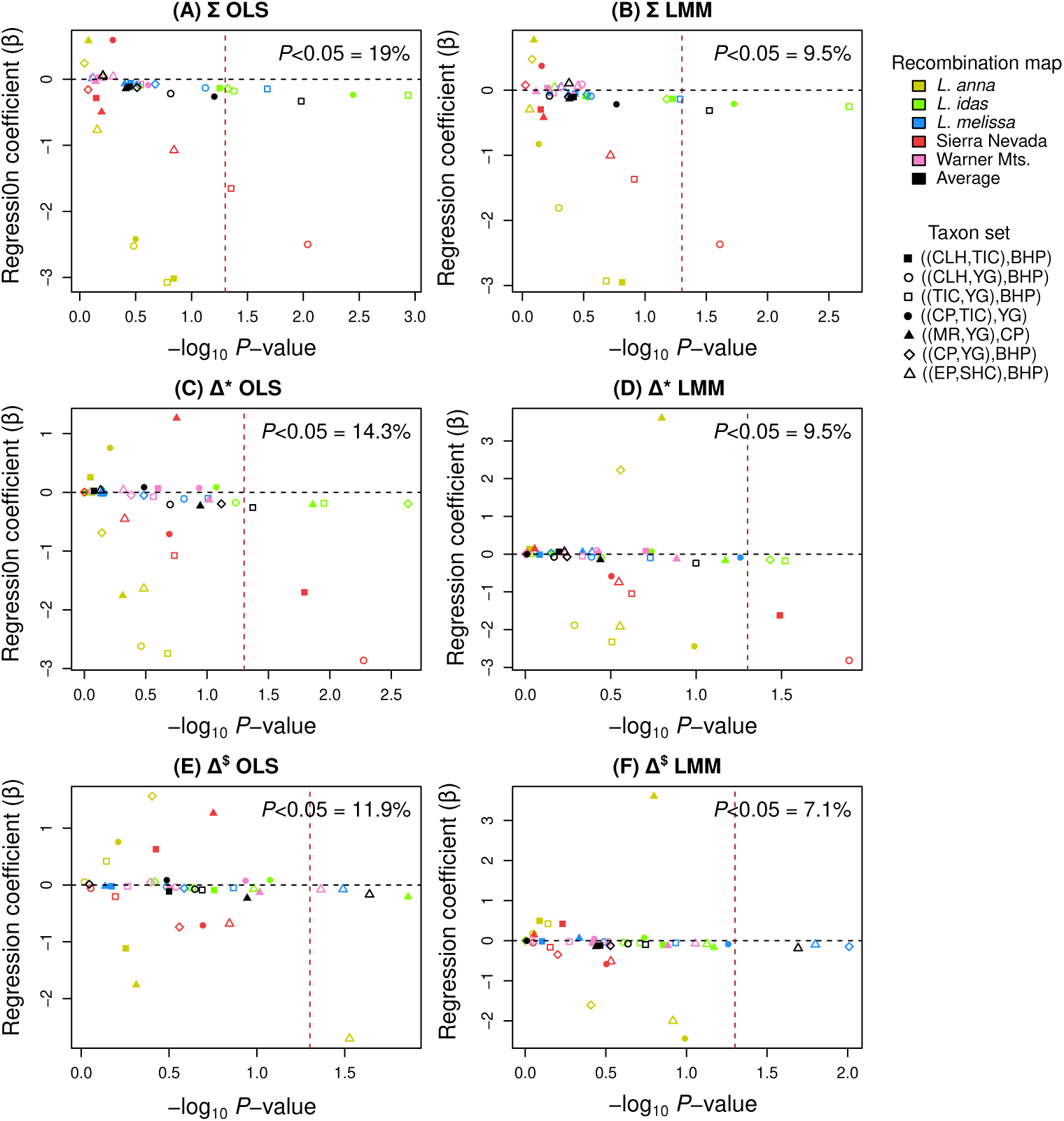
Scatterplots summarizing the relationship between recombination rate variation (*ρ* = 4*N_e_r*) and summaries of topology weights from CASTER for different combinations of four-taxon sets and recombination maps. Results are shown for the four-taxon sets in Figure 4. Each panel shows *P*-values and corresponding regression coefficients (*β*s) from ordinary least squares (OLS) linear models (A, C, E) or linear mixed models (LMMs) (B, D, F) relating topology summaries (dependent variables) to recombination rates (independent variables). In the latter, a random-effects term accounts for differences among chromosomes. Results are shown for the sum of topology weights (Σ; A, B) and the difference in weights for the alternative topologies (Δ), where alternative topologies are defined either as those not recovered by BEAST (Δ*; C, D) or as the two less common topologies (Δ^$^; E, F), which are sometimes, but not always, the same. Horizontal dashed lines denote *β* = 0, and vertical dashed lines denote −log_10_(*P* = 0.05). The percentage of regression models (four-taxon set × recombination map combinations) with *P <* 0.05 is reported in each panel.

**Figure S38:**
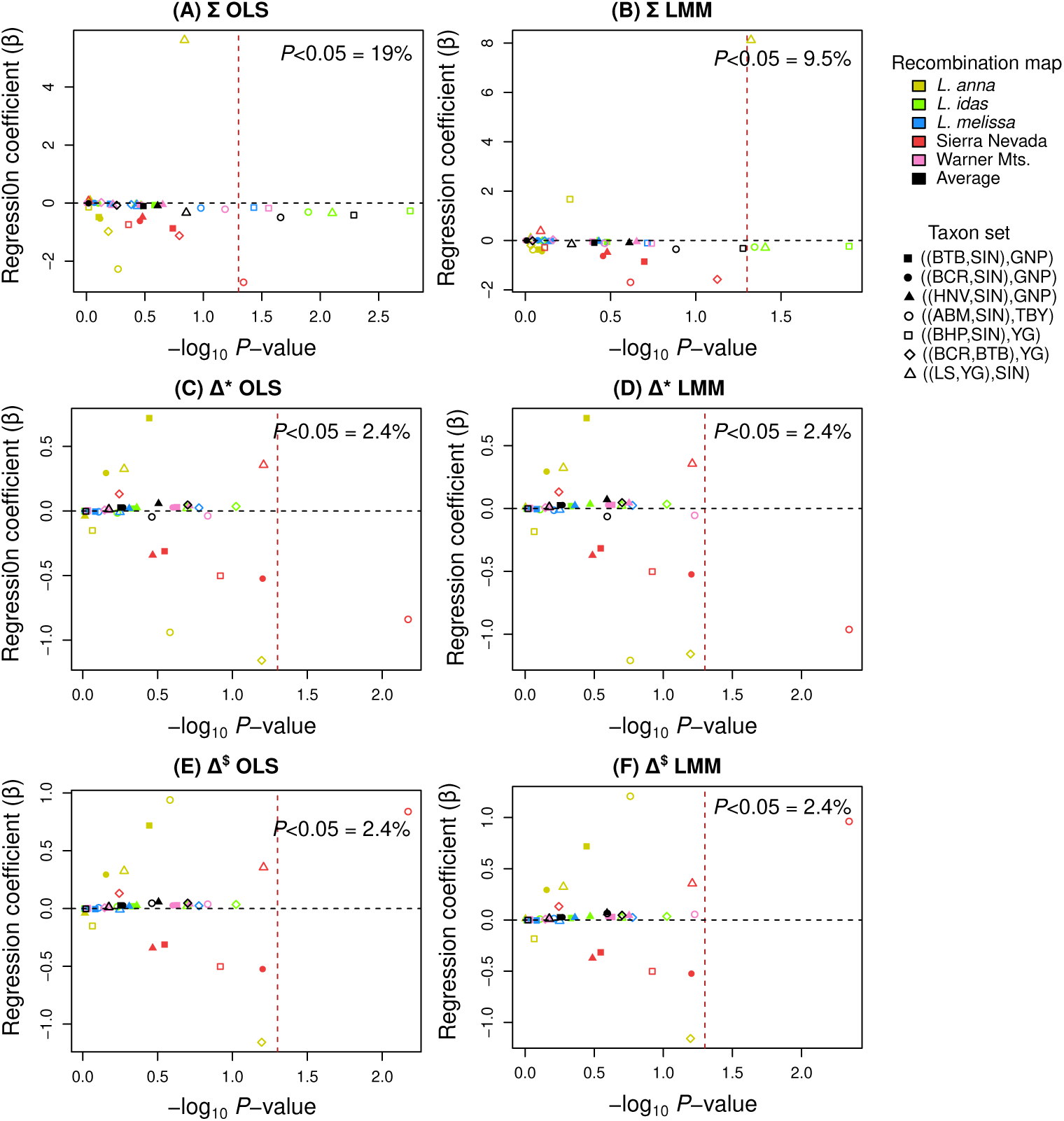
Scatterplots summarizing the relationship between recombination rate variation (*ρ* = 4*N_e_r*) and summaries of topology weights from CASTER for different combinations of four-taxon sets and recombination maps. Results are shown for the four-taxon sets in Figure 5. Each panel shows *P*-values and corresponding regression coefficients (*β*s) from ordinary least squares (OLS) linear models (A, C, E) or linear mixed models (LMMs) (B, D, F) relating topology summaries (dependent variables) to recombination rates (independent variables). In the latter, a random-effects term accounts for differences among chromosomes. Results are shown for the sum of topology weights (Σ; A, B) and the difference in weights for the alternative topologies (Δ), where alternative topologies are defined either as those not recovered by BEAST (Δ*; C, D) or as the two less common topologies (Δ^$^; E, F), which are sometimes, but not always, the same. Horizontal dashed lines denote *β* = 0, and vertical dashed lines denote −log_10_(*P* = 0.05). The percentage of regression models (four-taxon set × recombination map combinations) with *P <* 0.05 is reported in each panel.

**Figure S39:**
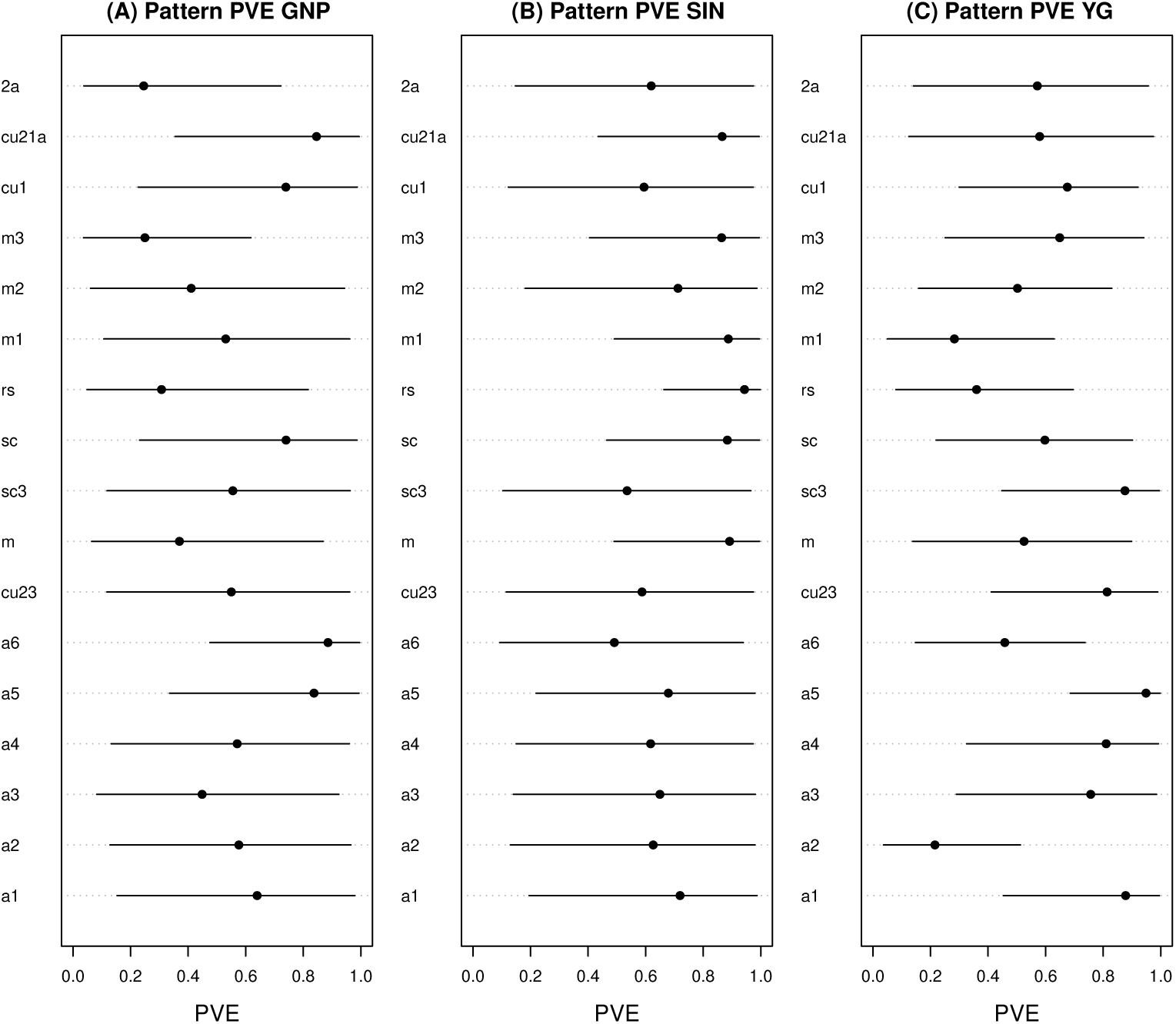
Dot plots depict Bayesian estimates of the proportion of wing pattern variation explained (PVE) by additive genetic effects for each of 17 pattern traits. Points and horizontal lines denote posterior medians and 80% equal-tail probability intervals, respectively. Results are shown for three populations: GNP (A), SIN (B) and YG (C). Pattern elements are defined in Figure 6.

**Figure S40:**
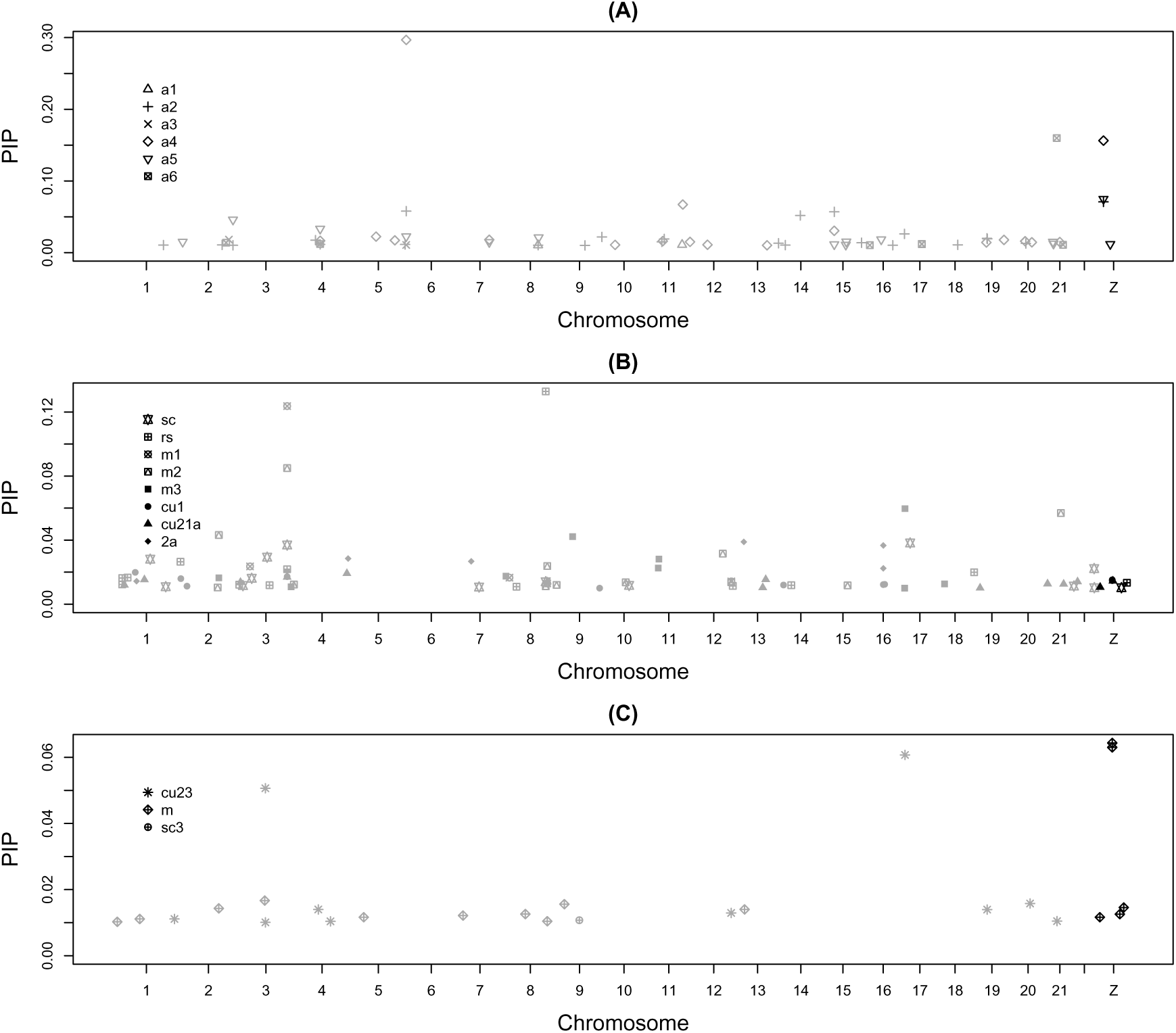
Manhattan plots of SNP posterior inclusion probabilities (PIPs) from polygenic association mapping of wing pattern traits in GNP. Only SNPs with PIPs ≥ 0.01 are shown. Panels correspond to different subsets of wing pattern traits: (A) border eyespots (aurorae), (B) distal-center black dots, and (C) proximal-center black dots. SNPs on autosomes are shown in gray, whereas SNPs on the Z chromosome are shown in black.

**Figure S41:**
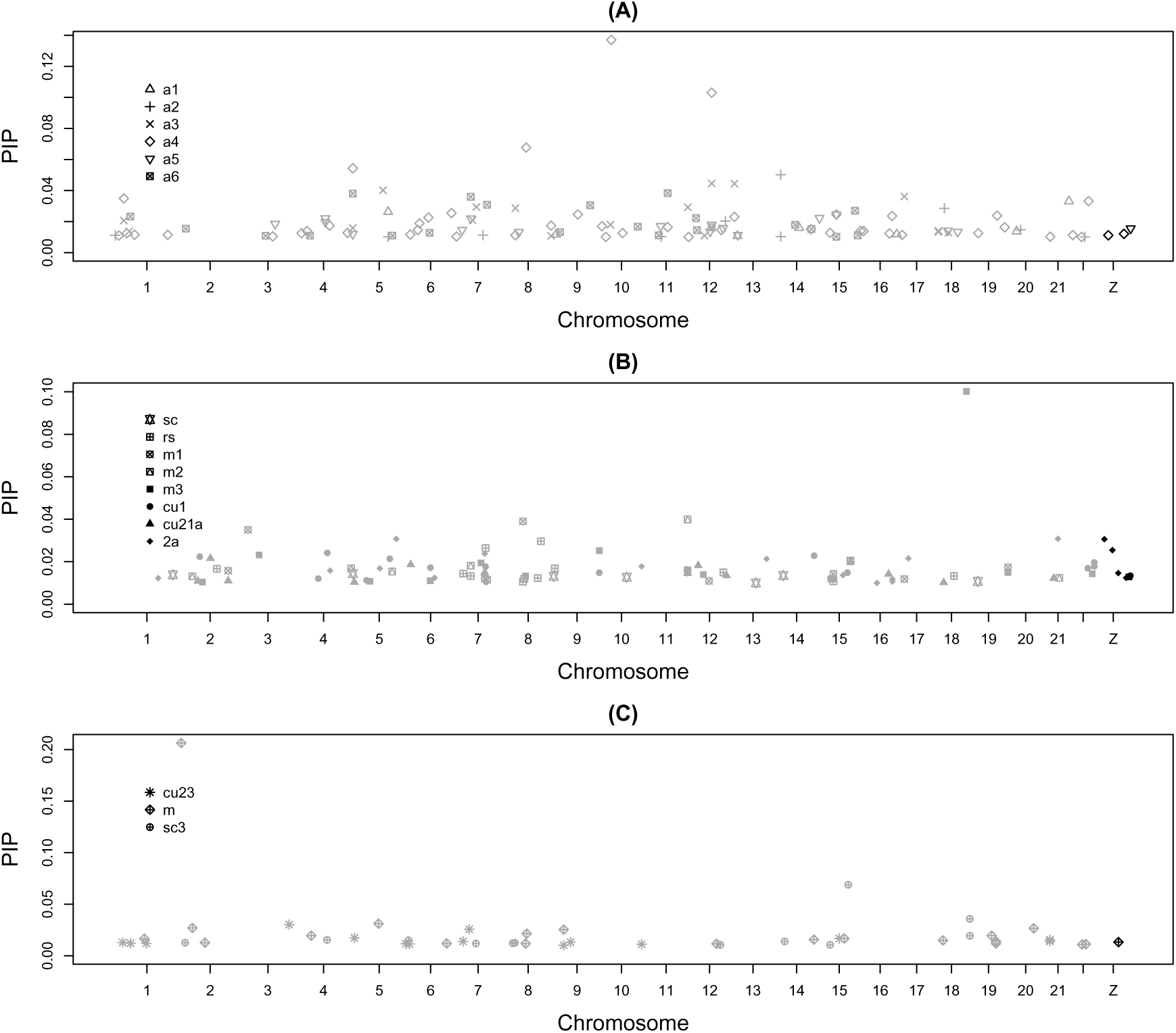
Manhattan plots of SNP posterior inclusion probabilities (PIPs) from polygenic association mapping of wing pattern traits in SIN. Only SNPs with PIPs ≥ 0.01 are shown. Panels correspond to different subsets of wing pattern traits: (A) border eyespots (aurorae), (B) distal-center black dots, and (C) proximal-center black dots. SNPs on autosomes are shown in gray, whereas SNPs on the Z chromosome are shown in black.

**Figure S42:**
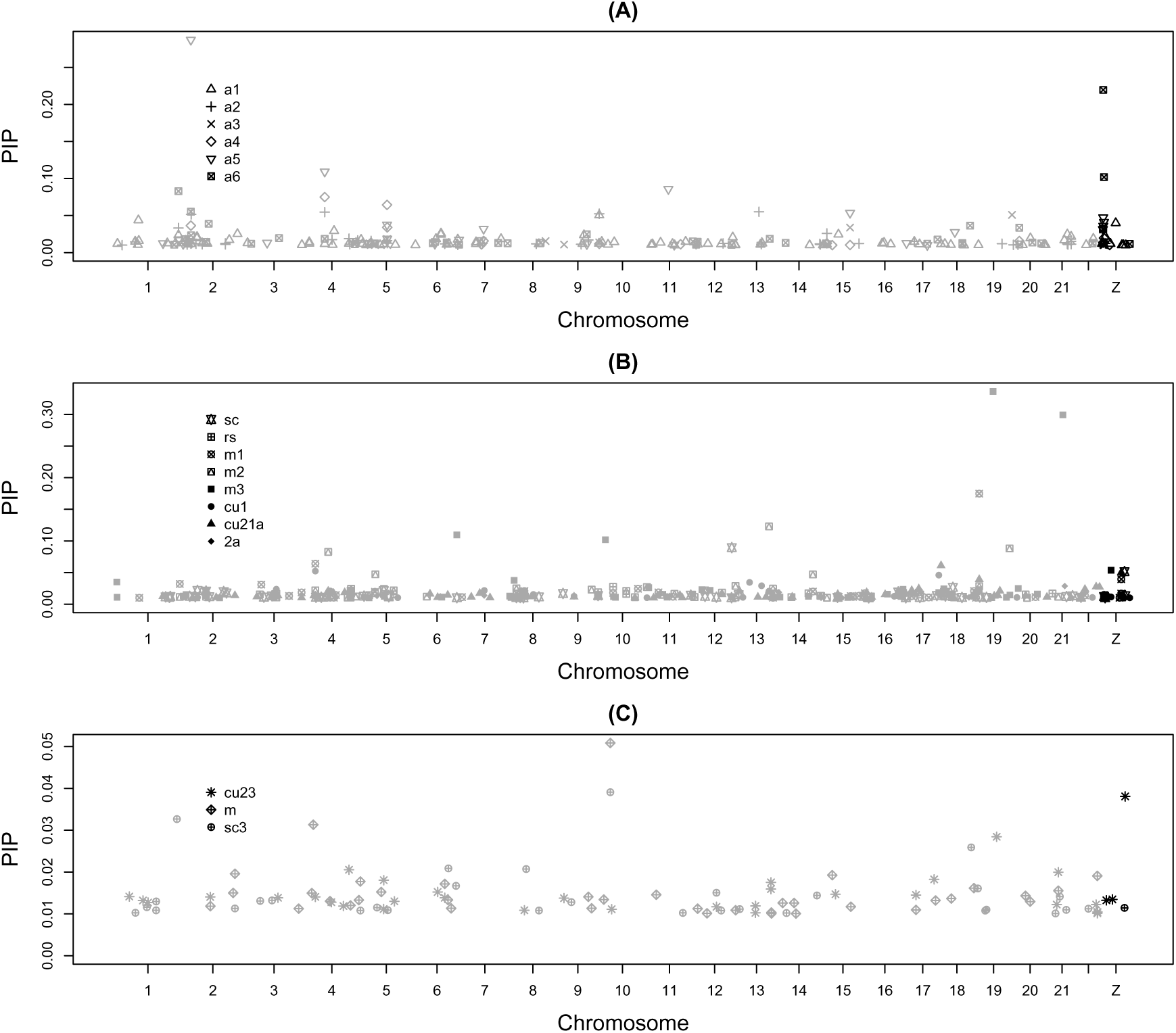
Manhattan plots of SNP posterior inclusion probabilities (PIPs) from polygenic association mapping of wing pattern traits in YG. Only SNPs with PIPs ≥ 0.01 are shown. Panels correspond to different subsets of wing pattern traits: (A) border eyespots (aurorae), (B) distal-center black dots, and (C) proximal-center black dots. SNPs on autosomes are shown in gray, whereas SNPs on the Z chromosome are shown in black.

## References

Abramoff M, Magelhaes P, Ram S (2004) Image processing with ImageJ. Biophotonics International, 11, 36–42.

Akopyan M, Tigano A, Jacobs A, Wilder AP, Baumann H, Therkildsen NO (2022) Comparative linkage mapping uncovers recombination suppression across massive chromosomal inversions associated with local adaptation in Atlantic silversides. Molecular Ecology, 31, 3323–3341.

Banker SE, Bonhomme F, Nachman MW (2022) Bidirectional introgression between *Mus musculus domesticus* and *Mus spretus*. Genome Biology and Evolution, 14, evab288.

Barroso G, Puzović N, Dutheil JY (2019) Inference of recombination maps from a single pair of genomes and its application to ancient samples. PLoS Genetics, 15, e1008449.

Barton N, Bengtsson B (1986) The barrier to genetic exchange between hybridizing populations. Heredity, 57, 357–376.

Bates D, Mächler M, Bolker B, Walker S (2015) Fitting linear mixed-effects models using lme4. Journal of Statistical Software, 67, 1–48.

Betancourt M (2017) A conceptual introduction to Hamiltonian Monte Carlo. *arXiv preprint arXiv:1701.02434*.

Bolte CE, Phannareth T, Zavala-Paez M, et al. (2024) Genomic insights into hybrid zone formation: The role of climate, landscape, and demography in the emergence of a novel hybrid lineage. Molecular Ecology, 33, e17430.

Boman J, Nolen ZJ, Backström N (2025) On the origin of an insular hybrid butterfly lineage. Evolution, 79, 510–524.

Bouckaert R, Vaughan TG, Barido-Sottani J, et al. (2019) BEAST 2.5: An advanced software platform for Bayesian evolutionary analysis. PLoS Computational Biology, 15, e1006650.

Brandvain Y, Kenney AM, Flagel L, Coop G, Sweigart AL (2014) Speciation and introgression between *Mimulus nasutus* and *Mimulus guttatus*. PLoS genetics, 10, e1004410.

Carling MD, Brumfield RT (2008) Haldane’s rule in an avian system: using cline theory and divergence population genetics to test for differential introgression of mitochondrial, autosomal, and sex-linked loci across the Passerina Bunting hybrid zone. Evolution, 62, 2600–2615.

Chaturvedi S, Lucas LK, Buerkle CA, et al. (2020) Recent hybrids recapitulate ancient hybrid outcomes. Nature Communications, 11, 1–15.

Chen Y, Chen Y, Shi C, et al. (2018) SOAPnuke: a MapReduce acceleration-supported software for integrated quality control and preprocessing of high-throughput sequencing data. Gigascience, 7, gix120.

Coyne JA, Orr HA (1989) Patterns of speciation in *Drosophila*. Evolution, 43, 362–381.

Degnan JH, Rosenberg NA (2009) Gene tree discordance, phylogenetic inference and the multispecies coalescent. Trends in Ecology & Evolution, 24, 332–340.

Dobzhansky TH (1937) Genetics and the Origin of Species. Columbia University Press, New York.

Dod B, Jermiin LS, Boursot P, Chapman VH, Nielsen JT, Bonhomme F (1993) Counter-selection on sex chromosomes in the *Mus musculus* European hybrid zone. Journal of Evolutionary Biology, 6, 529–546.

Drummond AJ, Rambaut A (2007) BEAST: Bayesian evolutionary analysis by sampling trees. BMC Evolutionary Biology, 7, 214.

Drummond AJ, Suchard MA (2010) Bayesian random local clocks, or one rate to rule them all. BMC Biology, 8, 1–12.

Durand EY, Patterson N, Reich D, Slatkin M (2011) Testing for ancient admixture between closely related populations. Molecular Biology and Evolution, 28, 2239–2252.

Ebbert MT, Wadsworth ME, Staley LA, et al. (2016) Evaluating the necessity of PCR duplicate removal from next-generation sequencing data and a comparison of approaches. BMC Bioinformatics, 17, 239.

Edelman NB, Mallet J (2021) Prevalence and adaptive impact of introgression. Annual review of genetics, 55, 265–283.

Felsenstein J (1981) Skepticism towards Santa Rosalia, or why are there so few kinds of animals? Evolution, 35, 124–138.

Flaxman SM, Feder JL, Nosil P (2013) Genetic hitchhiking and the dynamic buildup of genomic divergence during speciation with gene flow. Evolution, 67, 2577–2591.

Fordyce JA, Nice CC, Forister ML, Shapiro AM (2002) The significance of wing pattern diversity in the Lycaenidae: mate discrimination by two recently diverged species. Journal of Evolutionary Biology, 15, 871–879.

Forister M, Philbin C, Marion Z, et al. (2020) Predicting patch occupancy reveals the complexity of host range expansion. Science Advances, 6, eabc6852.

Forister ML, Fordyce JA, Nice CC, Gompert Z, Shapiro AM (2006) Egg morphology varies among populations and habitats along a suture zone in the *Lycaeides idas-melissa* species complex (Lepidoptera: Lycaenidae). Annals of the Entomological Society of America, 99, 933–937.

Forister ML, Gompert Z, Fordyce JA, Nice CC (2011) After 60 years, an answer to the question: what is the Karner blue butterfly. Biology Letters, 7, 399–402.

Forister ML, Nice CC, Fordyce JA, Gompert Z (2009) Host range evolution is not driven by the optimization of larval performance: the case of *Lycaeides melissa* (Lepidoptera: Lycaenidae) and the colonization of alfalfa. Oecologia, 160, 551–561.

Gompert Z, Brady M, Chalyavi F, et al. (2019) Genomic evidence of genetic variation with pleiotropic effects on caterpillar fitness and plant traits in a model legume. Molecular Ecology, 28, 2967–2985.

Gompert Z, Feder JL, Parchman TL, Planidin NP, Whiting FJ, Nosil P (2025) Adaptation repeatedly uses complex structural genomic variation. Science, 388, eadp3745.

Gompert Z, Lucas LK, Buerkle CA, Forister ML, Fordyce JA, Nice CC (2014) Admixture and the organization of genetic diversity in a butterfly species complex revealed through common and rare genetic variants. Molecular Ecology, 23, 4555–4573.

Gompert Z, Lucas LK, Fordyce JA, Forister ML, Nice CC (2010) Secondary contact between *Lycaeides idas* and *L. melissa* in the Rocky Mountains: extensive introgression and a patchy hybrid zone. Molecular Ecology, 19, 3171–3192.

Gompert Z, Lucas LK, Nice CC, Buerkle CA (2013a) Genome divergence and the genetic architecture of barriers to gene flow between *Lycaeides idas* and *L. melissa*. Evolution, 67, 2498–2514.

Gompert Z, Lucas LK, Nice CC, Fordyce JA, Buerkle CA, Forister ML (2013b) Geographically multifarious phenotypic divergence during speciation. Ecology and Evolution, 3, 595–613.

Gompert Z, Lucas LK, Nice CC, Fordyce JA, Forister ML, Buerkle CA (2012) Genomic regions with a history of divergent selection affect fitness of hybrids between two butterfly species. Evolution, 66, 2167–2181.

Gompert Z, Mandeville EG, Buerkle CA (2017) Analysis of population genomic data from hybrid zones. Annual Review of Ecology, Evolution, and Systematics, 48, 207–229.

Gompert Z, Nice CC, Fordyce JA, Forister ML, Shapiro AM (2006) Identifying units for conservation using molecular systematics: the cautionary tale of the Karner blue butterfly. Molecular Ecology, 15, 1759–1768.

Gompert Z, Saley T, Philbin C, et al. (2022) Additive genetic effects in interacting species jointly determine the outcome of caterpillar herbivory. Proceedings of the National Academy of Sciences, 119, e2206052119.

Gompert Z, Springer A, Brady M, Chaturvedi S, Lucas LK (2021) Genomic time-series data show that gene flow maintains high genetic diversity despite substantial genetic drift in a butterfly species. Molecular Ecology, 30, 4991–5008.

Goodwin KB, Chaturvedi S, Lucas LK, Gompert Z (2026) Genomic forecasts of maladptation in lycaeides butterflies. bioRxiv, pp. 2026–05.

Green RE, Krause J, Briggs AW, et al. (2010) A draft sequence of the Neandertal genome. Science, 328, 710–722.

Harrison RG, Larson EL (2014) Hybridization, introgression, and the nature of species boundaries. Journal of Heredity, 105, 795–809.

van der Heijden ES, Näsvall K, Seixas FA, et al. (2025) Genomics of neotropical biodiversity indicators: Two butterfly radiations with rampant chromosomal rearrangements and hybridization. Proceedings of the National Academy of Sciences, 122, e2410939122.

Heled J, Bouckaert RR (2013) Looking for trees in the forest: summary tree from posterior samples. BMC Evolutionary Biology, 13, 1–11.

Hooper DM, Griffith SC, Price TD (2019) Sex chromosome inversions enforce reproductive isolation across an avian hybrid zone. Molecular Ecology, 28, 1246–1262.

Hübner S, Bercovich N, Todesco M, et al. (2019) Sunflower pan-genome analysis shows that hybridization altered gene content and disease resistance. Nature Plants, 5, 54–62.

Iasi LN, Chintalapati M, Skov L, et al. (2024) Neanderthal ancestry through time: Insights from genomes of ancient and present-day humans. Science, 386, eadq3010.

Irwin DE (2018) Sex chromosomes and speciation in birds and other ZW systems. Molecular Ecology, 27, 3831–3851.

Jiggins CD, Salazar C, Linares M, Mavarez J (2008) Hybrid trait speciation and *Heliconius* butterflies. Philosophical Transactions of the Royal Society B-Biological Sciences, 363, 3047–3054.

Jin M, Peng Y, Peng J, et al. (2024) A supergene controls facultative diapause in the crop pest *Helicoverpa armigera*. Cell Reports, 43.

Kawahara AY, Storer C, Carvalho APS, et al. (2023) A global phylogeny of butterflies reveals their evolutionary history, ancestral hosts and biogeographic origins. Nature Ecology & Evolution, 7, 903–913.

Keightley PD, Pinharanda A, Ness RW, et al. (2015) Estimation of the spontaneous mutation rate in *Heliconius melpomene*. Molecular Biology and Evolution, 32, 239–243.

Kitano J, Ross JA, Mori S, et al. (2009) A role for a neo-sex chromosome in stickleback speciation. Nature, 461, 1079–1083.

Koch EL, Morales HE, Larsson J, et al. (2021) Genetic variation for adaptive traits is associated with polymorphic inversions in *Littorina saxatilis*. Evolution letters, 5, 196–213.

Langdon QK, Groh JS, Aguillon SM, et al. (2024) Swordtail fish hybrids reveal that genome evolution is surprisingly predictable after initial hybridization. PLoS Biology, 22, e3002742.

Le Corre V, Siol M, Vigouroux Y, Tenaillon MI, Délye C (2020) Adaptive introgression from maize has facilitated the establishment of teosinte as a noxious weed in Europe. Proceedings of the National Academy of Sciences, 117, 25618–25627.

Lenormand T, Roze D (2025) A single theory for the evolution of sex chromosomes and the two rules of speciation. Science, 389, eado9032.

Lewontin RC, Birch LC (1966) Hybridization as a source of variation for adaptation to new environments. Evolution, 20, 315–336.

Li H (2011) A statistical framework for SNP calling, mutation discovery, association mapping and population genetical parameter estimation from sequencing data. Bioinformatics, 27, 2987–2993.

Li H, Durbin R (2009) Fast and accurate short read alignment with burrows–wheeler transform. Bioinformatics, 25, 1754–1760.

Li H, Handsaker B, Wysoker A, et al. (2009) The Sequence Alignment/Map format and SAMtools. Bioinformatics, 25, 2078–2079.

Li L, Comi TJ, Bierman RF, Akey JM (2024) Recurrent gene flow between neanderthals and modern humans over the past 200,000 years. Science, 385, eadi1768.

Lipson M (2020) Applying f4-statistics and admixture graphs: Theory and examples. Molecular Ecology Resources, 20, 1658–1667.

Lopez H, Mercier R (2024) Meiosis: The silk moth and the elephant. Current Biology, 34, R211–R213.

Lowry DB, Willis JH (2010) A widespread chromosomal inversion polymorphism contributes to a major life-history transition, local adaptation, and reproductive isolation. PLoS Biology, 8, e1000500.

Lucas LK, Fordyce JA, Nice CC (2008) Patterns of genitalic morphology around suture zones in North American *Lycaeides* (Lepidoptera: Lycaenidae): implications for taxonomy and historical biogeography. Annals of the Entomological Society of America, 101, 172–180.

Lucas LK, Nice CC, Gompert Z (2018) Genetic constraints on wing pattern variation in *Lycaeides* butterflies: A case study on mapping complex, multifaceted traits in structured populations. Molecular Ecology Resources, 18, 892–907.

Lucas LK, Scholl CF, Jahner JJ, Fordyce JA, Nice CC, Forister ML (2026) Artificial hybridization of *Lycaeides* butterflies: asymmetrical effects and partial recovery of hybrid species phenotypes. Ecology and Evolution, p. Manuscript in preparation.

Mallet J (2005) Hybridization as an invasion of the genome. Trends in Ecology and Evolution, 20, 229–237.

Maroja LS, Larson EL, Bogdanowicz SM, Harrison RG (2015) Genes with restricted introgression in a field cricket (*Gryllus firmus*/*Gryllus pennsylvanicus*) hybrid zone are concentrated on the X chromosome and a single autosome. G3: Genes, Genomes, Genetics, 5, 2219–2227.

Marques DA, Meier JI, Seehausen O (2019) A combinatorial view on speciation and adaptive radiation. Trends in Ecology & Evolution, 34, 531–544.

Martin SH, Davey JW, Jiggins CD (2015) Evaluating the use of ABBA–BABA statistics to locate introgressed loci. Molecular Biology and Evolution, 32, 244–257.

Martin SH, Davey JW, Salazar C, Jiggins CD (2019) Recombination rate variation shapes barriers to introgression across butterfly genomes. PLoS biology, 17, e2006288.

Martin SH, Van Belleghem SM (2017) Exploring evolutionary relationships across the genome using topology weighting. Genetics, 206, 429–438.

Masly JP, Presgraves DC (2007) High-resolution genome-wide dissection of the two rules of speciation in drosophila. PLoS Biology, 5, e243.

Matute DR, Comeault AA, Earley E, et al. (2020) Rapid and predictable evolution of admixed populations between two *Drosophila* species pairs. Genetics, 214, 211–230.

Mavarez J, Salazar CA, Bermingham E, Salcedo C, Jiggins CD, Linares M (2006) Speciation by hybridization in *Heliconius* butterflies. Nature, 441, 868–871.

Mayr E (1963) Animal Species and Evolution. Harvard University Press, Cambridge, MA.

McKenna A, Hanna M, Banks E, et al. (2010) The Genome Analysis Toolkit: A MapReduce framework for analyzing next-generation DNA sequencing data. Genome Research, 20, 1297–1303.

Meier JI, McGee MD, Marques DA, et al. (2023) Cycles of fusion and fission enabled rapid parallel adaptive radiations in African cichlids. Science, 381, eade2833.

Müller NF, Bouckaert RR (2020) Adaptive Metropolis-coupled MCMC for BEAST 2. PeerJ, 8, e9473.

Nabokov V (1943) The nearctic forms of *Lycaeides* Hub. (Lycaenidae, Lepidoptera). Psyche, 50, 87–99.

Nabokov V (1949) The nearctic members of *Lycaeides* Hübner (Lycaenidae, Lepidoptera). Bulletin of the Museum of Comparative Zoology, 101, 479–541.

Neal RM (2011) MCMC using Hamiltonian dynamics. Handbook of Markov chain Monte Carlo, pp. 47–95.

Nice CC, Anthony N, Gelembiuk G, Raterman D, ffrench Constant R (2005) The history and geography of diversification within the butterfly genus *Lycaeides* in North America. Molecular Ecology, 14, 1741–1754.

Nice CC, Fordyce JA, Shapiro AM, ffrench Constant R (2002) Lack of evidence for reproductive isolation among ecologically specialised lycaenid butterflies. Ecological Entomology, 27, 702–712.

Nice CC, Gompert Z, Fordyce JA, Forister ML, Lucas LK, Buerkle CA (2013) Hybrid speciation and independent evolution in lineages of alpine butterflies. Evolution, 67, 1055–1068.

Nice CC, Shapiro AM (1999) Molecular and morphological divergence in the butterfly genus *Lycaeides* (Lepidoptera: Lycaenidae) in North America: evidence of recent speciation. Journal of Evolutionary Biology, 12, 936–950.

Nikolakis ZL, Schield DR, Westfall AK, et al. (2022) Evidence that genomic incompatibilities and other multilocus processes impact hybrid fitness in a rattlesnake hybrid zone. Evolution, 76, 2513–2530.

Nosil P, Soria-Carrasco V, Villoutreix R, et al. (2023) Complex evolutionary processes maintain an ancient chromosomal inversion. Proceedings of the National Academy of Sciences, 120, e2300673120.

Nouhaud P, Martin SH, Portinha B, Sousa VC, Kulmuni J (2022) Rapid and predictable genome evolution across three hybrid ant populations. PLoS Biology, 20, e3001914.

Orr HA (1997) Haldane’s rule. Annual Review of Ecology and Systematics, 28, 195–218.

Oziolor EM, Reid NM, Yair S, et al. (2019) Adaptive introgression enables evolutionary rescue from extreme environmental pollution. Science, 364, 455–457.

Payseur BA, Presgraves DC, Filatov DA (2018) Sex chromosomes and speciation. Molecular Ecology, 27, 3745.

Peñalba JV, Runemark A, Meier JI, et al. (2024) The role of hybridization in species formation and persistence. Cold Spring Harbor Perspectives in Biology, 16, a041445.

Poelstra JW, Vijay N, Bossu CM, et al. (2014) The genomic landscape underlying phenotypic integrity in the face of gene flow in crows. Science, 344, 1410–1414.

Presgraves DC (2008) Sex chromosomes and speciation in *Drosophila*. Trends in Genetics, 24, 336–343.

Rambaut A (2018) FigTree v1.4.4: A Graphical Viewer of Phylogenetic Trees. Computer program and documentation distributed by the author.

Rasmussen SW (1977) The transformation of the synaptonemal complex into the “elimination chromatin” in *Bombyx mori* oocytes. Chromosoma, 60, 205–221.

Rosenberg NA (2002) The probability of topological concordance of gene trees and species trees. Theoretical Population Biology, 61, 225–247.

Rosser N, Seixas F, Queste LM, et al. (2024) Hybrid speciation driven by multilocus introgression of ecological traits. Nature, 628, 811–817.

Rossi M, Hausmann AE, Alcami P, et al. (2024) Adaptive introgression of a visual preference gene. Science, 383, 1368–1373.

Sánchez-Ramírez S, Weiss JG, Thomas CG, Cutter AD (2021) Widespread misregulation of inter-species hybrid transcriptomes due to sex-specific and sex-chromosome regulatory evolution. PLoS Genetics, 17, e1009409.

Schield DR, Carter JK, Scordato ES, et al. (2024) Sexual selection promotes reproductive isolation in barn swallows. Science, 386, eadj8766.

Schumer M, Xu C, Powell DL, et al. (2018) Natural selection interacts with recombination to shape the evolution of hybrid genomes. Science, 360, 656–660.

Smith MR (2020a) Information theoretic generalized Robinson–Foulds metrics for comparing phylogenetic trees. Bioinformatics, 36, 5007–5013.

Smith MR (2020b) TreeDist: Distances between Phylogenetic Trees. R package version 2.11.0.

Soria-Carrasco V, Gompert Z, Comeault AA, et al. (2014) Stick insect genomes reveal natural selection’s role in parallel speciation. Science, 344, 738–742.

Sperling FA (1994) Sex-linked genes and species differences in Lepidoptera. The Canadian Entomologist, 126, 807–818.

Springer A, Kissmer B, Gompert Z (2025) Admixture affects the rate and repeatability of experimental adaptation to a stressful environment in *Callosobruchus maculatus*. Molecular Ecology, p. e17809.

Stan Development Team (2019) RStan: the R interface to Stan. R package version 2.19.2.

Storchová R, Reif J, Nachman MW (2010) Female heterogamety and speciation: reduced introgression of the Z chromosome between two species of nightingales. Evolution, 64, 456–471.

Suvorov A, Kim BY, Wang J, et al. (2022) Widespread introgression across a phylogeny of 155 *Drosophila* genomes. Current Biology, 32, 111–123.

Swaegers J, Sánchez-Guillén RA, Chauhan P, Wellenreuther M, Hansson B (2022) Restricted X chromosome introgression and support for Haldane’s rule in hybridizing damselflies. Proceedings of the Royal Society B, 289, 20220968.

Talavera G, Lukhtanov VA, Pierce NE, Vila R (2013) Establishing criteria for higher-level classification using molecular data: the systematics of *Polyommatus* blue butterflies (Lepidoptera, Lycaenidae). Cladistics, 29, 166–192.

Taylor SA, Larson EL (2019) Insights from genomes into the evolutionary importance and prevalence of hybridization in nature. Nature Ecology & Evolution, 3, 170–177.

Thawornwattana Y, Seixas F, Yang Z, Mallet J (2023) Major patterns in the introgression history of *Heliconius* butterflies. eLife, 12, RP90656.

Todesco M, Owens GL, Bercovich N, et al. (2020) Massive haplotypes underlie ecotypic differentiation in sunflowers. Nature, 584, 602–607.

Toro-Delgado E, Gauthier J, Hinojosa JC, et al. (2025) Genomic analysis of *Plebejus* Kluk (Lycaenidae: Polyommatinae) clarifies taxonomy within Europe. Systematic Entomology.

Vasimuddin M, Misra S, Li H, Aluru S (2019) Efficient architecture-aware acceleration of BWA-MEM for multicore systems. In: 2019 IEEE international parallel and distributed processing symposium (IPDPS), pp. 314–324, IEEE.

Vehtari A, Gabry J, Magnusson M, et al. (2024) loo: Efficient leave-one-out cross-validation and waic for Bayesian models. R package version 2.8.0.

Vehtari A, Gelman A, Gabry J (2017) Practical Bayesian model evaluation using leave-one-out cross-validation and WAIC. Statistics and Computing, 27, 1413–1432.

Vila R, Bell CD, Macniven R, et al. (2011) Phylogeny and palaeoecology of *Polyommatus* blue butterflies show Beringia was a climate-regulated gateway to the New World. Proceedings of the Royal Society B: Biological Sciences, 278, 2737–2744.

Westram AM, Stankowski S, Surendranadh P, Barton N (2022) What is reproductive isolation? Journal of Evolutionary Biology, 35, 1143–1164.

Wu CI (2001) The genic view of the process of speciation. Journal of Evolutionary Biology, 14, 851–865.

Zhang C, Nielsen R, Mirarab S (2025) CASTER: Direct species tree inference from whole-genome alignments. Science, 387, eadk9688.

Zhang L, Chaturvedi S, Nice CC, Lucas LK, Gompert Z (2023) Population genomic evidence of selection on structural variants in a natural hybrid zone. Molecular Ecology, 32, 1497–1514.

Zhou X, Carbonetto P, Stephens M (2013) Polygenic modeling with Bayesian sparse linear mixed models. PLoS Genetics, 9, e1003264.

Zhou X, Stephens M (2012) Genome-wide efficient mixed-model analysis for association studies. Nature Genetics, 44, 821–824.

